# Human SHED-derived extracellular cues activate a specialized neuroprotective and regenerative program in developing retinal ganglion cells

**DOI:** 10.64898/2026.06.25.733625

**Authors:** Marian Mellen, Gema Garcia-Guirado, Laura Botana, Enrique Calvo, Yariliz Sencion, Mattia Biondo, Javier Díez-Mata, Jesus Vazquez, Ismael Santa-Maria, Maite Iglesias

## Abstract

Axonal degeneration and insufficient neuronal survival remain major barriers to central nervous system repair. Stem cells from human exfoliated deciduous teeth (SHED) represent an accessible, developmentally immature, neural crest-derived mesenchymal stem cell population with potential relevance for neuroregenerative medicine. Here, we show that SHED display enhanced proliferative stability, preserved mesenchymal identity, and more sustained expansion capacity than adult dental pulp stem cells, supporting their suitability for scalable regenerative applications. Using embryonic chick retinal explants at neurogenic and post-neurogenic stages, we demonstrate that SHED robustly promote retinal ganglion cell axonogenesis, axonal regeneration, and neuronal survival. At embryonic day 5, SHED enhanced axonal outgrowth in both newly generated EdU⁺/TUJ1⁺ neurons and pre-existing EdU⁻/TUJ1⁺ retinal ganglion cells. At embryonic day 13, when retinal neurons are post-mitotic and intrinsically less regenerative, SHED still significantly increased regenerative axonal extension and reduced developmental cell death. To investigate the molecular mechanisms underlying the neuroprotective and axogenic effects of SHED, proteomic profiling of SHED-retina co-culture secretomes was performed, revealing a highly enriched extracellular environment containing matrix-associated and neurodevelopmental proteins, including thrombospondin-1 (THBS1), galectin1 and 3, and multiple proteins associated with IGF2 pathway. Proteomic analysis of the SHED secretome, together with prior evidence implicating thrombospondin signaling in neuronal development and synaptogenesis, identified THBS1 as a strong candidate mediator of SHED-induced effects in chick retinal co-culture systems. Neutralization of THBS1, particularly in combination with gabapentin-mediated blockade of α2δ-1-dependent thrombospondin signaling, markedly reduced SHED-induced axonal growth and induced neuritic swellings consistent with impaired axonal integrity. In contrast, inhibition of THBS1 signaling did not significantly abolish the neuroprotective effect of SHED on neuronal survival, suggesting that distinct paracrine mechanisms independently regulate axonal regeneration and cell survival. Together, these findings demonstrate that SHED-derived combined secreted factors promote neuronal survival and axonal regeneration through partially divergent extracellular matrix-associated developmental pathways, positioning SHED and their secretome as promising candidates for cell-based and cell-free neuroregenerative strategies.

## INTRODUCTION

The vertebrate retina provides a well-established model to study neuronal development and regeneration. In particular, the embryonic chick retina offers a unique opportunity to temporally decouple retinal ganglion cells (RGC) neurogenesis from axonal regeneration. RGC are among the first neurons generated during retinogenesis, with chick RGC differentiation beginning as early as embryonic day 2 and peaking around E5–E6 (1). By mid-incubation (E10-E11), chick RGC have extended axons to their central target (optic tectum) and completed much of their developmental maturation. Notably, young chick RGC possess a robust intrinsic capacity for axon outgrowth and regeneration that is progressively lost as the retina matures (1). For example, explant studies have shown that RGC from early embryos extend long neurites rapidly, whereas RGC from later embryonic stages (beyond E13–E14) display markedly slower and limited axon regrowth. Thus, by choosing specific developmental time points in the chick (e.g., E5 vs. E13), one can separate the phase of new RGC neurogenesis/axonogenesis (largely accomplished by E5–E7) from the phase of axonal regeneration in a more mature retinal environment (2). This experimental paradigm allows investigation of how pro-regenerative interventions affect immature versus mature RGC in the same organism.

During normal chick retinal development, a significant fraction of RGC undergo programmed cell death, which is a critical process for refining neuronal populations. Two distinct waves of apoptosis have been documented: an early wave coinciding with the onset of RGC neurogenesis (around E5–E7), and a later wave after RGC axons have innervated the tectum (approximately E10–E14) (2). This later phase corresponds to the well-known period of target-dependent RGC death, wherein excess neurons are eliminated after reaching the brain target. Importantly, such developmental neuron death is not an autonomous process; it is profoundly influenced by extracellular survival signals in the retinal environment. Classic neurotrophic factors can modulate RGC survival during these critical periods. For instance, in the chick embryo, exogenous brain-derived neurotrophic factor (BDNF) administration during early retinal development dramatically rescues neurons from apoptosis, yielding ∼70% more RGC than normal by E6–E9 (2). Conversely, certain endogenous cues like nerve growth factor (NGF) acting through the p75 neurotrophin receptor can actively induce apoptosis in early retinal progenitors (3). These observations underscore that programmed RGC death is regulated by extracellular signals, and that providing pro-survival factors can enhance retinal neuron retention. Such developmental principles set the stage for exploring therapeutic strategies to prevent neuron loss or boost regeneration by mimicking pro-survival signals.

In the context of neural repair, mesenchymal stem cells (MSC) have emerged as a promising source of neurotrophic support. Recently, stem cells from human exfoliated deciduous teeth (SHED) have gained attention as a unique MSC population with remarkable neuro-regenerative potential (4). SHED are isolated from the pulp of naturally shed baby teeth and are considered an immature, neural-crest-derived MSC source. They exhibit high proliferative capacity and multilineage differentiation ability, having been shown to differentiate into osteogenic, chondrogenic, adipogenic, and even neural-like cells *in vitro* and *in vivo* (5). In contrast, dental pulp stem cells (DPSC), derived from permanent teeth, represent a more mature dental MSC population and provide an important comparative model to evaluate the unique biological and neuroregenerative properties of SHED. Furthermore, SHED cells are easily accessible with minimal invasiveness and display low immunogenicity, making them attractive for transplantation purposes. Of relevance to neurobiology, SHED secrete a rich repertoire of growth factors and cytokines, a neurotrophic secretome, that can support neuronal survival and outgrowth (6). Indeed, previous studies have demonstrated that SHED release numerous trophic factors (e.g., BDNF, NGF, IGF-1, HGF) and anti-inflammatory cytokines, which collectively help create a pro-survival microenvironment. In various CNS injury models, these cells have shown the ability to promote neural repair: for example, conditioned media from SHED cultures can protect motor neurons and reduce toxicity in vitro, and SHED transplantation improves functional recovery in rodent models of neural injury (7).

The mechanisms by which SHED exert their beneficial effects appear to involve both direct cellular interactions and paracrine signaling. On one hand, SHED can directly engage with neural tissue – for instance, they can migrate, differentiate, or form cell–cell contacts that replace or support damaged cells (7). Transplantation studies in spinal cord injury have reported that engrafted SHED not only survive but can differentiate into mature glial cells (such as oligodendrocytes) at the injury site, thereby contributing to tissue repair by cell replacement. On the other hand, a wealth of evidence indicates that paracrine effects are a dominant mode of action: SHED secrete soluble factors that modulate the extracellular milieu and activate endogenous cells’ regenerative programs (5). Notably, SHED-derived conditioned medium itself confers neuroprotection and axon growth promotion in the absence of the cells, underscoring the role of secreted trophic molecules (6). In fact, teeth-derived stem cells have been shown to promote axon growth and inhibit apoptosis in injured CNS tissue largely via secreted factors, while also providing metabolic support and immunomodulation (8). Both cell-autonomous (direct) and paracrine mechanisms are likely to work in concert in SHED-mediated neuroprotection. This dual mode of action is highly relevant for developmental and regenerative contexts: SHED could directly integrate into developing retinal tissue or, more feasibly, secrete factors that reactivate growth pathways in host neurons. However, the specific influence of SHED on retinal neurons, especially across different developmental stages, remains to be fully elucidated.

Based on these considerations, we hypothesize that SHED-derived signals can promote axonogenesis and neuroprotection in retinal ganglion cells at both immature and more mature stages by activating conserved developmental programs. In an early developmental context (such as the embryonic chick at E5, when RGC are still being generated and extending nascent axons), we propose that SHED-secreted factors will stimulate axon outgrowth and enhance neuronal survival, effectively boosting the intrinsic growth state of immature RGC. Likewise, in a later developmental stage (such as E13, when RGC are differentiated and normally exhibit limited regenerative capacity), we expect that SHED-derived cues can re-activate embryonic growth pathways and protect mature RGC from cell death while encouraging regenerative axon sprouting. This hypothesis is grounded in the idea that the pro-growth and pro-survival signals from SHED mimic those present during early development, thereby conferring an ability to recapitulate early developmental programs even in older neurons.

To test this hypothesis, the present study investigates the effects of SHED on RGC axon growth and survival in the developing chick retina at two distinct embryonic stages (E5 and E13). We set out to dissect how SHED-mediated signals influence neurons that are either immature (actively undergoing neurogenesis at E5) or relatively mature (post-mitotic and projecting axons at E13). By using the chick retina model, we examine whether SHED co-culture or SHED-conditioned medium can promote axonal extension/regeneration in vitro and enhance RGC survival (mitigating developmental cell death) at each stage. Thus, elucidating SHED’s neuroprotective and axonogenic mechanisms supports the development of stem cell-based strategies in retinal neuroprotection and regeneration.

## MATERIALS AND METHODS

### SHED isolation and cell culture

Human exfoliated deciduous teeth were collected from healthy pediatric donors (ages 7–10) following informed consent and institutional ethical approval. Human SHED were successfully isolated from exfoliated deciduous teeth using an explant culture approach following procedures previously described (4). Tissue fragments from the pulp of naturally exfoliated teeth were plated in tissue culture dishes with Dulbecco’s modified Eagle’s medium/F12 (DMEM/F12) (Gibco, cat #11320033) supplemented with 10% fetal bovine serum (FBS; Gibco), 100 U/mL penicillin, and 100 μg/mL streptomycin. Cells began migrating out from the explants within 48–72 hours. By day 5, spindle-shaped fibroblast-like cells with bipolar extensions emerged in high density, showing a morphology consistent with mesenchymal stem cells (MSC). Cultures were maintained at 37°C in a humidified incubator with 5% CO₂. Migrating cells were passaged upon confluence using 0.05% trypsin-EDTA and expanded up to passage 5. Four independent donor-derived SHED lines were used. Dental Pulp Stem Cells (DPSC) (Lonza) and HaCaT keratinocytes (Fisher Scientific) were also cultured with DMEM/F12 medium (Gibco) supplemented with 10% fetal bovine serum (FBS; Gibco), 100 U/mL penicillin, and 100 μg/mL streptomycin. HaCaT keratinocytes were used as negative controls.

### SHED characterization

SHED were phenotyped by immunocytochemistry using anti-vimentin (Abcam # ab137321, 1:500), anti-CD105 (ABclonal cat #4000000446, 1:200), and anti-STRO-1 (Novus Biological cat #NBP1-48356, 1:1000) antibodies. Multilineage differentiation was performed using commercial osteogenic, adipogenic, and chondrogenic induction media (Gibco). Alizarin Red S, Oil Red O, and Alcian Blue staining were used to confirm lineage-specific differentiation. Cell morphology was imaged with phase-contrast microscopy (DMi8 THUNDER, Leica, Wetzlar, Germany).

### Flow cytometry analysis

To confirm MSCs phenotype cells were analyzed by flow cytometry. Antibodies against the following human antigens were used: CD105-FITC (Miltenyi Biotect, Bergisch Gladbach, Germany, cat# 130-112-327, 1:50), CD90-FITC (Miltenyi Biotec, cat# 130-114-901, 1:50), CD44-VioBlue (Miltenyi Biotec, cat# 130-113-906, 1:50), CD73-APC (Miltenyi Biotec, cat# 130-111-909, 1:50), MSC Phenotyping Cocktail-PE (CD34, CD14, CD19, CD45, Miltenyi Biotec cat# 130-125-285, dilution according to the manufacturer’s instructions). Briefly, cells were seeded at a concentration of 104 cells/cm2 and maintained in culture until they reached 80%–90% confluence. Then cells were trypsinized, surface labeled, washed and then analyzed using a flow cytometer (Miltenyi Biotech). The data were analyzed with MACSQuantify™ Software.

### Proliferation capacity of SHED and DPSC

To examine the proliferation capacity of the SHED and DPSC *in vitro*, the cumulative population doubling (PD) was calculated over 30 days (from passage 3–8). Cells were placed in triplicate into 6 multiwell cell culture plates at a concentration of 10^4^ cells/cm2 and subcultured after 5 days at the same density. The cells were counted using a hemocytometer. The cumulative cell doubling of the cell populations was plotted against time in the culture to determine the growth kinetics of SHED and DPSC expansion. The number of population doubling was determined by counting the number of adherent cells at the start and end of each passage. The population doubling was calculated at every passage according to the equation: log_2_ (number of harvested cells/ number of seeded cells). The finite population doubling was determined by the cumulative addition of the total numbers generated from each passage until the cells stopped dividing.

### Retinal explant culture and SHED co-culture

Fertilized White Leghorn chicken eggs were incubated at 37.5°C in a humidified incubator with constant rotation every 2 hours. Embryonic day 5 (E5) and day 13 (E13) retinas were dissected in cold PBS and dissociated with Worthington’s Papain System (20 units/mL; Worthington-Biochemical Corporation, Lakewood, NJ, cat# LK003150). Dissociated chicken retinas were plated onto SHED (3 x 10^4^ or 5 x 10^4^) and HaCaT (5 x 10^4^) monolayers in 24 multiwell cell culture plates and the co-cultures were maintained for 72 hours in DMEM/F12, supplemented with 10 μM of EdU from (EdU Staining Protocol Kit at# ab219801). Dissociated chicken retinal cells were also plated in 2D cultures as negative control. Co-cultures were fixed with 4%paraformaldehyde (PFA).

To evaluate the involvement of thrombospondin-1 (THBS1) signaling in the neuroprotective effects exerted by SHED, inhibition assays were performed using gabapentin (GBP, Merck cat #10900010) and an anti-THBS1 blocking antibody (Invitrogen cat #14-9756-82). SHED were seeded in 24 multiwell cell culture plates at a density of 5 x 10⁴ cells per well in DMEM/F12 medium. The anti-THBS1 antibody was prepared from a 0.5 mg/mL stock solution and used at a final concentration of 10 μg/mL. Gabapentin was prepared from a 100 mM stock solution and used at a final concentration of 32 μM. An isotype-matched IgG antibody (Invitrogen cat #16-4714-82) was included as a control at a final concentration of 10 μg/mL. The experimental groups consisted of: (i) RGC cultured alone (control), (ii) RGC directly co-cultured with SHED, (iii) RGC co-cultured with SHED in the presence of IgG, (iv) RGC co-cultured with SHED in the presence of the anti-THBS1 antibody, (v) RGC co-cultured with SHED in the presence of gabapentin, and (vi) RGC co-cultured with SHED in the presence of both the anti-THBS1 antibody and gabapentin. Inhibitors and control antibodies were added at the onset of co-culture and maintained throughout the experiment. Cultures were incubated for 72 h prior to analysis.

Each experimental condition was established from three to six biological replicates.

### Immunofluorescence and imaging

Explants were permeabilized with 0.5% Triton X-100 and blocked in 10% BSA. Primary antibodies used were anti-TUJ1 (Abcam cat #ab78078, 1:500), anti-PAX6 (ABclonal cat #5500010825 1:200). Secondary antibodies conjugated to Alexa Fluor dyes (Thermo Fisher) were used at 1:1000. EdU incorporation was detected using the Click-iT EdU Imaging Kit (Thermo Fisher). TUNEL labeling was performed using the In Situ Cell Death Detection Kit (Roche cat #11684795910). For sectioned retinas immunohistochemistry, E5 heads were fixed in Histofix (4% formaldehyde), dehydrated through a graded ethanol series (96% and 100%), cleared in xylene, and embedded in paraffin. Paraffin blocks were sectioned at 3 µm thickness using a microtome (Thermo Scientific HM 325), and sections were mounted onto treated glass slides. For immunofluorescence analysis, sections were deparaffinized, rehydrated, and subjected to heat-induced antigen retrieval using citrate buffer. Non-specific binding sites were blocked prior to overnight incubation at 4°C with the primary antibodies anti-PAX6 and anti-TUJ1. After washing, sections were incubated with the appropriate fluorophore-conjugated secondary antibodies. Nuclei were counterstained with DAPI contained in the mounting medium. The samples were observed and images acquired in an inverted fluorescent microscope (DMi8 THUNDER, Leica, Wetzlar, Germany). Images were processed using Fiji (ImageJ).

### Quantification and statistical analysis

Axon outgrowth was quantified by measuring the average neurite length per TUJ1+ neuron using NeuronJ. Apoptosis was quantified as the percentage of TUNEL+ nuclei over total DAPI+ nuclei in defined regions of interest. At least five independent biological replicates and three technical replicates were used per condition. Data was analyzed using GraphPad Prism 11. Statistical comparisons were made using one-way ANOVA with Tukey’s post hoc test or two-tailed Student’s t-test where appropriate. A p-value < 0.05 was considered significant.

### Sample preparation for proteomics analysis

Three independent SHED and SHED co-cultured with dissociated E5 chicken retinas secretomes were collected in serum-free medium for 48 h (Mesenpro, Gibco), concentrated using Amicon Ultra-15 centrifugal filters (3 kDa cutoff). Around 3 ml of secretome were subjected to in-filter reduction and alkylation using iodoacetamide followed by overnight trypsin digestion (Nanosep Centrifugal Devices with Omega Membrane-10K, PALL), and the resulting tryptic peptides were desalted with Oasis HLB cartridges (Waters), and TMT-labelled following the instructions of the manufacturer (Thermo Fisher). A portion of the labelled-peptide mixtures was used for whole proteome analysis after high pH reversed phase fractionation (Pierce) into 5 peptide fractions containing 5%, 12.5%, 17.5%, 22.5% and 50% acetonitrile, respectively.

### Liquid chromatography-tandem mass spectrometry (LC-MS/MS)

Labeled peptides were loaded and washed on Evotips for chromatographic separation in an evosep one HPLC system (30 SPD method, with Endurance Column 15 cm x 150 µm ID, 1.9 µm beads-EV1106, Evosep) coupled to a stainless steel emitter of 30 μm ID. Eluted peptides were subjected to real-time ionization and MS analysis in an Orbitrap Eclipse Tribrid mass spectrometer (Thermo Fisher, San José, CA, USA) with a 2 seconds-TopSpeed method. MS spectra were acquired in the Orbitrap analyser with a 390-1700 m/z range and 60,000 FT resolution. Higher-energy collision dissociation (HCD) fragmentation was performed at 33 eV of normalized collision energy and MS/MS spectra were analyzed at a resolution of 30,000 in the Orbitrap. Dynamic exclusion was set to 20 s.

### Protein identification and quantification

Protein identification was performed using the SEQUEST HT algorithm integrated in Proteome Discoverer 2.5 (Thermo Scientific). MS/MS scans were searched against a human reference proteome database (human_202603_pro-sw.target-decoy.fasta). For database searching, parameters were selected as follows: trypsin digestion with 2 maximum missed cleavage sites, precursor mass tolerance of 2 Da, fragment mass tolerance of 0.03 Da. Methionine oxidation (+15.994915 Da) and asparagine and glutamine deamidation (+0.984016 Da) were set as variable modifications, while cysteine carbamidomethylation (+57.021464 Da) and TMT labeling at peptide N-terminal end and Lys were considered as fixed modifications. False discovery rates (FDR) of peptide identifications were calculated using the refined method (9) with an additional filter for precursor mass tolerance of 10 ppm (10). 1%FDR was used as criterion for peptide identification. Protein quantification analysis was performed following established protocols (9,11) with the SanXoT software package (12). Quantitative information was extracted from the MS/MS spectra of TMT-labeled peptides. Peptide and protein quantification were analyzed using the WSPP model, which uses raw quantifications as input data and computes the log_2_-fold changes for each reporter with respect to the reference internal standard sample. In this model protein log_2_-ratios are expressed as standardized variables in units of standard deviation according to their estimated variances (Zq values). Functional profiling was assessed with G-Profile(https://biit.cs.ut.ee/gprofiler/r) using Gene Ontology as data source, and Reactome (www.reactome.org) pathway browser. Vesiclepedia data of top 100 EV-associated proteins were downloaded from www.microvesicles.org.

## RESULTS

### Isolation, characterization, and comparative analysis of SHED demonstrate enhanced proliferative stability relative to DPSC

Human SHED were successfully isolated from exfoliated deciduous teeth and expanded using an explant-based culture approach. Cells migrated from dental pulp explants within 48–72 h and displayed a characteristic spindle-shaped fibroblast-like morphology consistent with mesenchymal stem cells. Comparative analyses between SHED and DPSC demonstrated that both populations retained mesenchymal characteristics but SHED exhibited superior proliferative stability and long-term expansion potential.

Phase-contrast microscopy revealed that SHED maintained a homogeneous fibroblastoid morphology throughout serial passaging, whereas DPSC cultures progressively displayed enlarged, flattened cells and features compatible with replicative senescence (Supplementary Figure 1a–d). Both populations retained multilineage differentiation potential, as demonstrated by osteogenic, adipogenic and chondrogenic differentiation assays (Supplementary Figure 1e–g).

Immunofluorescence analyses confirmed expression of Vimentin, CD105 and STRO-1 in SHED and DPSC cultures, whereas HaCaT epithelial cells lacked expression of these markers (Figure 1a-b). Flow cytometric characterization further confirmed expression of canonical MSC markers and absence of hematopoietic markers (Figure 1c). Importantly, cumulative population doubling analyses demonstrated that SHED maintained significantly greater proliferative capacity than DPSC across serial passages (Figure 1d), supporting their suitability for scalable regenerative applications.

**Figure 1.**
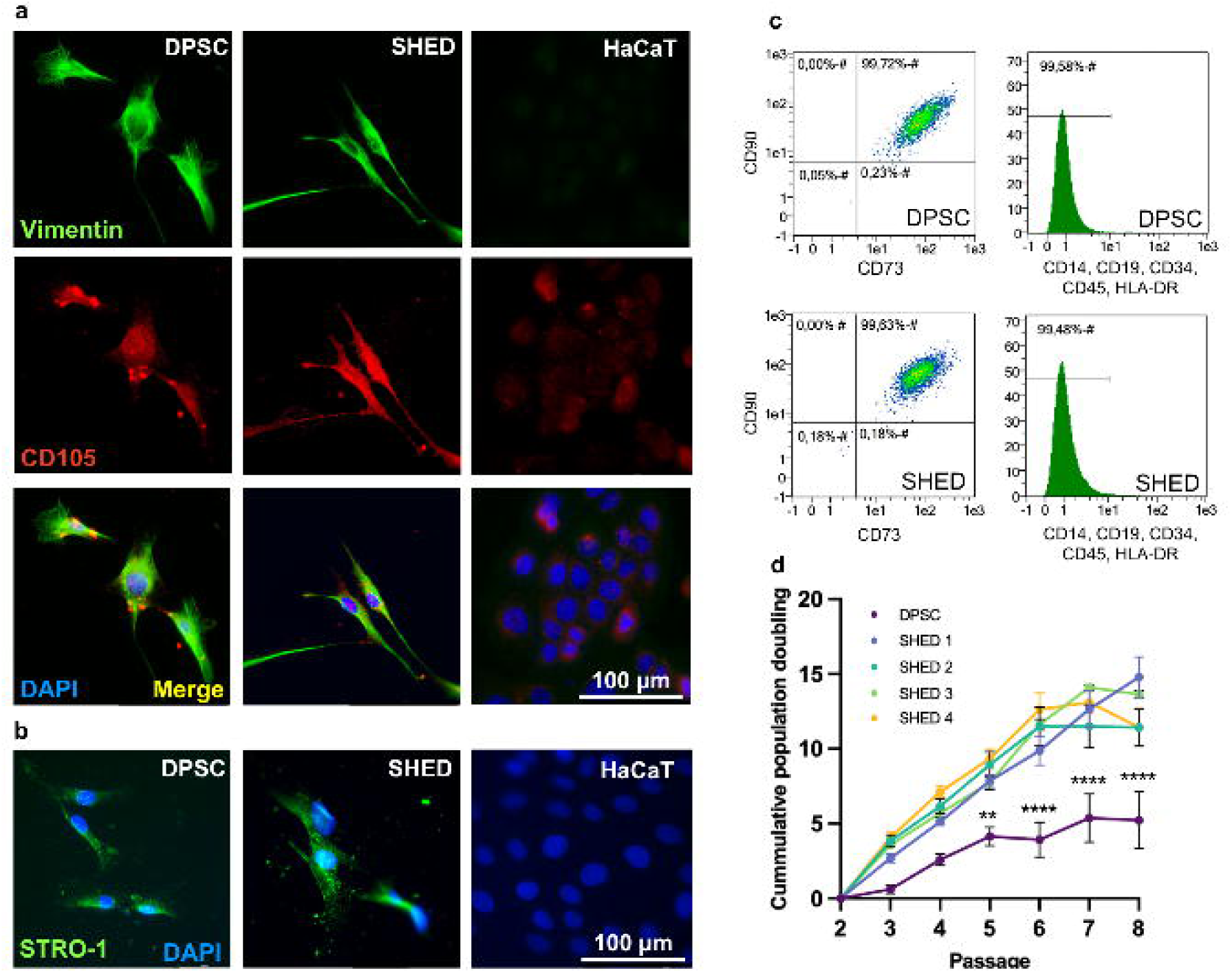
Immunophenotypic characterization and proliferative analysis of SHED. (a) Immunofluorescence staining showing expression of Vimentin (green) and CD105 (red) in DPSC and SHED compared with HaCaT epithelial controls. (b) STRO-1 (green) immunostaining in SHED. Nuclei were counterstained with DAPI (blue). Scale bar = 100μm (c) Representative flow cytometry analyses demonstrating expression of mesenchymal stem cell markers (CD73, CD90, CD105, HLA-ABC) and absence of hematopoietic markers (CD34, CD45). All experiments were perfomed at passage 5 (d) Growth curves comparing cumulative population doublings of DPSC and SHED derived from four independent donors across serial passages. SHED displayed enhanced proliferative capacity and reduced donor-to-donor variability.

### SHED promote axonogenesis and regenerative axon growth in developing and mature retinal ganglion cells

To determine whether SHED-derived extracellular signals influence neuronal growth, we established co-culture systems using embryonic chick retinal explants at two distinct developmental stages. E5 retinas represent an actively neurogenic environment containing retinal progenitors and newly generated retinal ganglion cells (RGC), whereas E13 retinas contain mature postmitotic neurons undergoing axotomy during explant preparation (Supplementary Figure 2).

At E5, TUJ1-positive RGC are specifically localized within the emerging ganglion cell layer (GCL) (Figure 2a). After retinal dissociation, EdU incorporation in culture allowed discrimination between newly generated TUJ1+/EdU+ neurons and pre-existing TUJ1+/EdU− RGC (Supplementary Figure 2 a). SHED treatment significantly increased axon length compared with control and HaCaT conditions (Figure 2a–c). Notably, SHED enhanced axonal extension in both newborn and mature neuronal populations, indicating activation of conserved growth programs independently of neuronal maturation status.

**Figure 2.**
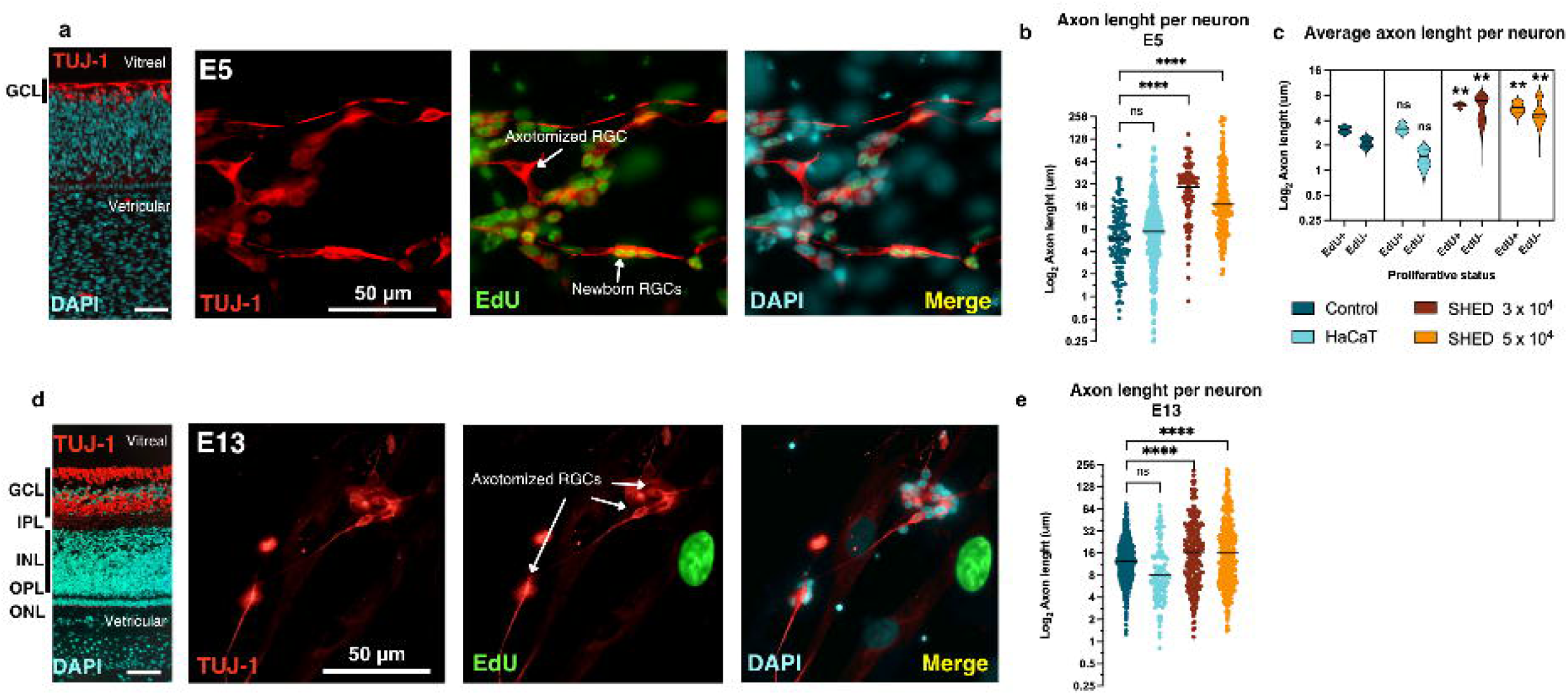
SHED promote axonal growth in both newly generated and axotomized retinal ganglion cells. (a) Representative section of an E5 chick retina stained for TUJ1 and DAPI, showing the presence of retinal ganglion cells (RGC) within the developing ganglion cell layer (GCL). Representative immunofluorescence images of E5 chick retinal co-cultures labeled with TUJ-1 (red), EdU (green), and DAPI (blue). EdU-positive/TUJ-1-positive cells correspond to newly generated retinal ganglion cells (RGC), whereas TUJ-1-positive/EdU-negative cells represent axotomized mature RGC. Scale bar = 50 μm. (b) Quantification of axon length per neuron in E5 RGC under control, HaCaT, and SHED co-culture conditions. SHED treatment significantly increased neurite extension compared with control conditions, whereas HaCaT cells had no significant effect. (c) Quantification of average axon length per neuron in EdU-positive newborn neurons and EdU-negative axotomized RGC under the indicated co-culture conditions. SHED significantly enhanced axonal regeneration relative to controls in both populations. (d) Representative section of an E13 chick retina stained for TUJ1 and DAPI, showing the presence of RGC within the GCL. Representative immunofluorescence images showing TUJ-1-positive axotomized RGC under the indicated conditions. (e) Quantification of axon length per neuron in E13 RGC under control, HaCaT, and SHED co-culture conditions. SHED treatment significantly increased neurite extension compared with control conditions, whereas HaCaT cells had no significant effect. Statistical analysis was performed using ANOVA followed by post hoc comparisons. Data are presented as mean ± SEM; n=4.

At E13, TUJ1-positive RGC are restricted to the fully developed GCL (Figure 2 d). Similarly, SHED robustly enhanced axonal regeneration in E13 retinal explants (Figure 2d–e). Because E13 neurons are postmitotic and intrinsically less regenerative, these findings indicate that SHED-derived signals reactivate developmental growth mechanisms in mature injured neurons. Additional representative images illustrating these effects are shown in Supplementary Figure 3.

### SHED exert potent neuroprotective effects during developmental and injury-associated cell death

As previously mentioned, programmed neuronal death represents a critical feature of chicken retinal development at these embyonic days (Supplementary Figure 2) and injury responses. TUNEL analyses revealed that SHED treatment markedly reduced apoptosis in both E5 and E13 retinal explants (Figure 3 a–c). At E5, SHED significantly reduced physiological developmental apoptosis occurring during early retinal neurogenesis. Similarly, at E13, SHED reduced neurotrophic and injury-associated cell death following axotomy. This effect was dose-dependent and absent in HaCaT controls, demonstrating specificity of the SHED-mediated response. Representative high-resolution images are provided in Supplementary Figure 4. Together, these results demonstrate that SHED simultaneously enhance neuronal survival and axonal growth, suggesting activation of coordinated regenerative programs.

**Figure 3.**
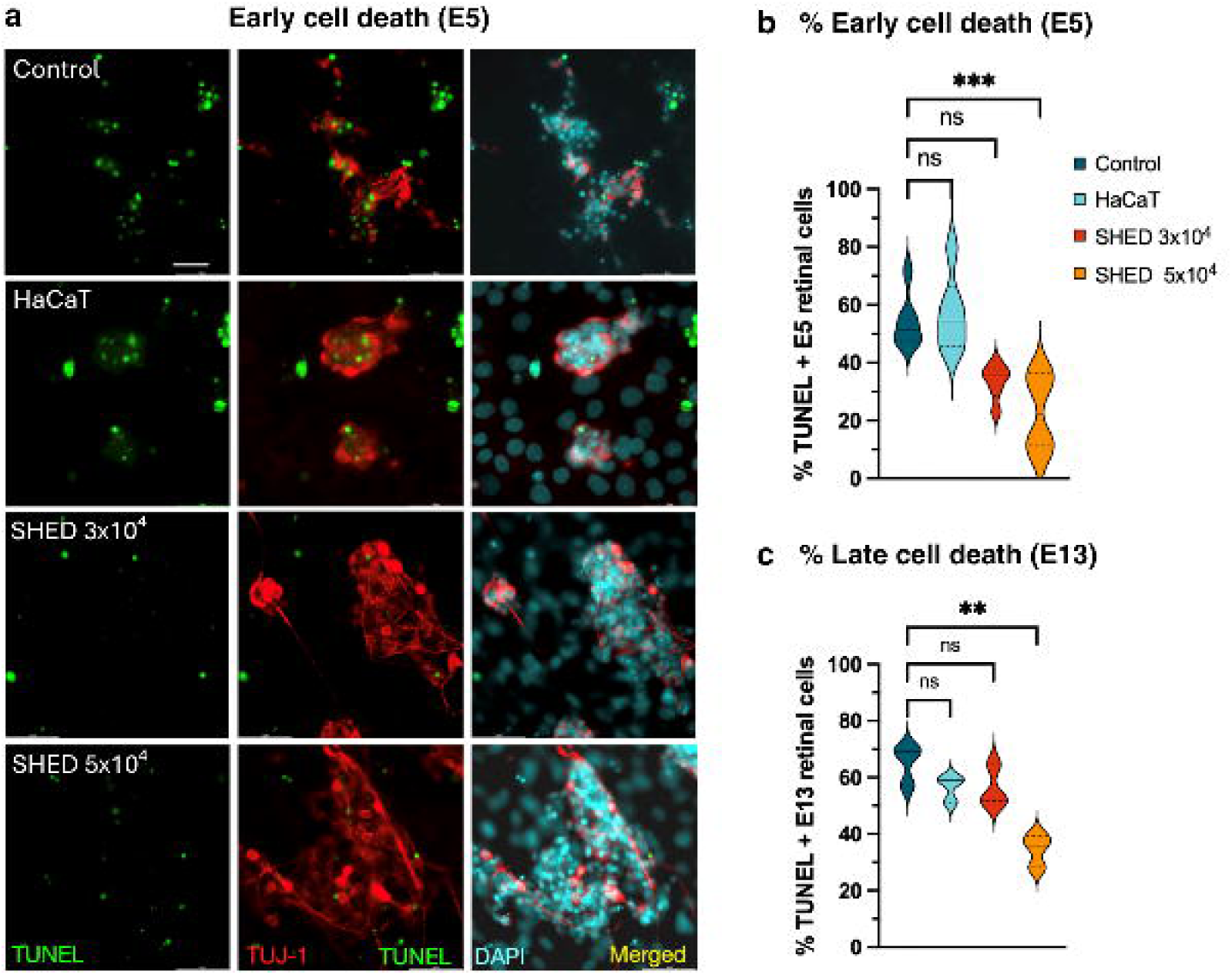
SHED reduce developmental neuronal cell death in embryonic chick retinal cultures. **(a**) Representative fluorescence microscopy images of E5 under different co-culture conditions. Dead nuclei were detected using TUNEL staining (green), RGC were labeled with TUJ-1 (red), and nuclei were counterstained with DAPI (blue). Conditions included control cultures, co-culture with HaCaT cells, and co-culture with SHED at densities of 3 × 10⁴ and 5 × 10⁴ cells. Scale bar = 50 μm. (b) Quantification of the percentage of TUNEL-positive nuclei during early developmental cell death at E5. SHED co-culture significantly reduced apoptosis relative to control and HaCaT conditions in a dose-dependent manner (n=6 biological replicates). (c) Quantification of late apoptotic cell death at E13 under the indicated conditions. SHED-mediated neuroprotection was markedly reduced at this developmental stage compared with E5. (n=3 biological replicates) Statistical analyses were performed using ANOVA. Data are presented as mean ± SEM. *p < 0.05, **p < 0.01, n.s., non-significant.

### SHED preferentially rescue retinal progenitors and nascent retinal ganglion cells in an early developmental stage

To identify which cellular populations were most responsive to SHED-mediated neuroprotection, E5 retinal explants were analyzed using combined TUJ1 and Pax6 immunostaining. Pax6-positive cells corresponded primarily to retinal progenitors and newly differentiating neurons, whereas TUJ1 identified differentiated RGC (Figure 4a).

**Figure 4.**
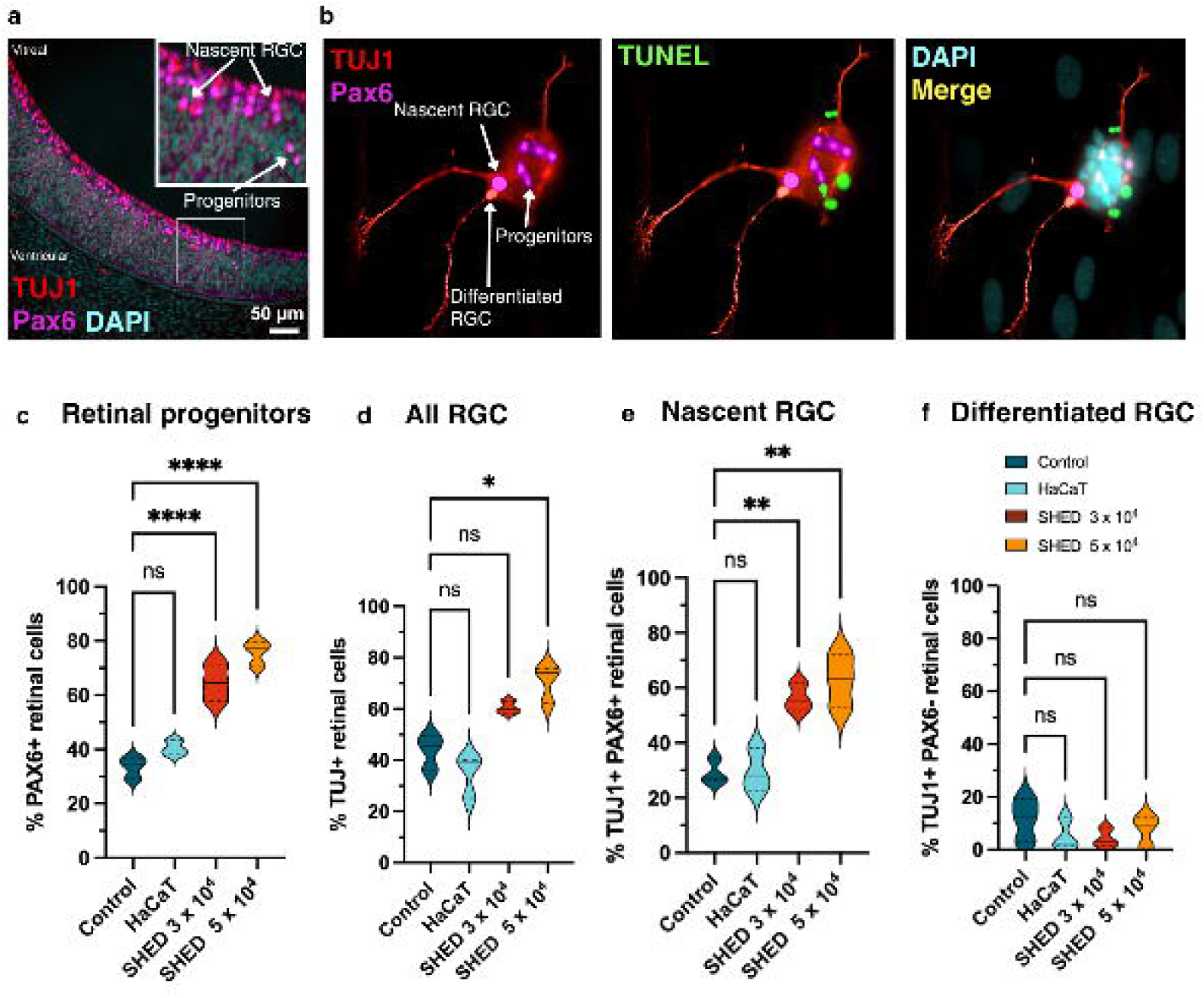
SHED selectively rescue retinal progenitors and newly differentiating neurons during early developmental cell death. (a) Representative immunofluorescence images of E5 chick retinal cultures stained for TUJ-1 (red), Pax6 (green), TUNEL (white/green), and DAPI (blue). Pax6-positive cells correspond to retinal progenitors and newly differentiating neurons. TUJ-1+/Pax6+ cells were identified as nascent RGC, whereas TUJ-1+/Pax6− cells corresponded to differentiated RGC. Scale bar = 50 μm. (b) Quantification of total Pax6-positive retinal progenitor cells under control and SHED-treated conditions. (c) Quantification of TUJ-1+/Pax6+ nascent RGC demonstrating increased survival following SHED treatment. (d) Quantification of TUJ-1+/Pax6− differentiated RGC showing no significant changes between conditions. (e) Quantification of total TUJ-1-positive neuronal populations. Statistical analyses were performed using ANOVA or Student’s t-test. Data are presented as mean ± SEM (n=3 biological replicates).

SHED treatment significantly increased the abundance of all Pax6-positive retinal progenitors and total TUJ1-positive neurons (Figure 4b–c). The most pronounced effect was observed within the TUJ1+/Pax6+ nascent RGC population (Figure 4d), whereas mature TUJ1+/Pax6− neurons remained largely unchanged (Figure 4e). These observations indicate that SHED preferentially rescue immature neuronal populations undergoing developmental commitment, rather than indiscriminately promoting survival across all retinal cell types.

### Proteomic profiling reveals a coordinated neuroregenerative secretome produced by SHED

To investigate the molecular basis of SHED-mediated axon growth, neuroregeneration and neuroprotection, we performed proteomic profiling of SHED-derived secretomes.

Protein abundance analyses identified a highly enriched extracellular environment containing proteins involved in axon guidance, extracellular matrix organization, thrombospondin signaling, IGF signaling and neuroprotection (Figure 5a).Global Reactome mapping (Supplementary Figure 5) and individual Reactome pathway analyses revealed strong enrichment of developmental biology pathways, including axonogenesis, axon guidance, neuronal regeneration and growth cone motility (Figure 5b). Additional enrichment was observed for IGF signaling pathways (Figure 5c) and extracellular matrix organization and remodeling pathways (Figure 5Dd.

**Figure 5.**
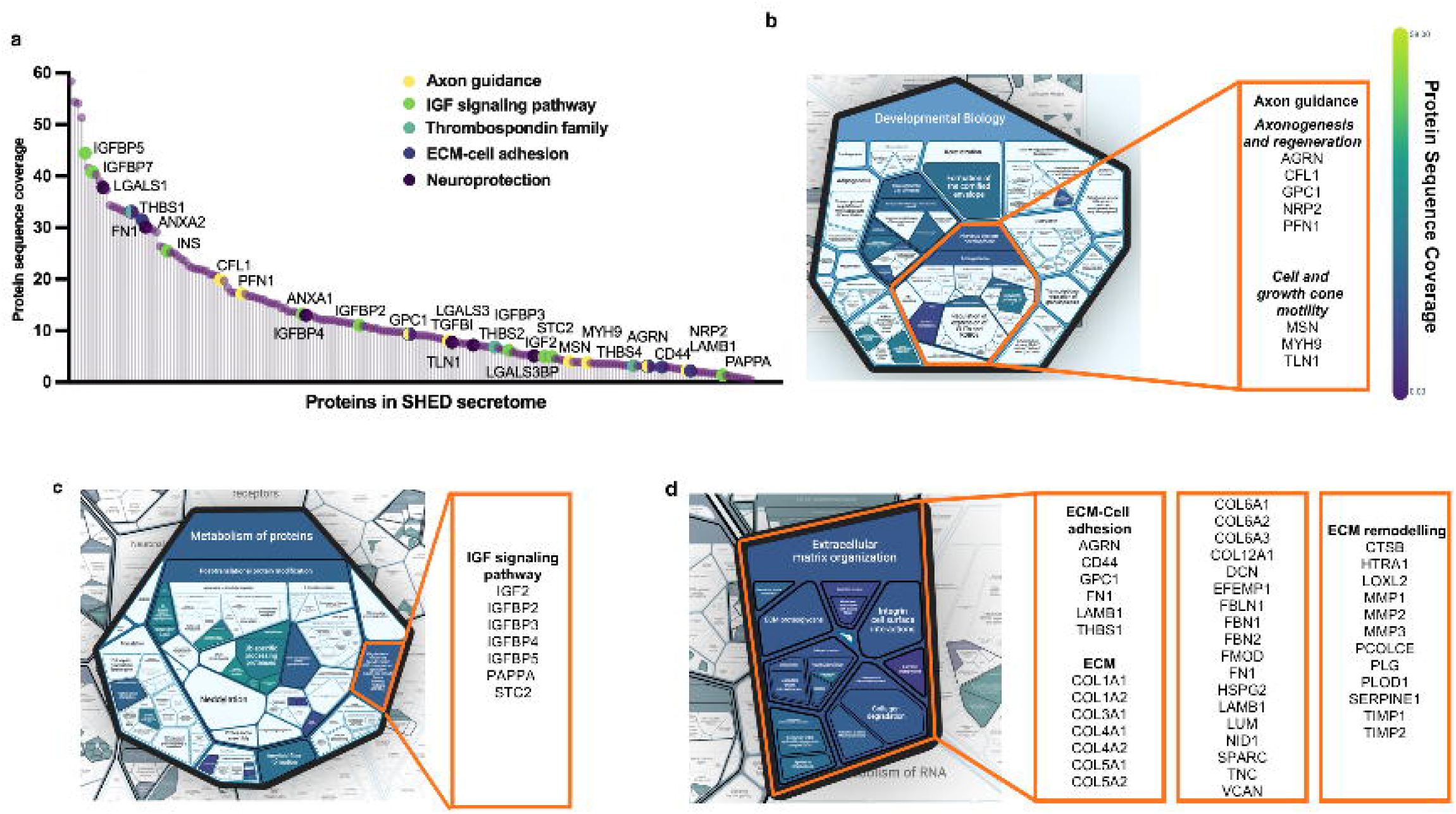
Functional characterization of the SHED secretome reveals enrichment in neurodevelopmental, IGF signalling pathway, and extracellular matrix-associated pathways. (a) Ranked distribution of all proteins identified in the SHED secretome, ordered according to protein sequence coverage, which reflects the relative abundance of detected peptides and, therefore, the representation of each protein within the secretome. Proteins associated with axon guidance, IGF signalling, the thrombospondin family, extracellular matrix (ECM)-cell adhesion, and neuroprotection—selected for detailed investigation in subsequent analyses—are highlighted within the ranking. (B–D) Reactome pathway enrichment analysis showing significantly enriched biological pathways relevant to the present study. Representative proteins contributing to each enriched pathway are indicated. (b) Developmental Biology and Nervous System Development/Axon Guidance pathways. (c) Metabolism of Proteins, including the IGF Signalling Pathway. (d) Extracellular Matrix Organization.

Gene Ontology analyses further validated these findings, revealing enrichment of biological processes associated with nervous system development, neurite morphogenesis, extracellular matrix assembly, cell adhesion and tissue remodeling (Supplementary Figure 6). Collectively, these findings support the existence of a coordinated repertoire of extracellular cues associated with neuroprotection, axonogenesis and regenerative tissue remodeling.

### THBS1–α2δ1 signaling mediates SHED-induced axonal regeneration

Among the proteins identified within the SHED secretome, thrombospondin-1 (THBS1) emerged as a particularly compelling candidate mediator of axonal growth. To investigate its functional contribution, retinal explants were treated with SHED in the presence of a neutralizing anti-THBS1 antibody, gabapentin-mediated inhibition of α2δ1 signaling, or both interventions simultaneously (Figure 6).

**Figure 6.**
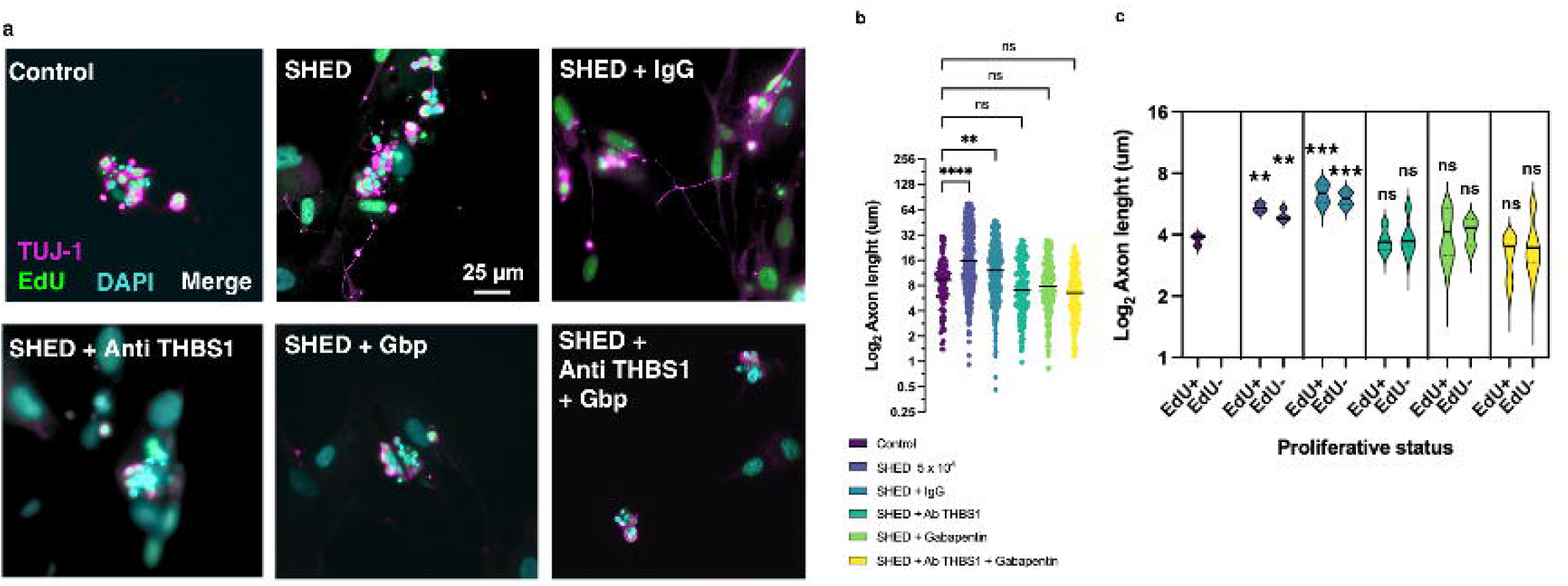
THBS1 signaling mediates SHED-induced axonal regeneration. (a) Representative immunofluorescence images of retinal cultures stained for TUJ-1 (majenta), EdU (green), and DAPI (blue) under the indicated conditions: control, SHED, SHED + IgG (isotype negative control), SHED + anti-THBS1 antibody, SHED + gabapentin (Gbp), and SHED + anti-THBS1 + Gbp. Scale bar = 25 μm. (b) Quantification of average axon length per neuron under the indicated experimental conditions. SHED treatment significantly increased neurite extension, whereas inhibition of THBS1 signaling using neutralizing anti-THBS1 antibody or gabapentin reduced axonal growth to near-control levels (n=8 biological replicates). (c) Quantification of average axon length per neuron in EdU-positive newborn neurons and EdU-negative axotomized RGC under the indicated co-culture conditions (n=4 biological replicates). Statistical analyses were performed using ANOVA with multiple comparisons. Data are presented as mean ± SEM (n=4 biological replicates).

SHED-induced neurite extension was significantly attenuated by either THBS1 neutralization or α2δ1 inhibition in both newborn EdU+ RGC and mature EdU- RGC in co-culture (Figure 6a-c). Combined inhibition did not produce substantial additional effects, suggesting convergence on a common signaling pathway. Despite the observed effects in axon growth, the total number of neurons with axon is only partially reduced ( Supplementary Figure 6b). Morphological analyses revealed not only partially reduced axon formation, but also abnormal neuritic swellings and diminished neurite branching following THBS1 pathway inhibition (Supplementary Figure 6a-c). These findings identify THBS1–α2δ1 signaling as a major mediator of SHED-induced newly axon and regenerative growth.

### Distinct secretory programs independently regulate axonal regeneration and neuroprotection

To determine whether THBS1 signaling also mediated SHED-induced neuronal survival, we examined apoptosis following blockade of THBS1-dependent pathways (using IgG isotype as negative control). Despite THBS1 neutralization neuroprotection induced by the presence of SHED remains in every experimental condition (Figure 7a–b). Moreover, total TUJ1-positive neurons increase induced by SHED is unaltered under THBS1 signaling inhibition (Fig. 7c). These findings suggest that neuronal survival and axonal regeneration are regulated through partially distinct extracellular mechanisms.

**Figure 7.**
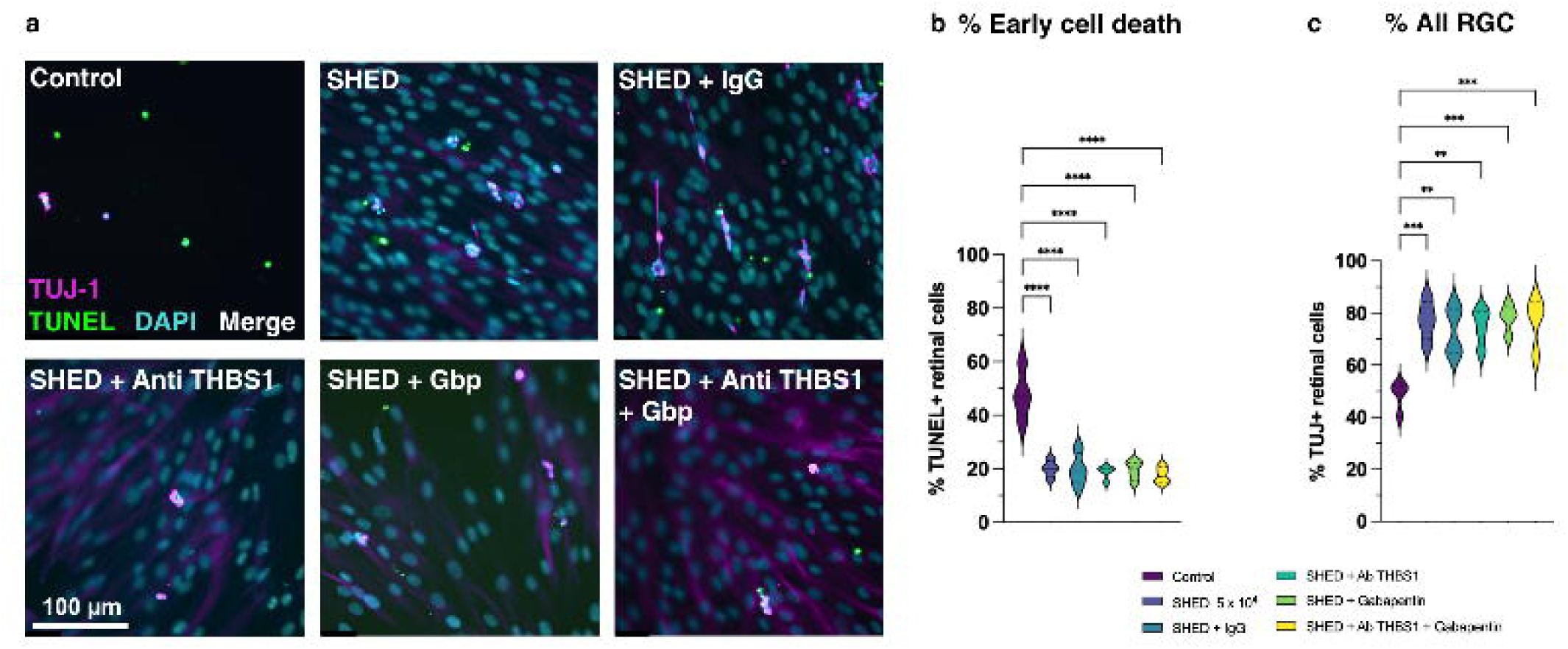
THBS1 signaling blockade does not abolish SHED-mediated neuroprotection. (a) Representative fluorescence images of retinal cultures stained for TUJ-1 (magenta), TUNEL (green), and DAPI (blue) under the indicated conditions: control, SHED, SHED + IgG (isotype negative control), SHED + anti-THBS1 antibody, SHED + gabapentin (Gbp), and SHED + anti-THBS1 + Gbp. Scale bar = 100 μm. (b) Quantification of TUNEL-positive apoptotic nuclei under the indicated experimental conditions. (c) Quantitative analyses demonstrate that neuronal survival remained significantly improved in SHED-treated cultures despite inhibition of THBS1 signaling. Neither anti-THBS1 antibody nor gabapentin abolished the anti-apoptotic rescue effect exerted by SHED. Statistical analyses were performed using ANOVA followed by post hoc comparisons. Data are presented as mean ± SEM (n=4 biological replicates).

### Retinal injury dynamically reprograms the SHED secretome toward a neuroprotective state

To investigate whether SHED respond adaptively to neuronal injury, we compared basal SHED secretomes with secretomes generated following co-culture with retinal explants at E5 (Figure 8a). Differential proteomic analyses identified a distinct damage-responsive secretory profile characterized by selective upregulation of 24 proteins associated with the biological process - programmed cell death, cellular component - extracellular matrix remodeling and stress adaptation (Supplementary Figure 7a). Strikingly, among the 24 identified proteins 10 are associated with neuroprotection (Figure 8b). Among the most strongly enriched proteins were LGALS3, LGALS1, IGFBP3, TNFRSF11B, HSP90AB1, ANXA1 and PRDX1.

**Figure 8.**
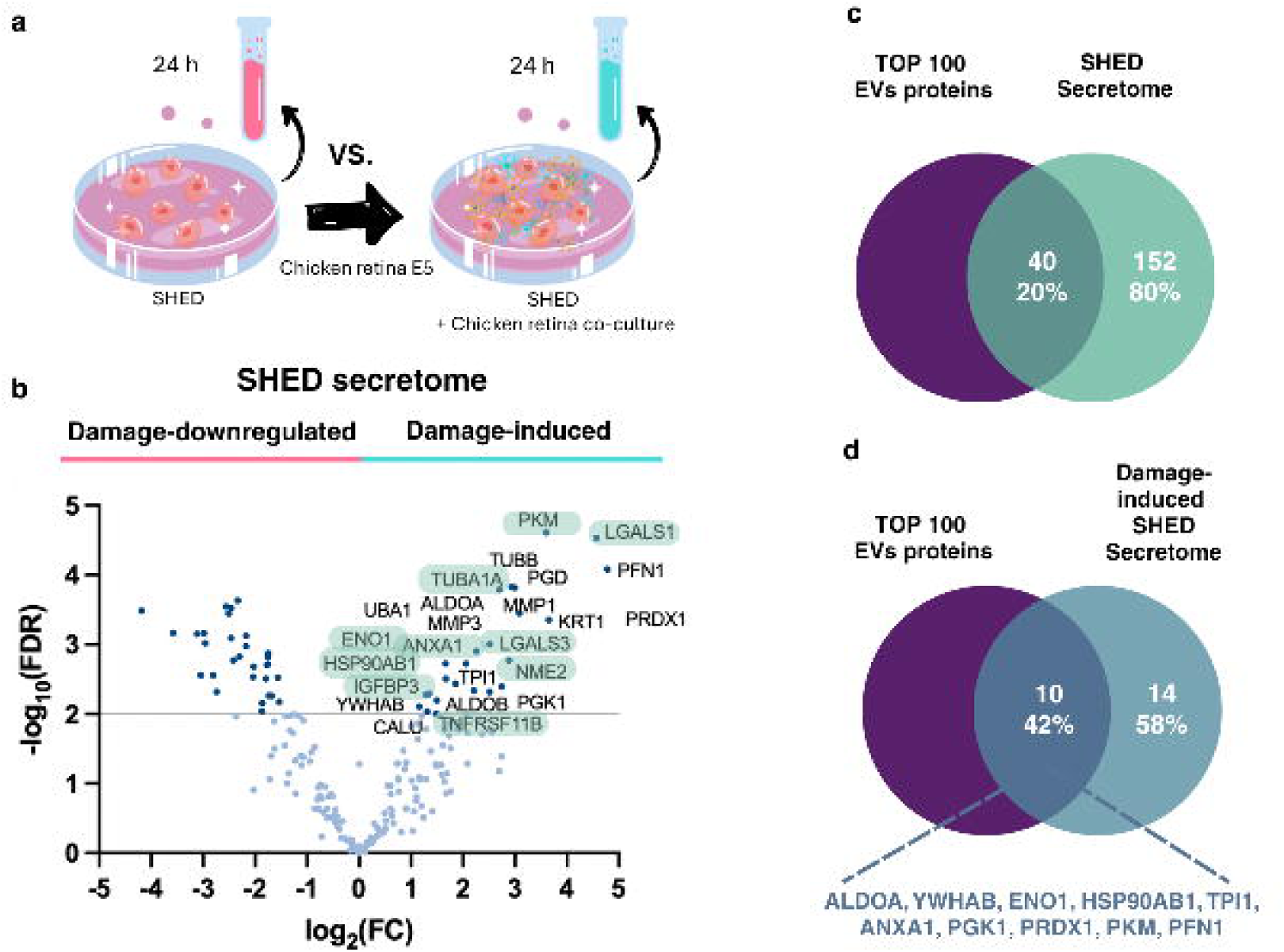
Damage-responsive remodeling of the SHED secretome identifies candidates strongly associated with neuroprotective pathways. (a) Schematic representation of the culture conditions compared in this study and the subsequent analyses performed. (b) Volcano plot comparative analysis identifying 24 significantly upregulated (damage-induced) and 29 downregulated proteins in SHED-retina co-culture secretomes relative to SHED monocultures. Proteins associated with neuroprotective processes according to Gene Ontology annotations are highlighted in green (c)Venn diagram illustrating the overlap between the top 100 most frequently reported extracellular vesicle (EV)-associated proteins listed in Vesiclepedia and the proteins identified in the SHED secretome in the present study. Forty-two of the 192 proteins identified in the SHED secretome overlapped with this curated set of well-established EV-associated proteins. (d) The same analysis was subsequently performed using the 24 proteins found to be upregulated in the presence of injured retinas in the co-culture system, revealing that 10 of these proteins also overlapped with the Vesiclepedia top 100 EV-associated protein dataset.

Comparison with top 100 Vesiclepedia described extracellular vesicle-associated proteins demonstrated substantial overlap between EV cargo and the basal induced SHED secretome (20% of all SHED secreted proteins) (Figure 8c). Interestingly, 42% of the 24 damage-responsive induced proteins are present in extracellular vesicles (Figure 8D).

Collectively, these findings support a model in which SHED dynamically sense injury-associated signals and activate specialized neuroprotective and regenerative secretory programs. Whereas THBS1-dependent signaling appears to drive axonogenesis and regenerative growth, injury-induced EV associated secretory responses may contribute predominantly to neuronal survival, suggesting the existence of mechanistically distinct but complementary regenerative pathways.

## DISCUSSION

SHED have emerged as a particularly attractive source of mesenchymal stem cells due to their neural crest origin, accessibility, high proliferative capacity, and intrinsic neurogenic potential (4,13). Consistent with previous reports, our data demonstrate that SHED maintain stable fibroblast-like morphology and robust proliferative behavior during in vitro expansion, outperforming adult DPSC, which displayed morphological alterations compatible with reduced proliferative fitness and early senescence. These observations further support the concept that SHED represent a biologically immature and highly plastic mesenchymal stem cell population with enhanced regenerative potential. Importantly, SHED consistently expressed canonical mesenchymal markers, including CD105, CD73, CD90, vimentin, and STRO-1, confirming preservation of stem cell identity across donors and passages.

The present study demonstrates that SHED exert robust neurotrophic, neuroprotective, and pro-regenerative effects in developing and mature retinal tissue. Using an embryonic chick retina model that enables discrimination between active developmental axonogenesis and regenerative responses following axotomy, we show that SHED-derived signals promote neuronal survival, enhance axonal growth, support retinal progenitor maintenance, and stimulate neuronal differentiation. In addition to the fact that globally the secretome composition constitutes a protein niche ideally suited to mediate its regenerative and protective effects, mechanistically, our findings identify THBS1-dependent signaling as a major contributor to these effects, highlighting extracellular matrix-associated pathways as critical mediators of SHED-induced neuroregeneration.

A major strength of the present work lies in the use of the embryonic chick retina, a model that offers a unique opportunity to dissect distinct biological processes occurring during neurodevelopment and regeneration. The chick retina exhibits a well-defined temporal organization in which retinal ganglion cell (RGC) neurogenesis and developmental axonogenesis occur predominantly between embryonic days 2 and 7, whereas later stages are characterized by synaptic refinement, target innervation, and programmed neuronal elimination (1). At embryonic day 5 (E5), retinal tissue contains proliferating neuroblasts and newly differentiating RGC actively undergoing axonogenesis, while embryonic day 13 (E13) retinas represent a more mature neuronal environment with intrinsically reduced regenerative competence. By combining EdU incorporation with TUJ-1 immunostaining, we were able to discriminate newly generated neurons from pre-existing axotomized neurons, allowing independent analysis of developmental axonogenesis and regenerative neurite growth.

Our findings demonstrate that SHED significantly enhance axonal extension in both immature and mature retinal explants. Notably, SHED-mediated effects were observed not only in newborn neurons undergoing physiological axonogenesis, but also in mature axotomized neurons attempting regenerative growth. This dual responsiveness is particularly relevant because regeneration of central nervous system axons is severely restricted in mature neurons due to both intrinsic transcriptional limitations and extrinsic inhibitory cues (14,15). The ability of SHED-derived factors to partially overcome these barriers suggests that the SHED secretome contains signals capable of reactivating conserved neuronal growth programs even in differentiated neural tissue.

Importantly, SHED treatment enhanced axonal outgrowth in both EdU+/TUJ-1+ newly generated neurons and EdU−/TUJ-1+ mature neurons at E5, indicating that SHED-derived signals support both *de novo* axonogenesis and regenerative neurite extension. Unlike classical neurotrophic factors such as BDNF or IGF-1, which frequently exert selective effects on defined neuronal subpopulations (2,16), the SHED secretome appears to broadly influence retinal neurons across maturational states. Interestingly, SHED treatment did not significantly alter proliferation rates or bias progenitor differentiation toward the RGC lineage, suggesting that the observed regenerative effects are primarily mediated through modulation of post-mitotic neuronal behavior rather than alterations in progenitor expansion.

In addition to promoting neurite extension, secretome-derived cues exerted potent neuroprotective effects by significantly reducing cell death in both E5 and E13 retinal explants. The chick retina undergoes tightly regulated waves of programmed cell death during development, including early apoptosis associated with neurogenesis and later neurotrophic cell death linked to synaptic maturation and target innervation (17,18). The reduction in TUNEL-positive neurons following SHED treatment suggests that SHED-derived trophic support enhances neuronal resilience across multiple developmental contexts. Importantly, the anti-apoptotic effects observed at E13 indicate that SHED-secreted factors remain effective even in relatively mature tissue undergoing target-dependent pruning. These findings are consistent with previous studies demonstrating neuroprotective effects of hMSC secretomes in models of retinal degeneration, spinal cord injury, and neurodegenerative disease (8,19,20).

These findings provide important insight into the developmental specificity of SHED-mediated neuroprotection. Rather than broadly increasing neuronal survival across all retinal populations, SHED preferentially rescued retinal progenitors and newly differentiating neurons that are normally subjected to early developmental cell death at E5. This selective effect is particularly relevant in the context of retinal neurogenesis, where programmed cell death is thought to function as a physiological mechanism to regulate neuronal number and refine early neural circuits during development. The increase in Pax6-positive progenitors together with the expansion of the TUJ-1+/Pax6+ nascent neuronal population suggests that SHED-derived signals specifically support cells undergoing the transition from progenitor state to neuronal differentiation. Importantly, the absence of changes in mature TUJ-1+/Pax6− neurons indicates that SHED do not simply induce generalized neuronal proliferation or indiscriminate survival but instead modulate a temporally restricted developmental window associated with neuronal commitment and early differentiation.

This observation is consistent with the concept that newly differentiating neurons represent a particularly vulnerable cellular population during early retinogenesis (21,22). Our results suggest that SHED-derived secreted factors can interfere with this developmental checkpoint, promoting survival of newly generated neurons that would otherwise be eliminated. Such activity may partially recapitulate endogenous developmental trophic support programs normally provided by neighboring retinal or glial populations during neurogenesis.

Rather than consisting of a generic trophic secretome, SHED produced a highly specialized repertoire of extracellular matrix-associated and neurodevelopment-related proteins linked to axonogenesis, neurite morphogenesis, cell adhesion, and neuronal survival. Particularly notable was the enrichment of thrombospondins, galectins, and IGFextracellular pathway related proteins, all of which have previously been implicated in nervous system development and injury responses. Thrombospondins are well established as astrocyte-derived synaptogenic molecules that promote neurite extension and synapse formation through α2δ-1-dependent mechanisms (23–26), whereas galectins have emerged as important regulators of inflammatory signaling, neuronal survival, and tissue remodeling following neural injury (27–32). Similarly, IGFBPs and Pappa are increasingly recognized not only as modulators of IGF bioavailability, but also as regulators of extracellular matrix interactions and neuronal differentiation.Importantly, gene ontology and Reactome pathway analyses indicated strong enrichment in pathways associated with nervous system development, extracellular matrix organization and axon guidance. These findings suggest that SHED do not act exclusively through passive trophic support but instead recreate molecular programs resembling those active during embryonic neural development and regenerative remodeling. The enrichment of extracellular matrix and integrin-associated pathways is particularly interesting, as extracellular matrix dynamics are known to critically regulate growth cone motility, neurite stabilization, and progenitor differentiation during CNS development. In this context, SHED may generate a permissive developmental-like microenvironment capable of simultaneously supporting neuronal survival and structural regeneration. Together, these findings support the concept that SHED-mediated regeneration is orchestrated through a complex trophic and extracellular signaling environment capable of recreating permissive developmental-like conditions for neuronal growth.

Among the identified factors, THBS1 emerged as a particularly compelling mediator. Thrombospondins are matricellular glycoproteins secreted by astrocytes and supportive glial populations that regulate synaptogenesis, neurite extension, extracellular matrix remodeling, and neuronal survival (24,27). THBS1 has previously been shown to promote synapse formation through interaction with the neuronal α2δ-1 receptor, a pathway pharmacologically inhibited by gabapentin (27). Our findings strongly support a functional role for THBS1 signaling in SHED-mediated neuroregeneration, as neutralization of THBS1 significantly impaired axonal growth, reduced neuronal survival, and induced pronounced neuritic abnormalities.

Importantly, combined inhibition using anti-THBS1 antibodies and gabapentin produced an even greater impairment of regenerative responses, suggesting that α2δ-1- dependent thrombospondin signaling contributes substantially to the trophic effects exerted by SHED. The appearance of rounded neuritic swellings following THBS1 inhibition is particularly intriguing, as these structures resemble axonal dystrophies associated with cytoskeletal disorganization, impaired axonal transport, or abortive regenerative attempts. Similar dystrophic neurites have been described in contexts of axonal stress and neurodegeneration, including traumatic injury and Alzheimer’s disease-associated pathology (33,34). These findings suggest that THBS1 signaling may not only stimulate neurite extension but also stabilize axonal architecture and preserve neurite integrity during regeneration.

The diverse composition of the SHED secretome likely explains the broad biological effects observed across developmental stages. In addition to THBS1, proteins such as fibronectin and IGF pathways may contribute to axonal pathfinding, metabolic support, extracellular matrix remodeling, and neuronal stabilization. The coexistence of structural extracellular matrix proteins together with trophic modulators suggests that SHED establish a permissive extracellular microenvironment that provides both biochemical and biophysical support for neuronal regeneration. This notion is consistent with increasing evidence indicating that hMSC-mediated tissue repair largely depends on paracrine signaling rather than direct tissue replacement (35,36). Strikingly, blockade of THBS1 signaling failed to suppress the neuroprotective rescue effect exerted by SHED, demonstrating that neuronal survival and axonal regeneration are controlled through mechanistically separable pathways. This distinction represents one of the most relevant findings of the present study, as regenerative growth and neuronal survival are frequently assumed to rely on overlapping trophic mechanisms. Instead, our results support a model in which SHED simultaneously activate independent extracellular programs dedicated to structural regeneration and apoptosis suppression. Such mechanistic divergence may have important translational implications, suggesting that distinct components of the SHED secretome could potentially be optimized to preferentially enhance either neuroprotection or regenerative growth depending on the pathological context.

The proteomic analysis of SHED-retina co-culture secretomes further supports the existence of an active developmental and regenerative signaling environment generated in response to injured retinal tissue. Consistent with this interpretation, damage-responsive secretome analyses identified a second injury-induced molecular program enriched in proteins associated with programmed cell death regulation and neuronal survival. Among the most strongly upregulated proteins were LGALS3 (Galectin-3), LGALS1 (Galectin-1), IGFBP3, ANXA1, PRDX1, and TNFRSF11B, several of which have been implicated in inflammatory modulation, tissue remodeling, and neuroprotection following CNS injury. In particular, Galectin-3 emerged as a compelling candidate mediator of the SHED-induced rescue effect (31). Galectin-3 has previously been described as an injury-responsive lectin capable of modulating microglial activation, phagocytosis, inflammatory signaling, and neuronal survival through TREM2-associated pathways. While additional mechanistic studies will be necessary to directly establish causality, our findings support the hypothesis that a Galectin-3/TREM2- associated axis contributes to the anti-apoptotic effects observed in SHED-treated retinas.

An additional important observation was the remarkably limited contribution of retina-derived proteins to the co-culture secretome. Only 19 proteins from the Gallus gallus UniProtKB database were identified at low expression levels. However, the detected peptides also showed 100% sequence homology with human UniProtKB entries, preventing their unambiguous assignment to the chicken proteome. Of these proteins, only two were found to be upregulated in SHED co-cultures exposed to damaged chicken retinal cells indicating that embryonic retinal cultures constitute a low-secretory cellular environment under these experimental conditions. This finding strengthens the conclusion that the vast majority of extracellular signaling molecules detected in co-cultures originate from SHED rather than from the retinal tissue itself. Consequently, the regenerative and neuroprotective phenotypes observed are likely driven predominantly by SHED-derived extracellular programs dynamically induced in response to neuronal injury.

Collectively, these findings support a model in which SHED respond to neuronal injury by activating coordinated but mechanistically distinct secretory programs that independently regulate axonal regeneration and neuronal survival. Due to their neural crest origin, SHED may retain developmental signaling properties particularly suited for supporting neural plasticity, tissue remodeling, and neuroregeneration. This developmental identity may distinguish SHED from other hMSC populations and help explain their remarkable ability to recreate embryonic-like neurodevelopmental and regenerative environments within injured neural tissue. Beyond retinal regeneration, these observations may have broader implications for neurodevelopmental and neurodegenerative disorders characterized by impaired neuronal survival, axonal dysfunction, or defective extracellular matrix remodeling.

Another important aspect deserving further investigation is the potential contribution of SHED-derived extracellular vesicles (SHED-EVs). SHED-EVs derived from hMSC are increasingly recognized as important mediators of intercellular communication through delivery of proteins, lipids, and regulatory RNAs. Several studies have shown that MSC-derived EVs can reproduce many regenerative effects of their parental cells (37). It is therefore possible that a substantial component of the neuroprotective and axonogenic activity observed in this study is mediated through EV-associated cargo. Isolation and functional characterization of SHED-derived EVs may therefore provide a scalable and clinically attractive therapeutic strategy.

The translational implications of these findings are considerable. RGC degeneration and axonal loss are central pathological features of glaucoma, optic neuropathies, traumatic injury, and several neurodegenerative conditions. Current therapeutic approaches primarily aim to slow disease progression but do not effectively promote neuronal survival or axonal regeneration (38). The demonstration that SHED-derived factors can simultaneously enhance neuronal survival, stimulate axonal growth, and support neurodevelopmental programs suggests that SHED-based therapies could represent a promising strategy for regenerative ophthalmology and central nervous system repair.

More broadly, our findings reinforce the idea that mature neurons retain a latent competence for regeneration that can be reactivated under appropriate extracellular conditions. This challenges the traditional view that regenerative capacity is irreversibly lost following developmental maturation (39). Instead, our data support a model in which permissive trophic and extracellular matrix-associated signaling can unlock conserved developmental programs in postmitotic neurons, consistent with observations from optic nerve and spinal cord injury models where manipulation of the extracellular environment restores axonal growth competence (40,41).

Several limitations of the present study should also be acknowledged. Although the chick retina provides a powerful developmental and regenerative model, additional validation in mammalian systems will be required to determine the translational relevance of these findings. Furthermore, while THBS1 appears to play a major role, the SHED secretome contains numerous additional extracellular matrix and trophic factors that likely act synergistically to mediate the observed effects. A key limitation of this study is the lack of functional characterization of galectin-1 and galectin-3, as both were among the most strongly upregulated proteins in response to the presence of injured retina in the co-culture system. Their marked induction suggests a potentially important contribution to the neuroprotective effects observed, which warrants further investigation. Future studies integrating transcriptomic, proteomic, and functional perturbation approaches, including neutralizing antibodies, RNA interference, or CRISPR-based strategies will be necessary to identify the minimal set of effectors required for optimal neuroregenerative activity. Likewise, characterization of downstream signaling pathways activated in retinal neurons, including PI3K/Akt, mTOR, and MAPK cascades, may provide additional mechanistic insight into how SHED-derived signals converge with intrinsic neuronal growth programs.

In conclusion, our study demonstrates that SHED own potent neuroprotective and neuroregenerative properties capable of enhancing neuronal survival, axonal regeneration, and retinal neurodevelopment across distinct developmental stages. These effects are mediated predominantly through paracrine mechanisms involving a complex secretome enriched in extracellular matrix-associated and neurotrophic factors, with THBS1-dependent signaling emerging as a major mechanistic contributor. By combining a temporally defined chick retina model with human-derived stem cells, this work establishes a translational platform for investigating regenerative signaling during neural development and injury. Collectively, our findings further support SHED as a promising human stem cell source for regenerative neuroscience and provide new mechanistic insights into extracellular pathways capable of reactivating neuronal growth and repair in the central nervous system. Importantly, this work also establishes the developing and injured chick retina as a versatile biological platform that recapitulates key processes of neuronal degeneration, survival, and regeneration, enabling future studies aimed at dissecting regenerative mechanisms and assessing the efficacy of diverse cell-based and EV-based therapeutic approaches.

