## Supplementary Figures 1 to 8 for "Human SHED-derived extracellular cues activate a specialized neuroprotective and regenerative program in developing retinal ganglion cells"

#### Supplementary Figure 1

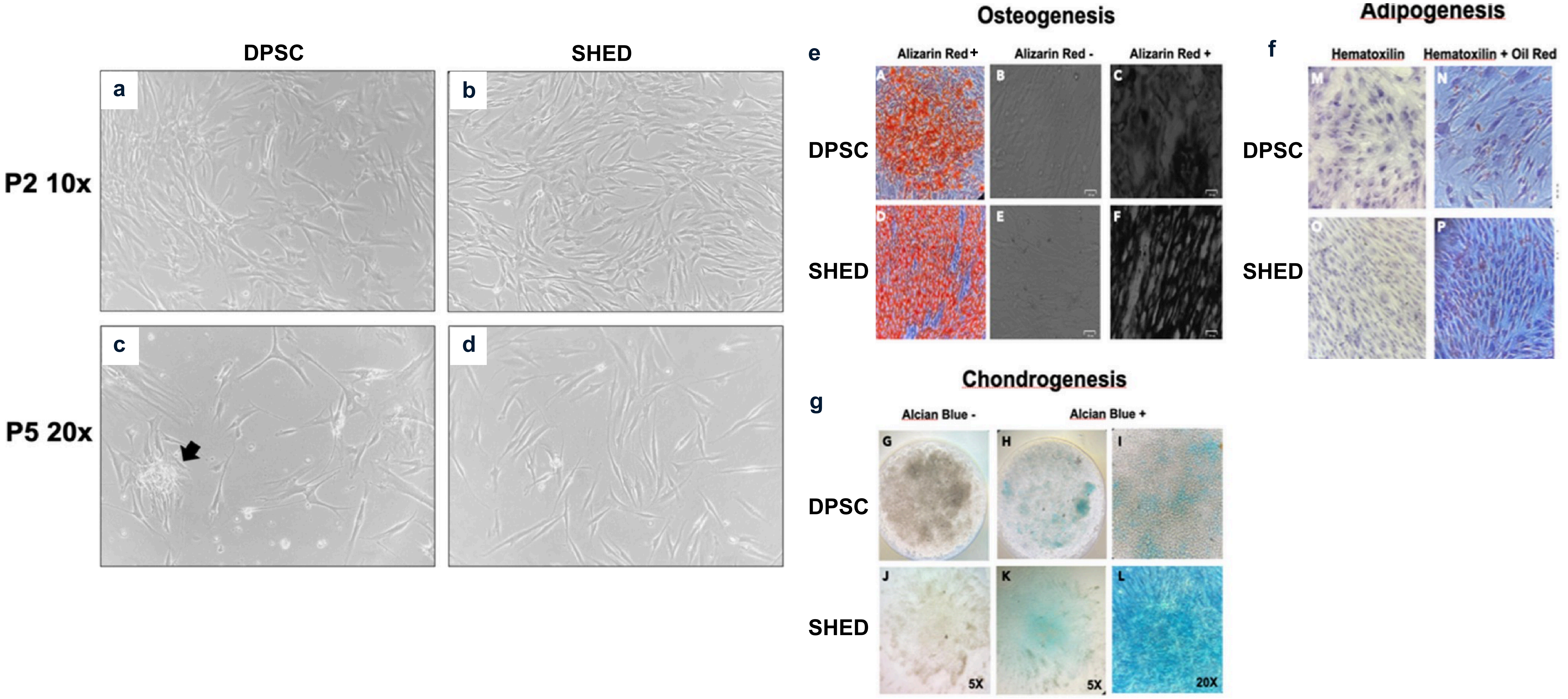

Supplementary Figure 2

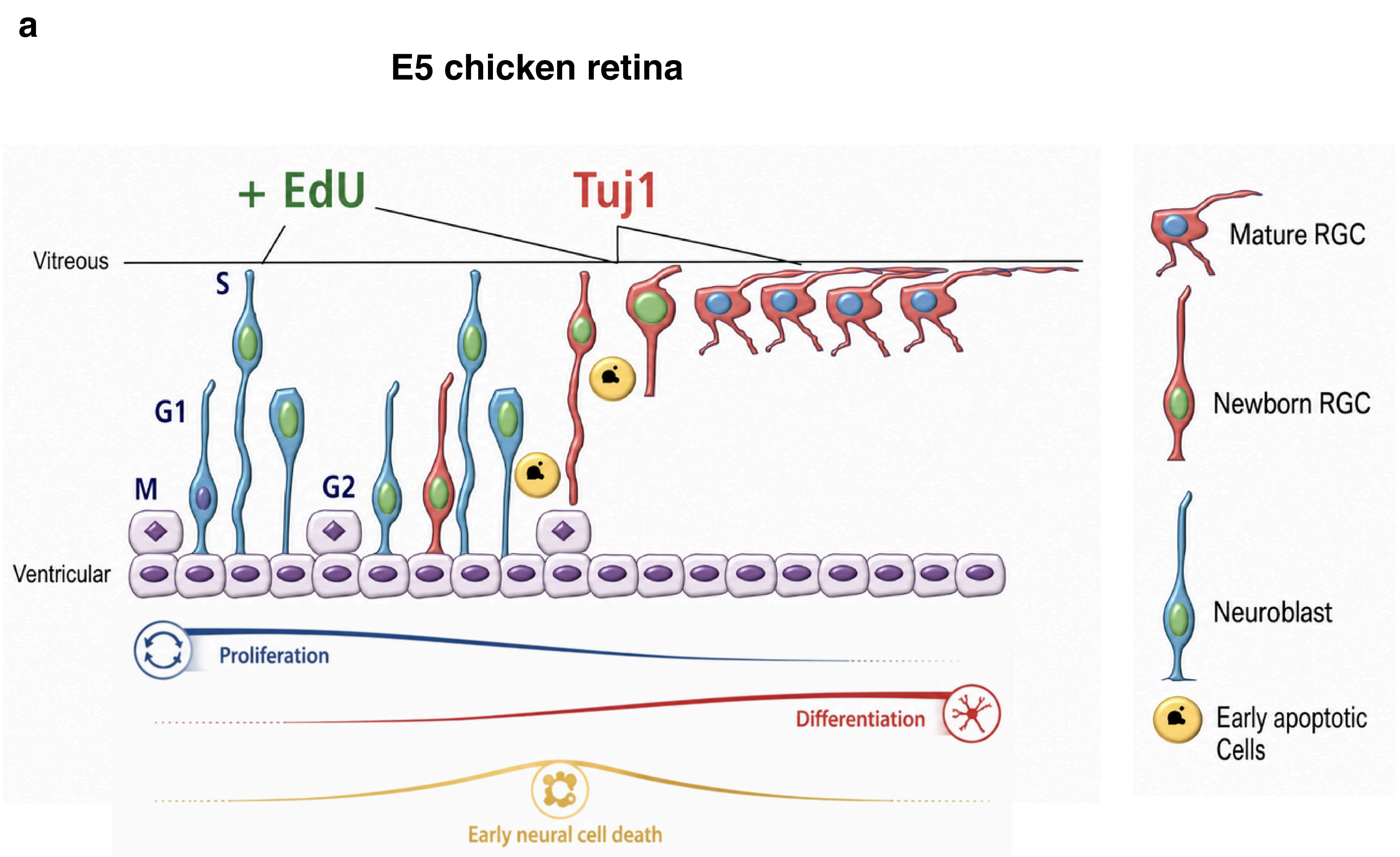


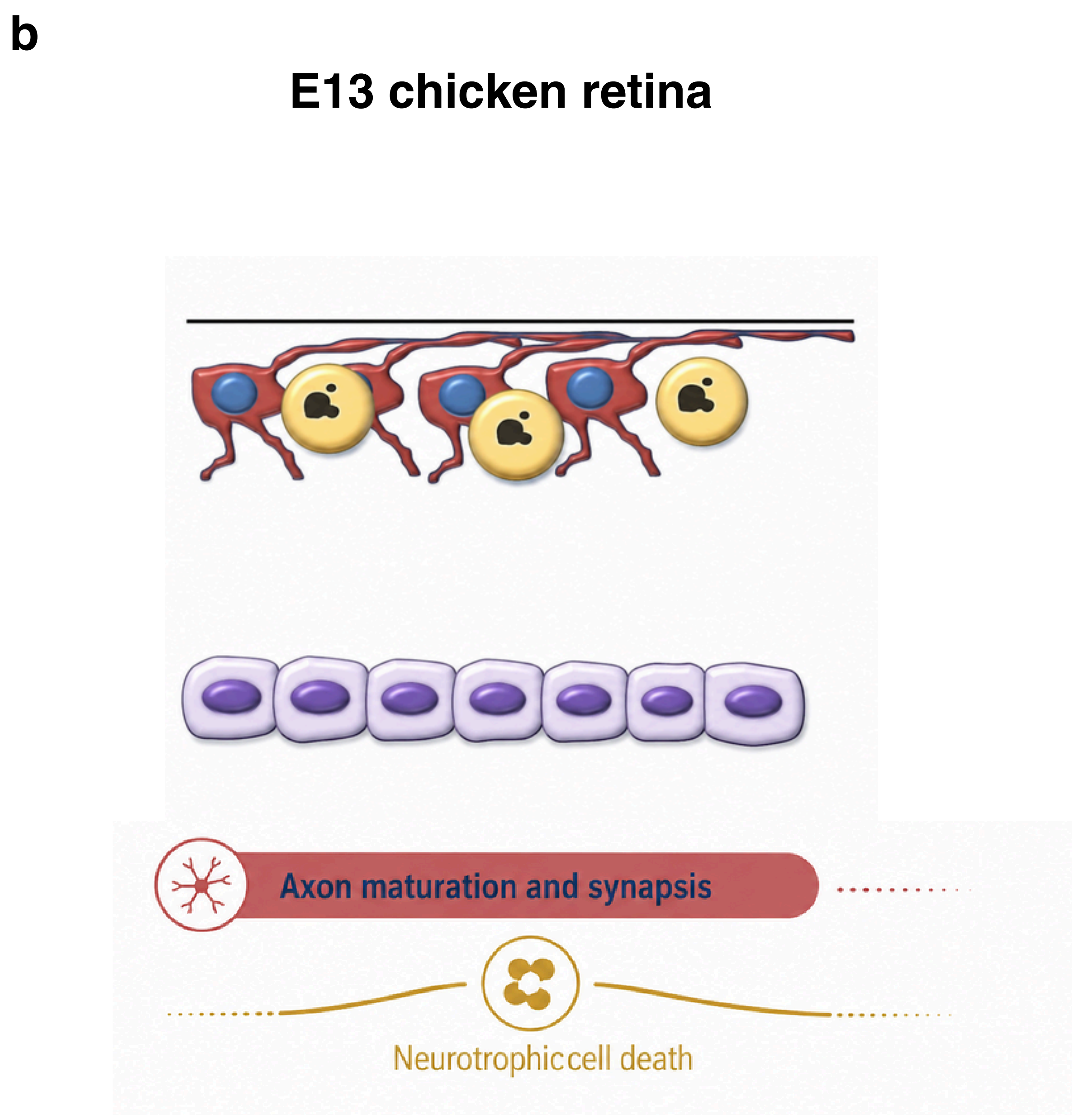

Supplementary figure 3

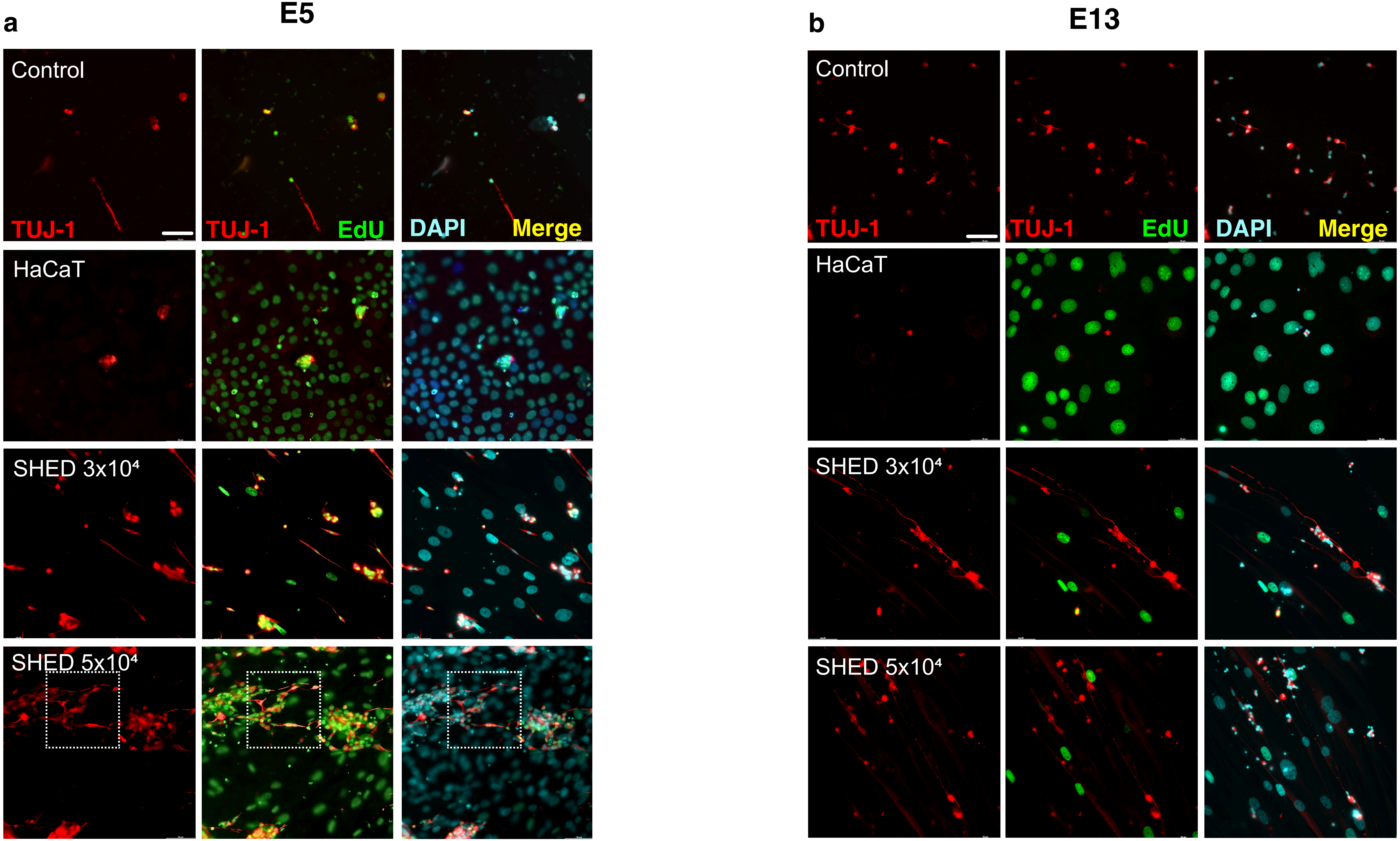

**Supplementary figure 4**

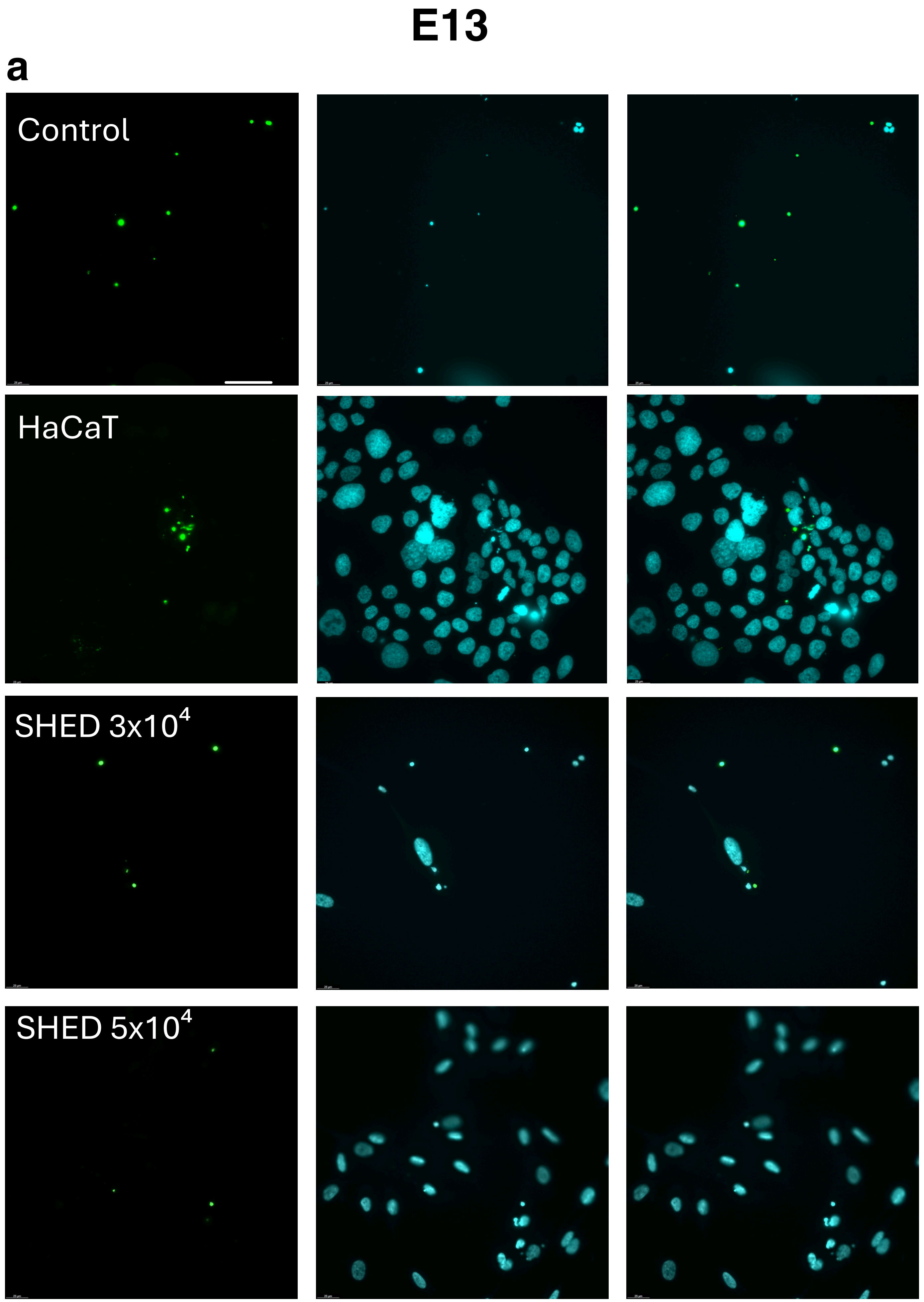

**Supplementary Figure 5**

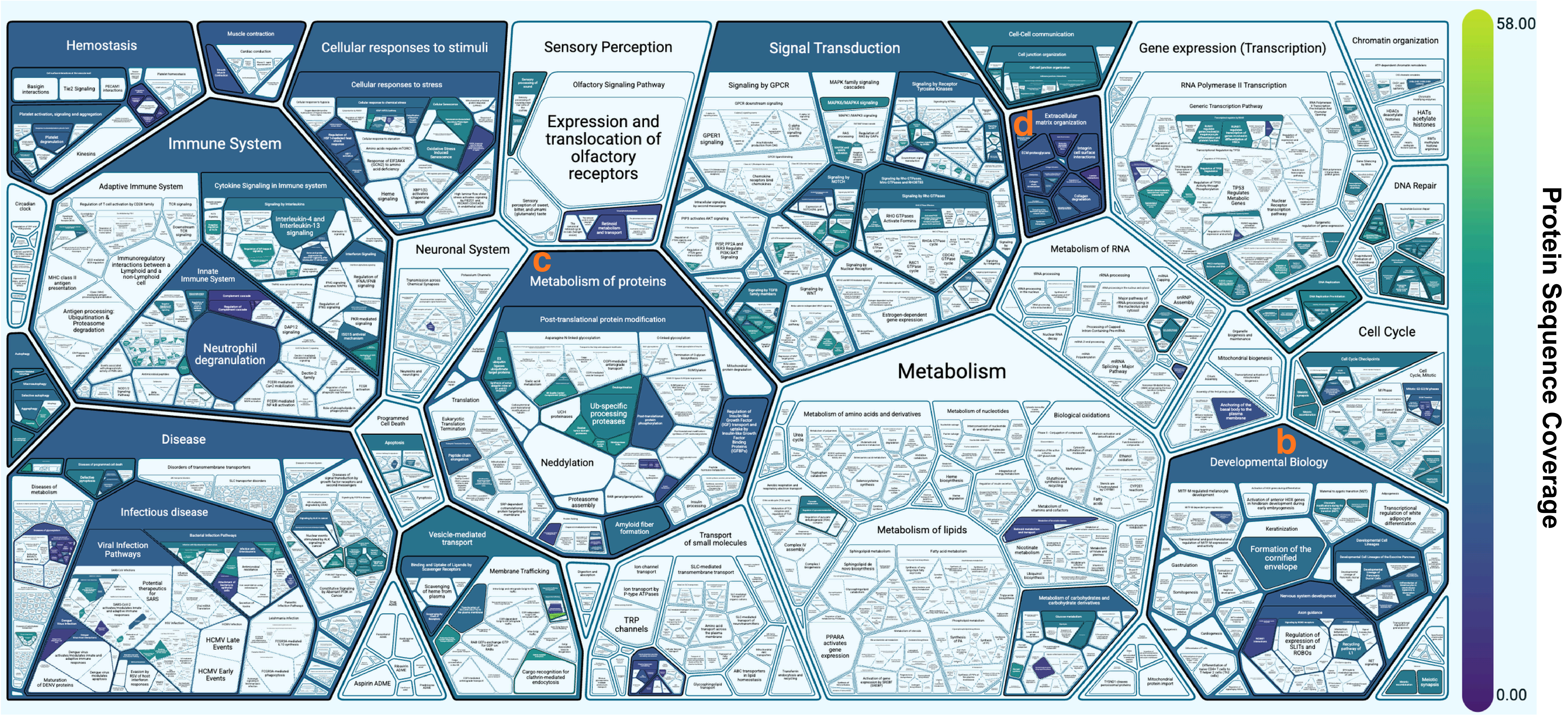

#### Supplementary Figure 6

**a**

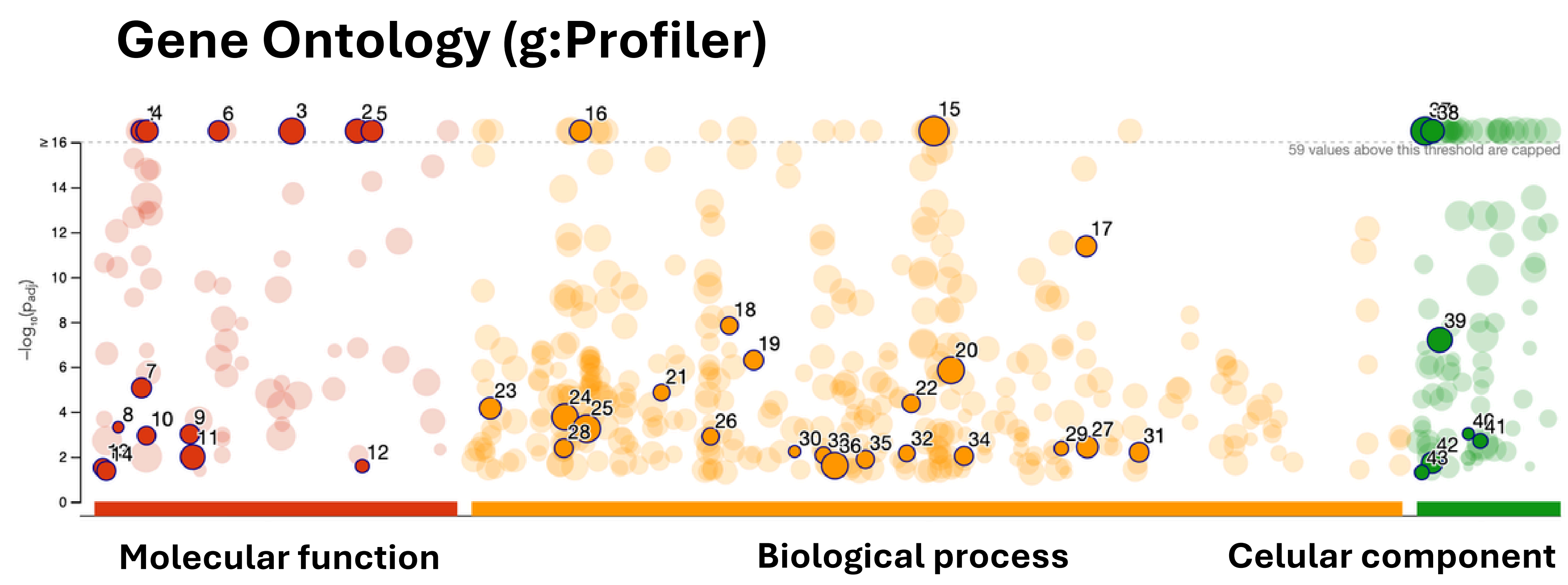


**b**

| ID | Source | Term ID | Term Name | $p_{adj}$ (query_... |
|---|---|---|---|---|
| 1 | GO:MF | GO:0005201 | extracellular matrix structural constituent | $3.570\times10^{-40}$ |
| 2 | GO:MF | GO:0050839 | cell adhesion molecule binding | $1.966\times10^{-25}$ |
| 3 | GO:MF | GO:0044877 | protein-containing complex binding | $1.318\times10^{-23}$ |
| 4 | GO:MF | GO:0005539 | glycosaminoglycan binding | $2.470\times10^{-21}$ |
| 5 | GO:MF | GO:0061134 | peptidase regulator activity | $7.508\times10^{-20}$ |
| 6 | GO:MF | GO:0019838 | growth factor binding | $3.739\times10^{-18}$ |
| 7 | GO:MF | GO:0005200 | structural constituent of cytoskeleton | $8.612\times10^{-6}$ |
| 8 | GO:MF | GO:0004332 | fructose-bisphosphate aldolase activity | $4.822\times10^{-4}$ |
| 9 | GO:MF | GO:0016209 | antioxidant activity | $9.826\times10^{-4}$ |
| 10 | GO:MF | GO:0005507 | copper ion binding | $1.145\times10^{-3}$ |
| 11 | GO:MF | GO:0016491 | oxidoreductase activity | $1.024\times10^{-2}$ |
| 12 | GO:MF | GO:0051920 | peroxiredoxin activity | $2.604\times10^{-2}$ |
| 13 | GO:MF | GO:0001786 | phosphatidylserine binding | $3.088\times10^{-2}$ |
| 14 | GO:MF | GO:0003725 | double-stranded RNA binding | $4.245\times10^{-2}$ |
| 15 | GO:BP | GO:0048856 | anatomical structure development | $2.063\times10^{-26}$ |
| 16 | GO:BP | GO:0007596 | blood coagulation | $2.190\times10^{-17}$ |
| 17 | GO:BP | GO:0072524 | pyridine-containing compound metabolic proc... | $4.268\times10^{-12}$ |
| 18 | GO:BP | GO:0031638 | zymogen activation | $1.464\times10^{-8}$ |
| 19 | GO:BP | GO:0032963 | collagen metabolic process | $5.040\times10^{-7}$ |
| 20 | GO:BP | GO:0051246 | regulation of protein metabolic process | $1.403\times10^{-6}$ |
| 21 | GO:BP | GO:0018149 | peptide cross-linking | $1.381\times10^{-5}$ |
| 22 | GO:BP | GO:0048009 | insulin-like growth factor receptor signaling pa... | $4.315\times10^{-5}$ |
| 23 | GO:BP | GO:0001906 | cell killing | $6.897\times10^{-5}$ |
| 24 | GO:BP | GO:0007010 | cytoskeleton organization | $1.684\times10^{-4}$ |
| 25 | GO:BP | GO:0009056 | catabolic process | $5.601\times10^{-4}$ |
| 26 | GO:BP | GO:0030212 | hyaluronan metabolic process | $1.246\times10^{-3}$ |
| 27 | GO:BP | GO:0072593 | reactive oxygen species metabolic process | $3.713\times10^{-3}$ |
| 28 | GO:BP | GO:0006956 | complement activation | $4.161\times10^{-3}$ |
| 29 | GO:BP | GO:0071492 | cellular response to UV-A | $4.269\times10^{-3}$ |
| 30 | GO:BP | GO:0035583 | sequestering of TGFbeta in extracellular matrix | $5.835\times10^{-3}$ |
| 31 | GO:BP | GO:0098869 | cellular oxidant detoxification | $6.259\times10^{-3}$ |
| 32 | GO:BP | GO:0046716 | muscle cell cellular homeostasis | $7.041\times10^{-3}$ |
| 33 | GO:BP | GO:0042026 | protein refolding | $8.575\times10^{-3}$ |
| 34 | GO:BP | GO:0051881 | regulation of mitochondrial membrane potential | $9.259\times10^{-3}$ |
| 35 | GO:BP | GO:0044788 | modulation by host of viral process | $1.299\times10^{-2}$ |
| 36 | GO:BP | GO:0042592 | homeostatic process | $2.478\times10^{-2}$ |
| 37 | GO:CC | GO:0005615 | extracellular space | $1.944\times10^{-118}$ |
| 38 | GO:CC | GO:0005925 | focal adhesion | $6.354\times10^{-19}$ |
| 39 | GO:CC | GO:0009986 | cell surface | $6.250\times10^{-8}$ |
| 40 | GO:CC | GO:0034363 | intermediate-density lipoprotein particle | $9.526\times10^{-4}$ |
| 41 | GO:CC | GO:0042627 | chylomicron | $1.979\times10^{-3}$ |
| 42 | GO:CC | GO:0005912 | adherens junction | $1.858\times10^{-2}$ |
| 43 | GO:CC | GO:0001527 | microfibril | $4.968\times10^{-2}$ |

**Molecular function**

**Biological process**

**Celular component**

**Supplementary Figure 7**

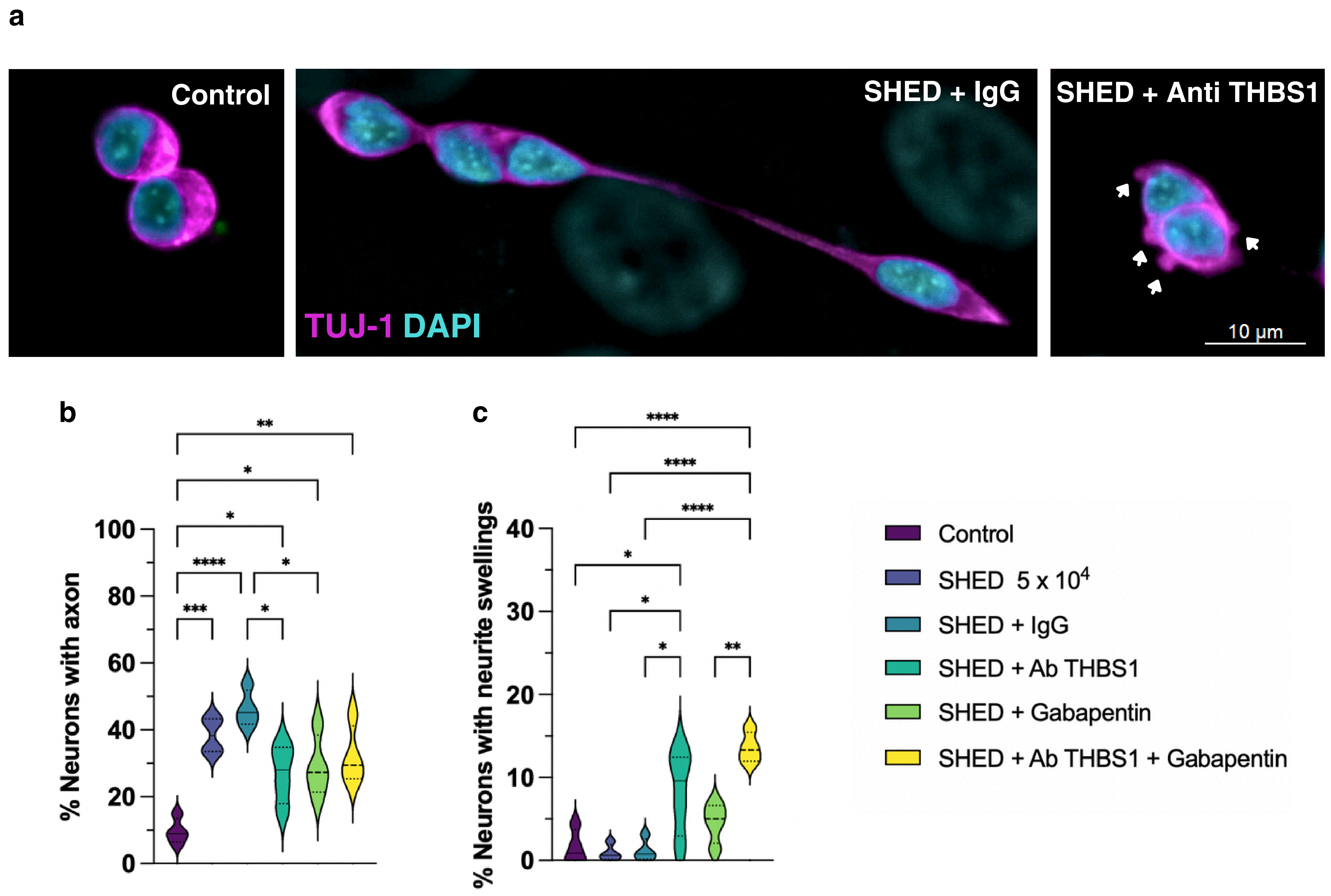

**Supplementary Figure 8**

### Gene Ontology (g:Profiler)

**a**

#### SHED secretome
#### Damage-Induced (24 proteins)

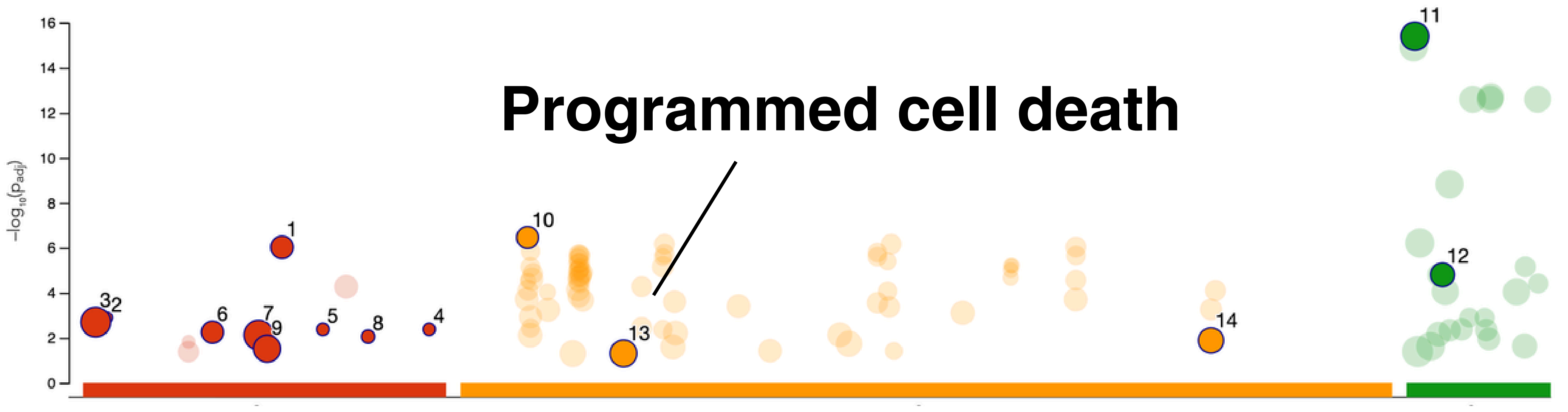


| ID | Source | Term ID | Term Name | $p_{adj}$ (query_... |
|---|---|---|---|---|
| 1 | GO:MF | GO:0045296 | cadherin binding | $9.433\times10^{-7}$ |
| 2 | GO:MF | GO:0004332 | fructose-bisphosphate aldolase activity | $1.262\times10^{-3}$ |
| 3 | GO:MF | GO:0003824 | catalytic activity | $2.000\times10^{-3}$ |
| 4 | GO:MF | GO:0141069 | receptor ligand inhibitor activity | $4.199\times10^{-3}$ |
| 5 | GO:MF | GO:0048030 | disaccharide binding | $4.199\times10^{-3}$ |
| 6 | GO:MF | GO:0030246 | carbohydrate binding | $5.597\times10^{-3}$ |
| 7 | GO:MF | GO:0036094 | small molecule binding | $7.495\times10^{-3}$ |
| 8 | GO:MF | GO:0070061 | fructose binding | $8.806\times10^{-3}$ |
| 9 | GO:MF | GO:0042802 | identical protein binding | $3.049\times10^{-2}$ |
| 10 | GO:BP | GO:0005996 | monosaccharide metabolic process | $3.482\times10^{-7}$ |
| 11 | GO:CC | GO:0005615 | extracellular space | $4.001\times10^{-16}$ |
| 12 | GO:CC | GO:0031012 | extracellular matrix | $1.569\times10^{-5}$ |
| 13 | GO:BP | GO:0012501 | programmed cell death | $4.911\times10^{-2}$ |
| 14 | GO:BP | GO:1901135 | carbohydrate derivative metabolic process | $1.305\times10^{-2}$ |

**b**

#### SHED secretome
#### Damage-Inhibited (30 proteins)

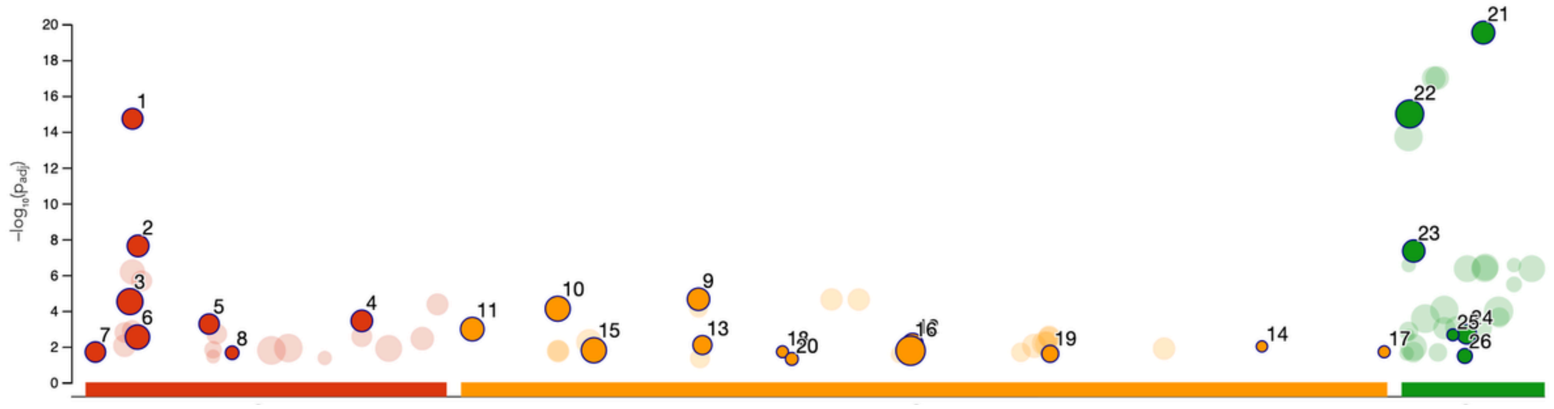


| ID | Source | Term ID | Term Name | $p_{adj}$ (query_... |
|---|---|---|---|---|
| 1 | GO:MF | GO:0005201 | extracellular matrix structural constituent | $1.900\times10^{-15}$ |
| 2 | GO:MF | GO:0005539 | glycosaminoglycan binding | $2.393\times10^{-8}$ |
| 3 | GO:MF | GO:0005102 | signaling receptor binding | $3.091\times10^{-5}$ |
| 4 | GO:MF | GO:0061134 | peptidase regulator activity | $3.670\times10^{-4}$ |
| 5 | GO:MF | GO:0019838 | growth factor binding | $5.527\times10^{-4}$ |
| 6 | GO:MF | GO:0005509 | calcium ion binding | $2.960\times10^{-3}$ |
| 7 | GO:MF | GO:0002020 | protease binding | $2.017\times10^{-2}$ |
| 8 | GO:MF | GO:0031995 | insulin-like growth factor II binding | $2.222\times10^{-2}$ |
| 9 | GO:BP | GO:0030198 | extracellular matrix organization | $2.275\times10^{-5}$ |
| 10 | GO:BP | GO:0007167 | enzyme-linked receptor protein signaling path... | $7.729\times10^{-5}$ |
| 11 | GO:BP | GO:0001501 | skeletal system development | $1.065\times10^{-3}$ |
| 12 | GO:BP | GO:0048592 | eye morphogenesis | $6.955\times10^{-3}$ |
| 13 | GO:BP | GO:0030500 | regulation of bone mineralization | $8.230\times10^{-3}$ |
| 14 | GO:BP | GO:1903225 | negative regulation of endodermal cell differen... | $9.831\times10^{-3}$ |
| 15 | GO:BP | GO:0009887 | animal organ morphogenesis | $1.616\times10^{-2}$ |
| 16 | GO:BP | GO:0048519 | negative regulation of biological process | $1.701\times10^{-2}$ |
| 17 | GO:BP | GO:2001205 | negative regulation of osteoclast development | $1.965\times10^{-2}$ |
| 18 | GO:BP | GO:0035583 | sequestering of TGFbeta in extracellular matrix | $1.965\times10^{-2}$ |
| 19 | GO:BP | GO:0071604 | transforming growth factor beta production | $2.537\times10^{-2}$ |
| 20 | GO:BP | GO:0035989 | tendon development | $4.903\times10^{-2}$ |
| 21 | GO:CC | GO:0062023 | collagen-containing extracellular matrix | $2.929\times10^{-20}$ |
| 22 | GO:CC | GO:0005615 | extracellular space | $1.017\times10^{-15}$ |
| 23 | GO:CC | GO:0005788 | endoplasmic reticulum lumen | $4.622\times10^{-8}$ |
| 24 | GO:CC | GO:0043202 | lysosomal lumen | $2.049\times10^{-3}$ |
| 25 | GO:CC | GO:0034363 | intermediate-density lipoprotein particle | $2.222\times10^{-3}$ |
| 26 | GO:CC | GO:0042627 | chylomicron | $3.342\times10^{-2}$ |

**FBN2**
**FBN1**
**LTF**
**LTBP2**
**ITIH3**
**C7**
**APOB**
**IGFBP4**
**COMP**
**AHSG**
**FBLN1**
**IGFBP2**
**ALB**
**MT2A**
**DCN**
**COL5A1**
**APOC3**
**AOC3**
**RPS23**
**EFEMP1**
**COL5A2**
**LUM**
**TIMP2**
**ITIH1**
**DKK3**
**PLG**
**AFP**
**COL3A1**
**POSTN**
**GC**
