## Supplementary Data 1 for "Human SHED-derived extracellular cues activate a specialized neuroprotective and regenerative program in developing retinal ganglion cells": Axon guidance.pdf

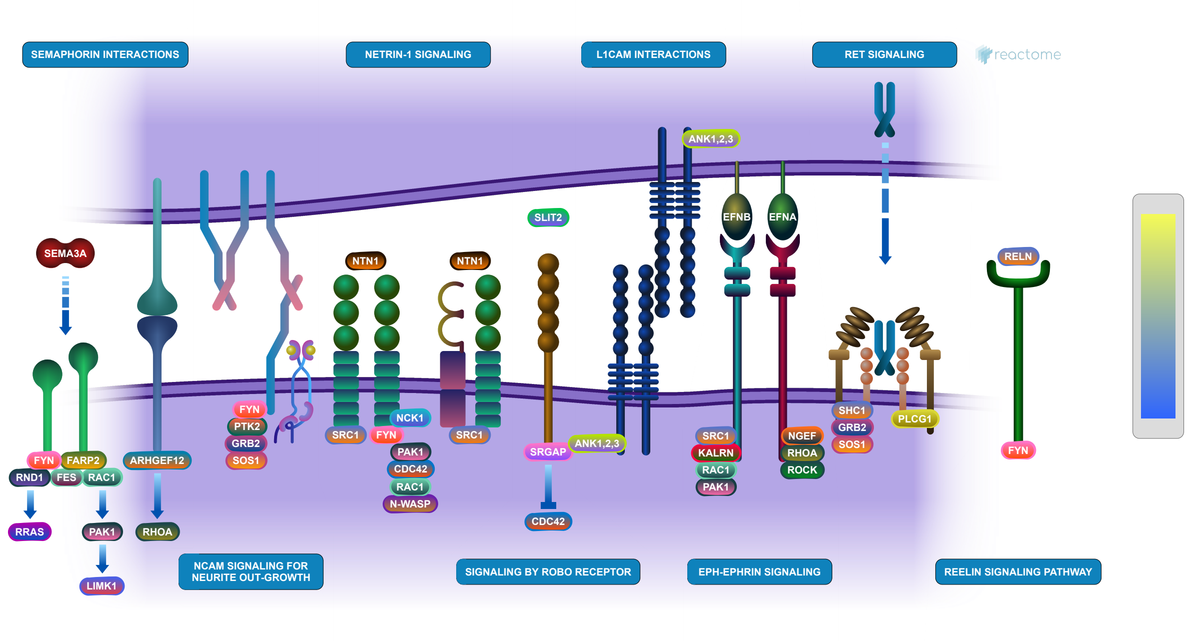


Cooper, HM., D'Eustachio, P., Garapati, P V., Ip, NY., Jassal, B., Jaworski, A., Jupe, S., Kidd, T., Kikutani, H., Kumanogoh, A., Luo, W., Maness, PF., Morales, D., Orlic-Milacic, M., Walmod, PS.

European Bioinformatics Institute, New York University Langone Medical Center, Ontario Institute for Cancer Research, Oregon Health and Science University.

The contents of this document may be freely copied and distributed in any media, provided the authors, plus the institutions, are credited, as stated under the terms of Creative Commons Attribution 4.0 International (CC BY 4.0) License. For more information see our license.

This is just an excerpt of a full-length report for this pathway. To access the complete report, please download it at the Reactome Textbook.

03/06/2026

### Introduction

Reactome is open-source, open access, manually curated and peer-reviewed pathway database. Pathway annotations are authored by expert biologists, in collaboration with Reactome editorial staff and cross-referenced to many bioinformatics databases. A system of evidence tracking ensures that all assertions are backed up by the primary literature. Reactome is used by clinicians, geneticists, genomics researchers, and molecular biologists to interpret the results of high-throughput experimental studies, by bioinformaticians seeking to develop novel algorithms for mining knowledge from genomic studies, and by systems biologists building predictive models of normal and disease variant pathways.

The development of Reactome is supported by grants from the US National Institutes of Health (P41 HG003751), University of Toronto (CFREF Medicine by Design), European Union (EU STRP, EMI-CD), and the European Molecular Biology Laboratory (EBI Industry program).

#### Literature references

Fabregat, A., Fabregat, A., Fabregat, A., Fabregat, A., Fabregat, A., Fabregat, A. et al. (2017). Reactome pathway analysis: a high-performance in-memory approach. *BMC bioinformatics, 18*, 142. ↗

Sidiropoulos, K., Sidiropoulos, K., Sidiropoulos, K., Sidiropoulos, K., Sidiropoulos, K., Sidiropoulos, K. et al. (2017). Reactome enhanced pathway visualization. *Bioinformatics, 33*, 3461-3467. ↗

Fabregat, A., Fabregat, A., Fabregat, A., Fabregat, A., Fabregat, A., Fabregat, A. et al. (2018). The Reactome Pathway Knowledgebase. *Nucleic Acids Res, 46*, D649-D655. ↗

Fabregat, A., Fabregat, A., Fabregat, A., Fabregat, A., Fabregat, A., Fabregat, A. et al. (2018). Reactome graph database: Efficient access to complex pathway data. *PLoS computational biology, 14*, e1005968. ↗

Reactome database release: 96

This document contains 9 pathways (see Table of Contents)

### Analysis properties

- This is an **expression** analysis: The numbers are used to produce a scaled coloured overlay over Reactome pathway diagrams, as a means to visualize relative expression levels. Note that the numeric values do not have to be expression data, for instance by using gene association scores the same analysis can be used to visualize genotyping results.
  See more
- 175 out of 192 identifiers in the sample were found in Reactome, where 1054 pathways were hit by at least one of them.
- All non-human identifiers have been converted to their human equivalent. ↗
- This report is filtered to show only results for species 'Homo sapiens' and resource 'all resources'.
- The unique ID for this analysis (token) is MjAyNjA2MDMxNDU1NDNfOTEzOA%3D%3D. This ID is valid for at least 7 days in Reactome's server. Use it to access Reactome services with your data.

### Axon guidance ↗

**Stable identifier:** R-HSA-422475

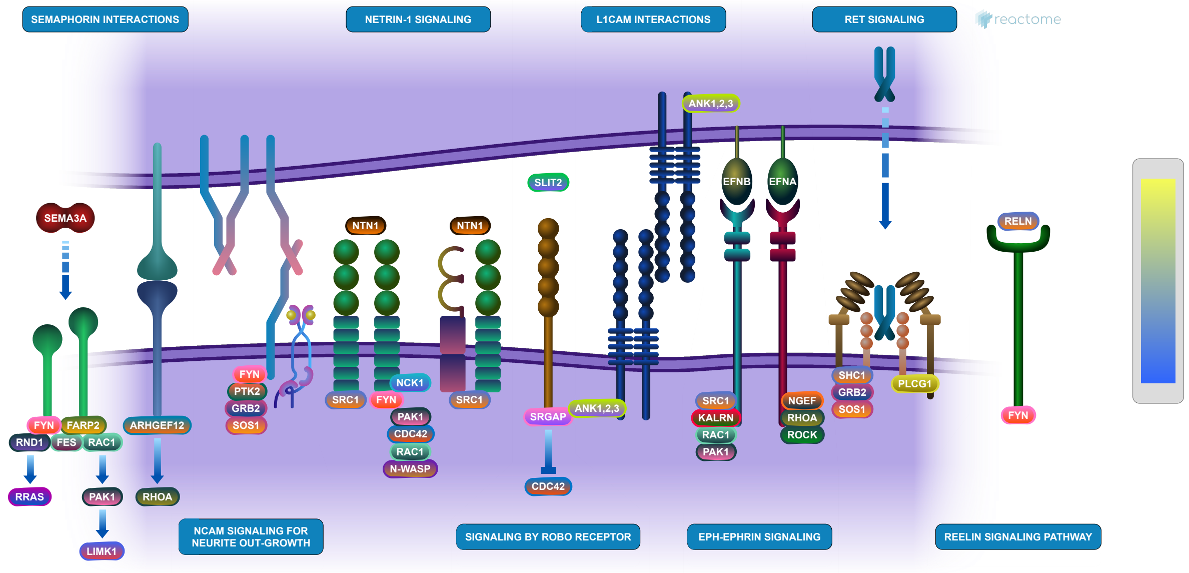


Axon guidance / axon pathfinding is the process by which neurons send out axons to reach the correct targets. Growing axons have a highly motile structure at the growing tip called the growth cone, which senses the guidance cues in the environment through guidance cue receptors and responds by undergoing cytoskeletal changes that determine the direction of axon growth.
Guidance cues present in the surrounding environment provide the necessary directional information for the trip. These extrinsic cues have been divided into attractive or repulsive signals that tell the growth cone where and where not to grow.
Genetic and biochemical studies have led to the identification of highly conserved families of guidance molecules and their receptors that guide axons. These include netrins, Slits, semaphorins, and ephrins, and their cognate receptors, DCC and or uncoordinated-5 (UNC5), roundabouts (Robo), neuropilin and Eph. In addition, many other classes of adhesion molecules are also used by growth cones to navigate properly which include NCAM and L1CAM.
For review of axon guidance, please refer to Russel and Bashaw 2018, Chedotal 2019, Suter and Jaworski 2019).
Axon guidance cues and their receptors are implicated in cancer progression (Biankin et al. 2012), where they likely contribute to cell migration and angiogenesis (reviewed by Mehlen et al. 2011).

#### Literature references

Chédotal, A. (2019). Roles of axon guidance molecules in neuronal wiring in the developing spinal cord. *Nat. Rev. Neurosci., 20*, 380-396. ↗

Suter, TACS., Jaworski, A. (2019). Cell migration and axon guidance at the border between central and peripheral nervous system. *Science, 365*. ↗

Russell, SA., Bashaw, GJ. (2018). Axon guidance pathways and the control of gene expression. *Dev. Dyn., 247*, 571-580 . ↗

#### Editions

| | | |
|---|---|---|
| 2009-05-26 | Reviewed | Maness, PF., Walmod, PS. |
| 2009-05-28 | Authored, Edited | Garapati, P V. |

#### 27 submitted entities found in this pathway, mapping to 33 Reactome entities

| Input | UniProt Id | col1 |
|---|---|---|
| COL6A3 | A8TX70, P12111 | 11 |
| HSPA8 | P11142 | 12 |

| Input | UniProt Id | col1 |
|---|---|---|
| COL6A1 | P12109 | 17 |
| COL6A2 | P12110 | 2 |
| COL4A1 | P02462 | 3 |
| MMP2 | P08253 | 26 |
| LAMB1 | P07942 | 2 |
| COL4A2 | P08572 | 4 |
| GPC1 | P35052 | 9 |
| ACTB | P60709, P63261 | 54 |
| TUBB4B | P04350, P68371 | 3 |
| COL5A1 | P20908 | 5 |
| COL5A2 | P05997 | 1 |
| COL3A1 | P02461 | 12 |
| CFL1 | P23528 | 20 |
| MSN | P26038 | 4 |
| HSP90AB1 | P08238 | 10 |
| AGRN | O00468 | 3 |
| PFN1 | P07737 | 17 |
| MYH9 | P35579 | 4 |
| RPS23 | P62266 | 8 |
| TUBA1A | Q71U36 | 14 |
| RPL35 | P42766 | 10 |
| UBB | P0CG47, P62979, P62987 | 33 |
| RPL31 | P62899 | 21 |
| NRP2 | O60462, Q99435 | 2 |
| TLN1 | Q9Y490 | 8 |

#### Semaphorin interactions ↗

**Location:** Axon guidance

**Stable identifier:** R-HSA-373755

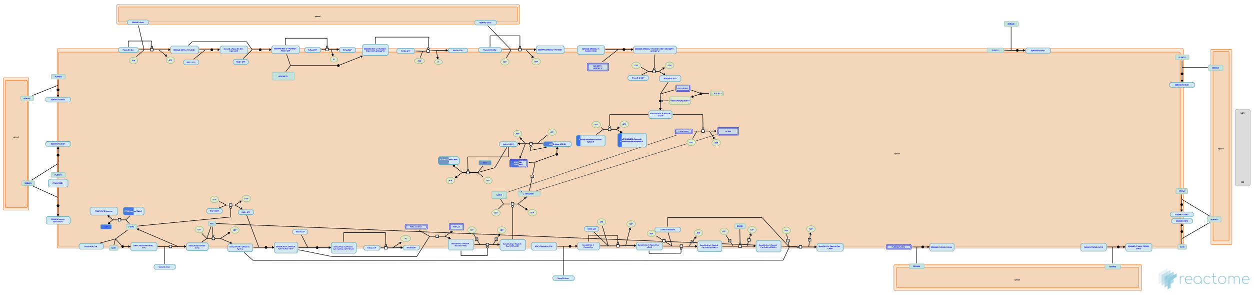

Semaphorins are a large family of cell surface and secreted guidance molecules divided into eight classes on the basis of their structures. They all have an N-terminal conserved sema domain. Semaphorins signal through multimeric receptor complexes that include other proteins such as plexins and neuropilins.

#### Literature references


Dickson, BJ. (2002). Molecular mechanisms of axon guidance. *Science, 298*, 1959-64. ↗

Pasterkamp, RJ., Kolodkin, AL. (2003). Semaphorin junction: making tracks toward neural connectivity. *Curr Opin Neurobiol, 13*, 79-89. ↗

Zhou, Y., Gunput, RA., Pasterkamp, RJ. (2008). Semaphorin signaling: progress made and promises ahead. *Trends Biochem Sci, 33*, 161-70. ↗

Pasterkamp, RJ., Verhaagen, J. (2006). Semaphorins in axon regeneration: developmental guidance molecules gone wrong?. *Philos Trans R Soc Lond B Biol Sci, 361*, 1499-511. ↗

Koncina, E., Roth, L., Gonthier, B., Bagnard, D. (2007). Role of semaphorins during axon growth and guidance. *Adv Exp Med Biol, 621*, 50-64. ↗


#### Editions

| | | |
|---|---|---|
| 2009-03-23 | Authored, Edited | Garapati, P V. |
| 2009-09-02 | Reviewed | Kikutani, H., Kumanogoh, A. |

#### 4 submitted entities found in this pathway, mapping to 4 Reactome entities

| Input | UniProt Id | col1 |
|---|---|---|
| MYH9 | P35579 | 4 |
| CFL1 | P23528 | 20 |
| TLN1 | Q9Y490 | 8 |
| HSP90AB1 | P08238 | 10 |

### NCAM signaling for neurite out-growth ↗

**Location:** Axon guidance

**Stable identifier:** R-HSA-375165

**Compartments:** plasma membrane

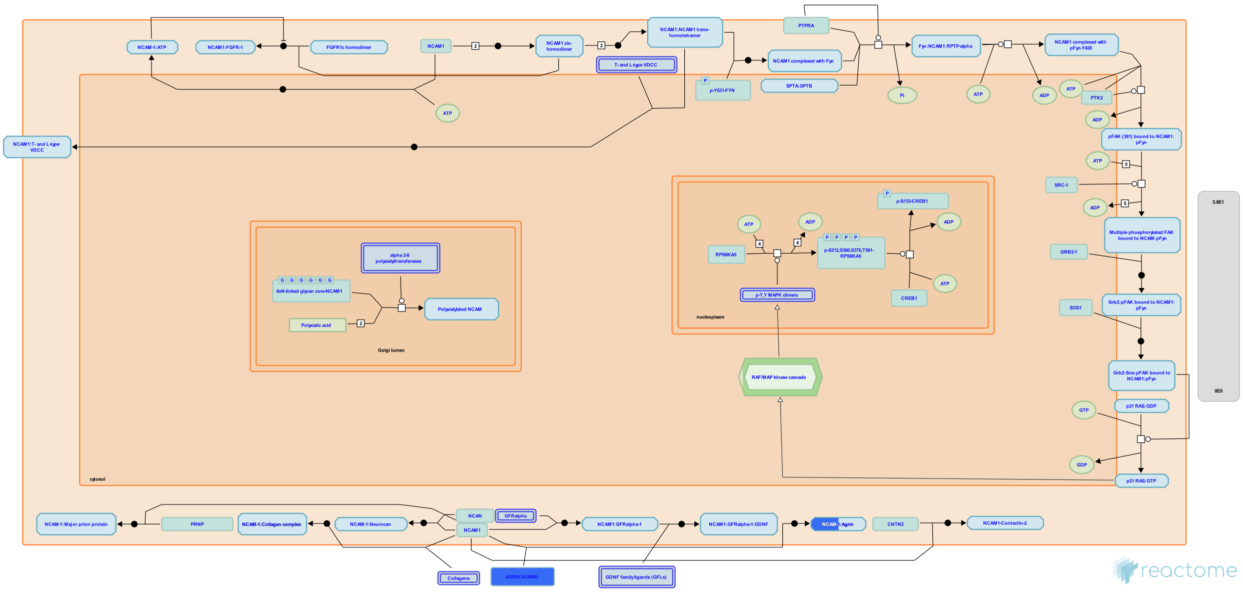

The neural cell adhesion molecule, NCAM, is a member of the immunoglobulin (Ig) superfamily and is involved in a variety of cellular processes of importance for the formation and maintenance of the nervous system. The role of NCAM in neural differentiation and synaptic plasticity is presumed to depend on the modulation of intracellular signal transduction cascades. NCAM based signaling complexes can initiate downstream intracellular signals by at least two mechanisms: (1) activation of FGFR and (2) formation of intracellular signaling complexes by direct interaction with cytoplasmic interaction partners such as Fyn and FAK. Tyrosine kinases Fyn and FAK interact with NCAM and undergo phosphorylation and this transiently activates the MAPK, ERK 1 and 2, cAMP response element binding protein (CREB) and transcription factors ELK and NFkB. CREB activates transcription of genes which are important for axonal growth, survival, and synaptic plasticity in neurons.

NCAM1 mediated intracellular signal transduction is represented in the figure below. The Ig domains in NCAM1 are represented in orange ovals and Fn domains in green squares. The tyrosine residues susceptible to phosphorylation are represented in red circles and their positions are numbered. Phosphorylation is represented by red arrows and dephosphorylation by yellow. Ig, Immunoglobulin domain; Fn, Fibronectin domain; Fyn, Proto-oncogene tyrosine-protein kinase Fyn; FAK, focal adhesion kinase; RPTPalpha, Receptor-type tyrosine-protein phosphatase; Grb2, Growth factor receptor-bound protein 2; SOS, Son of sevenless homolog; Raf, RAF proto-oncogene serine/threonine-protein kinase; MEK, MAPK and ERK kinase; ERK, Extracellular signal-regulated kinase; MSK1, Mitogen and stress activated protein kinase 1; CREB, Cyclic AMP-responsive element-binding protein; CRE, cAMP response elements.

#### Literature references


Walmod, PS., Kolkova, K., Berezin, V., Bock, E. (2004). Zippers make signals: NCAM-mediated molecular interactions and signal transduction. *Neurochem Res, 29*, 2015-35. ↗

Ditlevsen, DK., Povlsen, GK., Berezin, V., Bock, E. (2008). NCAM-induced intracellular signaling revisited. *J Neurosci Res, 86*, 727-43. ↗

Schmid, RS., Maness, PF. (2008). L1 and NCAM adhesion molecules as signaling coreceptors in neuronal migration and process outgrowth. *Curr Opin Neurobiol, 18*, 245-50. ↗

Panicker, AK., Buhusi, M., Thelen, K., Maness, PF. (2003). Cellular signalling mechanisms of neural cell adhesion molecules. *Front Biosci, 8*, d900-11. ↗

Kiryushko, D., Berezin, V., Bock, E. (2004). Regulators of neurite outgrowth: role of cell adhesion molecules. *Ann N Y Acad Sci, 1014*, 140-54. ↗

#### Editions

| | | |
|---|---|---|
| 2009-02-24 | Authored, Edited | Garapati, P V. |
| 2009-05-26 | Reviewed | Maness, PF., Walmod, PS. |

#### 9 submitted entities found in this pathway, mapping to 10 Reactome entities

| Input | UniProt Id | col1 |
|---|---|---|
| COL6A3 | A8TX70, P12111 | 11 |
| COL5A2 | P05997 | 1 |
| COL6A1 | P12109 | 17 |
| COL6A2 | P12110 | 2 |
| COL4A1 | P02462 | 3 |
| COL4A2 | P08572 | 4 |
| COL5A1 | P20908 | 5 |
| COL3A1 | P02461 | 12 |
| AGRN | O00468 | 3 |

### Netrin-1 signaling ↗

**Location:** Axon guidance

**Stable identifier:** R-HSA-373752

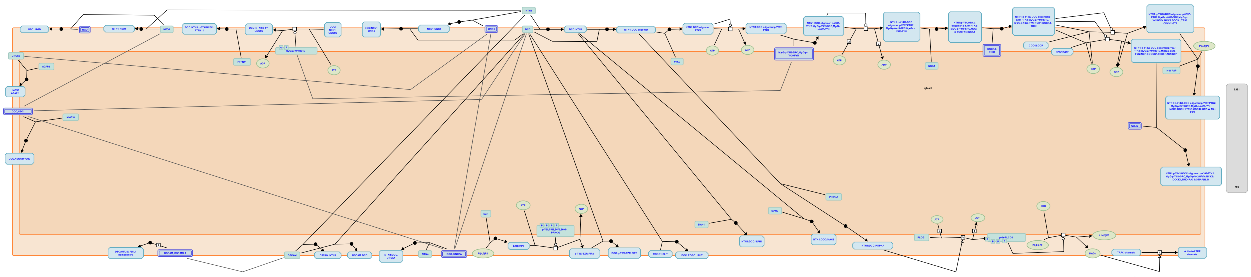

Netrins are secreted proteins that play a crucial role in neuronal migration and in axon guidance during the development of the nervous system. To date, several Netrins have been described in mouse and humans: Netrin-1, -3/NTL2, -4/h and G-Netrins. Netrin-1 is the most studied member of the family and has been shown to play a crucial role in neuronal navigation during nervous system development mainly through its interaction with its receptors DCC and UNC5. Members of the Deleted in colorectal cancer (DCC) family- which includes DCC and Neogenin in vertebrates- mediate netrin-induced axon attraction, whereas the C. elegans UNC5 receptor and its four vertebrate homologs Unc5a-Unc5d mediate repulsion.

#### Literature references

Moore, SW., Tessier-Lavigne, M., Kennedy, TE. (2007). Netrins and their receptors. *Adv Exp Med Biol, 621*, 17-31. ↗

Cooper, HM., Gad, JM., Keeling, SL. (1999). The Deleted in Colorectal Cancer netrin guidance system: a molecular strategy for neuronal navigation. *Clin Exp Pharmacol Physiol, 26*, 749-51. ↗

#### Editions

| | | |
|---|---|---|
| 2008-07-16 | Authored | Garapati, P V. |
| 2008-07-30 | Edited | Garapati, P V. |
| 2010-02-16 | Reviewed | Cooper, HM. |

#### Signaling by ROBO receptors ↗

**Location:** Axon guidance

**Stable identifier:** R-HSA-376176

**Compartments:** plasma membrane

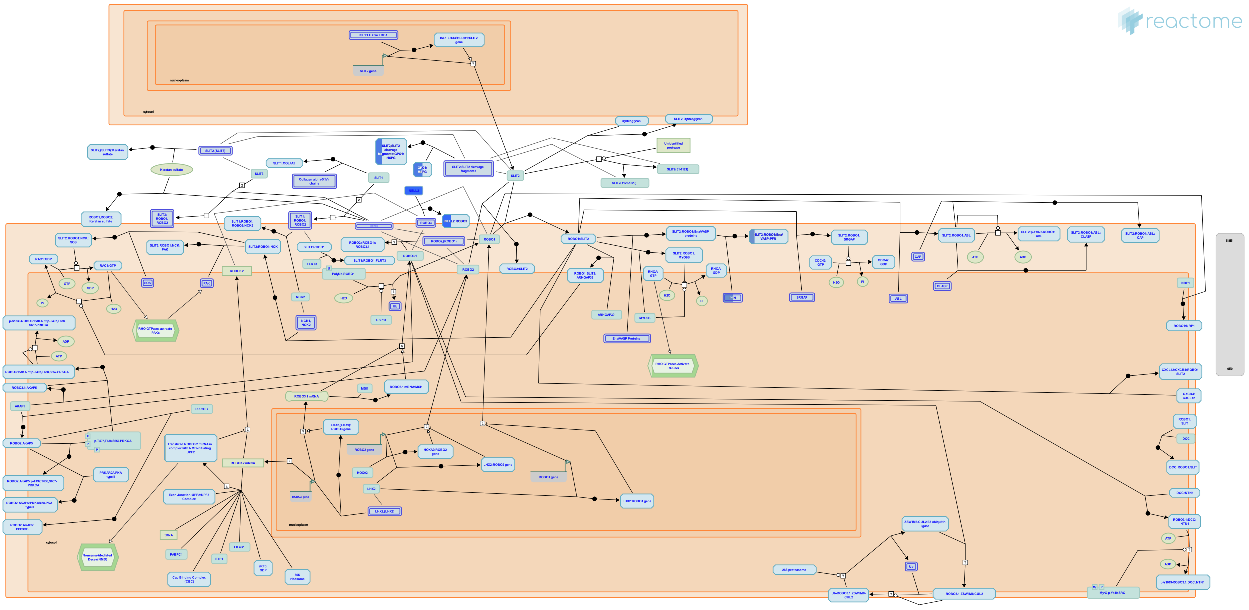

The Roundabout (ROBO) family encodes transmembrane receptors that regulate axonal guidance and cell migration. The major function of the Robo receptors is to mediate repulsion of the navigating growth cones. There are four human Robo homologues, ROBO1, ROBO2, ROBO3 and ROBO4. Most of the ROBOs have the similar ectodomain architecture as the cell adhesion molecules, with five Ig domains followed by three FN3 repeats, except for ROBO4. ROBO4 has two Ig and two FN3 repeats. The cytoplasmic domains of ROBO receptors are in general poorly conserved. However, there are four short conserved cytoplasmic sequence motifs, named CC0-3, that serve as binding sites for adaptor proteins. The ligands for the human ROBO1 and ROBO2 receptors are the three SLIT proteins SLIT1, SLIT2, and SLIT3; all of the SLIT proteins contain a tandem of four LRR (leucine rich repeat) domains at the N-terminus, termed D1-D4, followed by six EGF (epidermal growth factor)-like domains, a laminin G like domain (ALPS), three EGF-like domains, and a C-terminal cysteine knot domain. Most SLIT proteins are cleaved within the EGF-like region by unknown proteases (reviewed by Hohenster 2008, Ypsilanti and Chedotal 2014, Blockus and Chedotal 2016). NELL2 is a ligand for ROBO3 (Jaworski et al. 2015).

SLIT protein binding modulates ROBO interactions with the cytosolic adaptors. The cytoplasmic domain of ROBO1 and ROBO2 determines the repulsive responses of these receptors. Based on the studies from both invertebrate and vertebrate organisms it has been inferred that ROBO induces growth cone repulsion by controlling cytoskeletal dynamics via either Abelson kinase (ABL) and Enabled (Ena), or RAC1 activity (reviewed by Hohenster 2008, Ypsilanti and Chedotal 2014, Blockus and Chedotal 2016). While there is some redundancy in the function of ROBO receptors, ROBO1 is implicated as the predominant receptor for axon guidance in ventral tracts, and ROBO2 is the predominant receptor for axon guidance in dorsal tracts. ROBO2 also repels neuron cell bodies from the floor plate (Kim et al. 2011).

In addition to regulating axon guidance, ROBO1 and ROBO2 receptors are also implicated in regulation of proliferation and transition of primary to intermediate neuronal progenitors through a poorly characterized cross-talk with NOTCH-mediated activation of HES1 transcription (Borrell et al. 2012).

Thalamocortical axon extension is regulated by neuronal activity-dependent transcriptional regulation of ROBO1 transcription. Lower neuronal activity correlates with increased ROBO1 transcription, possibly mediated by the NFKB complex (Mire et al. 2012).

It is suggested that the homeodomain transcription factor NKX2.9 stimulates transcription of ROBO2, which is involved in regulation of motor axon exit from the vertebrate spinal code (Bravo-Ambrosio et al. 2012).

Of the four ROBO proteins, ROBO4 is not involved in neuronal system development but is, instead, involved in angiogenesis. The interaction of ROBO4 with SLIT3 is involved in proliferation, motility and chemotaxis of endothelial cells, and accelerates formation of blood vessels (Zhang et al. 2009).

#### Literature references


Hohenester, E. (2008). Structural insight into Slit-Robo signalling. *Biochem Soc Trans, 36*, 251-6. ↗

Borrell, V., Cárdenas, A., Ciceri, G., Galcerán, J., Flames, N., Pla, R. et al. (2012). Slit/Robo signaling modulates the proliferation of central nervous system progenitors. *Neuron, 76*, 338-52. ↗

Mire, E., Mezzera, C., Leyva-Díaz, E., Paternain, AV., Squarzoni, P., Bluy, L. et al. (2012). Spontaneous activity regulates Robo1 transcription to mediate a switch in thalamocortical axon growth. *Nat. Neurosci., 15*, 1134-43. ↗

Bravo-Ambrosio, A., Mastick, G., Kaprielian, Z. (2012). Motor axon exit from the mammalian spinal cord is controlled by the homeodomain protein Nkx2.9 via Robo-Slit signaling. *Development, 139*, 1435-46. ↗

Zhang, B., Dietrich, UM., Geng, JG., Bicknell, R., Esko, JD., Wang, L. (2009). Repulsive axon guidance molecule Slit3 is a novel angiogenic factor. *Blood, 114*, 4300-9. ↗


#### Editions

| | | |
|---|---|---|
| 2008-09-05 | Authored, Edited | Garapati, P V. |
| 2009-08-18 | Reviewed | Kidd, T. |
| 2017-06-23 | Edited, Revised | Orlic-Milacic, M. |
| 2017-07-31 | Reviewed | Jaworski, A. |
| 2017-08-04 | Edited | Orlic-Milacic, M. |

#### 7 submitted entities found in this pathway, mapping to 9 Reactome entities

| Input | UniProt Id | col1 |
|---|---|---|
| UBB | P0CG47, P62979, P62987 | 33 |
| GPC1 | P35052 | 9 |
| PFN1 | P07737 | 17 |
| RPS23 | P62266 | 8 |
| RPL35 | P42766 | 10 |
| RPL31 | P62899 | 21 |
| NRP2 | Q99435 | 2 |

#### L1CAM interactions ↗

**Location:** Axon guidance

**Stable identifier:** R-HSA-373760

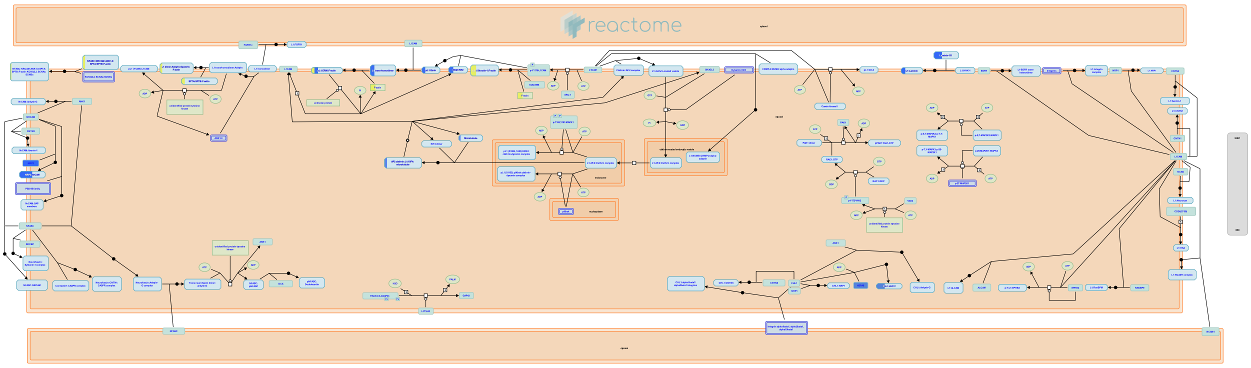

The L1 family of cell adhesion molecules (L1CAMs) are a subfamily of the immunoglobulin superfamily of transmembrane receptors, comprised of four structurally related proteins: L1, Close Homolog of L1 (CHL1), NrCAM, and Neurofascin. These CAMs contain six Ig like domains, five or six fibronectin like repeats, a transmembrane region and a cytoplasmic domain. The L1CAM family has been implicated in processes integral to nervous system development, including neurite outgrowth, neurite fasciculation and inter neuronal adhesion.
L1CAM members are predominately expressed by neuronal, as well as some nonneuronal cells, during development. Except CHL1 all the other members of L1 family contain an alternatively spliced 12-ncleuotide exon, encoding the amino acid residues RSLE in the neuronal splice forms but missing in the non-neuronal cells. The extracellular regions of L1CAM members are divergent and differ in their abilities to interact with extracellular, heterophilic ligands. The L1 ligands include other Ig-domain CAMs (such as NCAM, TAG-1/axonin and F11), proteoglycans type molecules (neurocan), beta1 integrins, and extra cellular matrix protein laminin, Neuropilin-1, FGF and EGF receptors. Some of these L1-interacting proteins also bind to other L1CAM members. For example TAG-1/axonin interact with L1 and NrCAM; L1, neurofascin and CHL1 binds to contactin family members. The cytoplasmic domains of L1CAM members are most highly conserved. Nevertheless, they have different cytoplasmic binding partners, and even those with similar binding partners may be involved in different signaling complexes and mechanisms. The most conserved feature of L1CAMs is their ability to interact with the actin cytoskeletal adapter protein ankyrin. The cytoplasmic ankyrin-binding domain, exhibits the highest degree of amino acid conservation throughout the L1 family.

##### Literature references


Kamiguchi, H. (2003). The mechanism of axon growth: what we have learned from the cell adhesion molecule L1. *Mol Neurobiol, 28*, 219-28. ↗

Kamiguchi, H., Lemmon, V. (1997). Neural cell adhesion molecule L1: signaling pathways and growth cone motility. *J Neurosci Res, 49*, 1-8. ↗

Herron, LR., Hill, M., Davey, F., Gunn-Moore, FJ. (2009). The intracellular interactions of the L1 family of cell adhesion molecules. *Biochem J, 419*, 519-31. ↗

Schmid, RS., Maness, PF. (2008). L1 and NCAM adhesion molecules as signaling coreceptors in neuronal migration and process outgrowth. *Curr Opin Neurobiol, 18*, 245-50. ↗

Maness, PF., Schachner, M. (2007). Neural recognition molecules of the immunoglobulin superfamily: signaling transducers of axon guidance and neuronal migration. *Nat Neurosci, 10*, 19-26. ↗


##### Editions

| | | |
|---|---|---|
| 2008-07-30 | Authored, Edited | Garapati, P V. |
| 2010-02-16 | Reviewed | Maness, PF. |

##### 7 submitted entities found in this pathway, mapping to 9 Reactome entities

| Input | UniProt Id | col1 |
|---|---|---|
| LAMB1 | P07942 | 2 |
| NRP2 | O60462 | 2 |

| Input | UniProt Id | col1 |
|---|---|---|
| HSPA8 | P11142 | 12 |
| MSN | P26038 | 4 |
| ACTB | P60709, P63261 | 54 |
| TUBB4B | P04350, P68371 | 3 |
| TUBA1A | Q71U36 | 14 |

### EPH-Ephrin signaling ↗

**Location:** Axon guidance

**Stable identifier:** R-HSA-2682334

**Compartments:** cytosol, plasma membrane

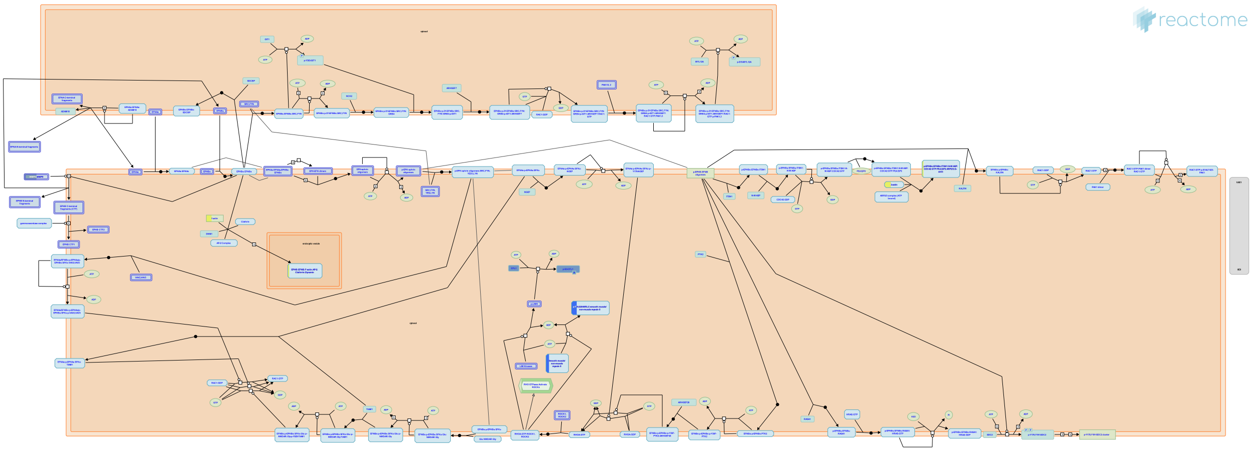

During the development process cell migration and adhesion are the main forces involved in morphing the cells into critical anatomical structures. The ability of a cell to migrate to its correct destination depends heavily on signaling at the cell membrane. Erythropoietin producing hepatocellular carcinoma (EPH) receptors and their ligands, the ephrins (EPH receptors interacting proteins, EFNs), orchestrates the precise control necessary to guide a cell to its destination. They are expressed in all tissues of a developing embryo and are involved in multiple developmental processes such as axon guidance, cardiovascular and skeletal development and tissue patterning. In addition, EPH receptors and EFNs are expressed in developing and mature synapses in the nervous system, where they may have a role in regulating synaptic plasticity and long-term potentiation. Activation of EPHB receptors in neurons induces the rapid formation and enlargement of dendritic spines, as well as rapid synapse maturation (Dalva et al. 2007). On the other hand, EPHA4 activation leads to dendritic spine elimination (Murai et al. 2003, Fu et al. 2007).
EPH receptors are the largest known family of receptor tyrosine kinases (RTKs), with fourteen total receptors divided into either A- or B-subclasses: EPHA (1-8 and 10) and EPHB (1-4 and 6). EPH receptors can have overlapping functions, and loss of one receptor can be partially compensated for by another EPH receptor that has similar expression pattern and ligand-binding specificities. EPH receptors have an N-terminal extracellular domain through which they bind to ephrin ligands, a short transmembrane domain, and an intracellular cytoplasmic signaling structure containing a canonical tyrosine kinase catalytic domain as well as other protein interaction sites. Ephrins are also sub-divided into an A-subclass (A1-A5), which are tethered to the plasma membrane by a glycosylphosphatidylinositol (GPI) anchor, and a B-subclass (B1-B3), members of which have a transmembrane domain and a short, highly conserved cytoplasmic tail lacking endogenous catalytic activity. The interaction between EPH receptors and its ligands requires cell-cell interaction since both molecules are membrane-bound. Close contact between EPH receptors and EFNs is required for signaling to occur. EPH/EFN-initiated signaling occurs bi-directionally into either EPH- or EFN-expressing cells or axons. Signaling into the EPH receptor-expressing cell is referred as the forward signal and signaling into the EFN-expressing cell, the reverse signal. (Dalva et al. 2000, Grunwald et al. 2004, Davy & Robbins 2000, Cowan et al. 2004)

#### Literature references


Arvanitis, D., Davy, A. (2008). Eph/ephrin signaling: networks. *Genes Dev.*, *22*, 416-29. ↗

Chen, Y., Fu, AK., Ip, NY. (2012). Eph receptors at synapses: implications in neurodegenerative diseases. *Cell. Signal.*, *24*, 606-11. ↗


#### Editions

| | | |
|---|---|---|
| 2013-07-23 | Authored, Edited | Garapati, P V. |
| 2014-05-19 | Reviewed | Ip, NY. |

#### 4 submitted entities found in this pathway, mapping to 5 Reactome entities

| Input | UniProt Id | col1 |
|---|---|---|
| MYH9 | P35579 | 4 |

| Input | UniProt Id | col1 |
|---|---|---|
| CFL1 | P23528 | 20 |
| MMP2 | P08253 | 26 |
| ACTB | P60709, P63261 | 54 |

### RET signaling ↗

**Location:** Axon guidance

**Stable identifier:** R-HSA-8853659

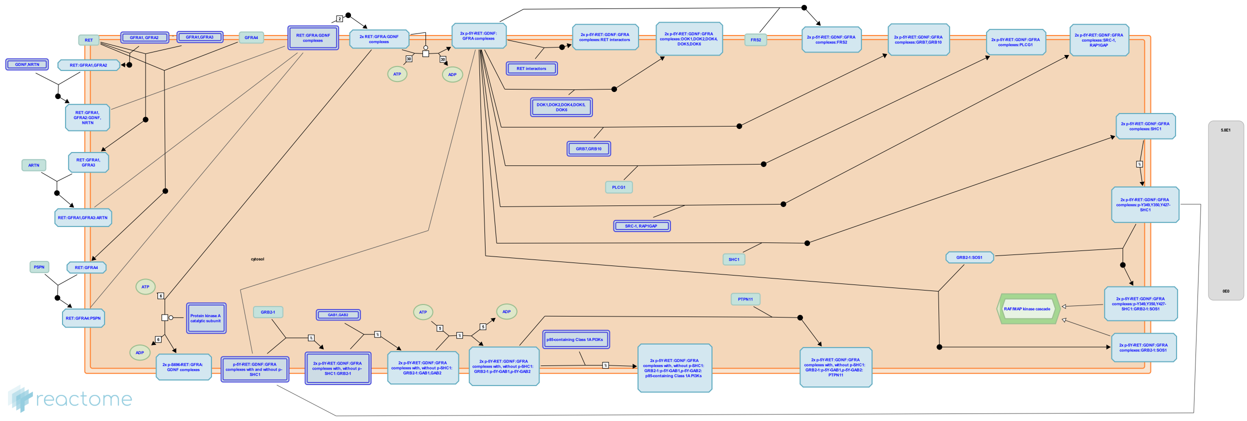

The RET proto-oncogene encodes a receptor tyrosine kinase expressed primarily in urogenital precursor cells, spermatogonocytes, dopaminergic neurons, motor neurons and neural crest progenitors and derived cells. . It is essential for kidney genesis, spermatogonial self-renewal and survivial, specification, migration, axonal growth and axon guidance of developing enteric neurons, motor neurons, parasympathetic neurons and somatosensory neurons (Schuchardt et al. 1994, Enomoto et al. 2001, Naughton et al. 2006, Kramer et al. 2006, Luo et al. 2006, 2009). RET was identified as the causative gene for human papillary thyroid carcinoma (Grieco et al. 1990), multiple endocrine neoplasia (MEN) type 2A (Mulligan et al. 1993), type 2B (Hofstra et al. 1994, Carlson et al. 1994), and Hirschsprung's disease (Romeo et al. 1994, Edery et al. 1994).

RET contains a cadherin-related motif and a cysteine-rich domain in the extracellular domain (Takahashi et al. 1988). It is the receptor for members of the glial cell-derived neurotrophic factor (GDNF) family of ligands, GDNF (Lin et al. 1993), neurturin (NRTN) (Kotzbauer et al. 1996), artemin (ARTN) (Baloh et al. 1998), and persephin (PSPN) (Milbrandt et al. 1998), which form a family of neurotrophic factors. To stimulate RET, these ligands need a glycosylphosphatidylinositol (GPI)-anchored co-receptor, collectively termed GDNF family receptor-alpha (GFRA) (Treanor et al. 1996, Jing et al. 1996). The four members of this family have different, overlapping ligand preferences. GFRA1, GFRA2, GFRA3, and GFRA4 preferentially bind GDNF, NRTN, ARTN and PSPN, respectively (Jing et al. 1996, 1997, Creedon et al. 1997, Baloh et al. 1997, 1998, Masure et al. 2000). The GFRA co-receptor can come from the same cell as RET, or from a different cell. When the co-receptor is produced by the same cell as RET, it is termed cis signaling. When the co-receptor is produced by another cell, it is termed trans signaling. Cis and trans activation has been proposed to diversify RET signaling, either by recruiting different downstream effectors or by changing the kinetics or efficacy of kinase activation (Tansey et al. 2000, Paratcha et al. 2001). Whether cis and trans signaling has significant differences in vivo is unresolved (Fleming et al. 2015). Different GDNF family members could activate similar downstream signaling pathways since all GFRAs bind to and activate the same tyrosine kinase and induce coordinated phosphorylation of the same four RET tyrosines (Tyr905, Tyr1015, Tyr1062, and Tyr1096) with similar kinetics (Coulpier et al. 2002). However the exact RET signaling pathways in different types of cells and neurons remain to be determined.

#### Literature references


Ichihara, M., Murakumo, Y., Takahashi, M. (2004). RET and neuroendocrine tumors. *Cancer Lett.*, *204*, 197-211. ↗

Murakumo, Y., Jijiwa, M., Asai, N., Ichihara, M., Takahashi, M. (2006). RET and neuroendocrine tumors. *Pituitary*, *9*, 179-92. ↗


#### Editions

| | | |
|---|---|---|
| 2016-01-25 | Authored | Jupe, S. |
| 2016-04-28 | Edited | Jupe, S. |
| 2016-05-06 | Reviewed | Morales, D. |
| 2016-05-17 | Reviewed | Luo, W. |

#### Reelin signalling pathway ↗

**Location:** Axon guidance

**Stable identifier:** R-HSA-8866376

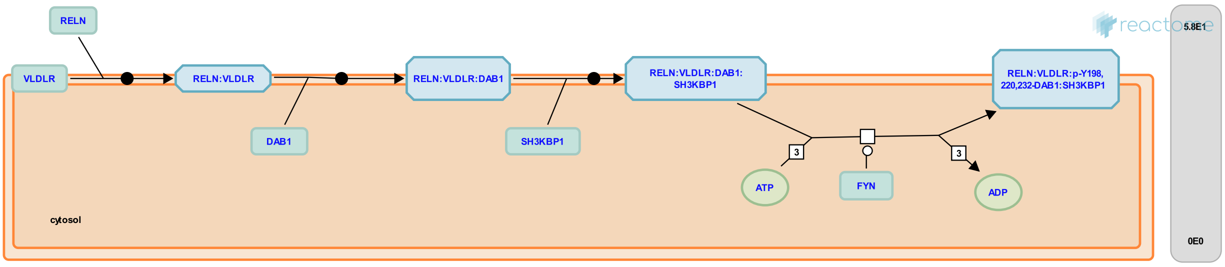


Reelin (RELN) is an extracellular, multifunctional signal glycoprotein that controls not only the positioning of neurons in the developing brain, but also their growth, maturation, and synaptic activity in the adult brain (Stranahan et al. 2013). Abnormal Reelin expression in the brain is implicated in a number of neuropsychiatric disorders including autism, schizophrenia, bipolar disorder and Alzheimer's disease (Folsom & Fatemi 2013).

#### Literature references


Stranahan, AM., Erion, JR., Wosiski-Kuhn, M. (2013). Reelin signaling in development, maintenance, and plasticity of neural networks. *Ageing Res. Rev., 12*, 815-22. ↗

Folsom, TD., Fatemi, SH. (2013). The involvement of Reelin in neurodevelopmental disorders. *Neuropharmacology, 68*, 122-35. ↗


#### Editions

| | | |
|---|---|---|
| 2016-03-31 | Authored, Edited | Jassal, B. |
| 2016-04-05 | Reviewed | D'Eustachio, P. |

### Table of Contents


Introduction 1

Analysis properties 2

Axon guidance 3

- Semaphorin interactions 5
- NCAM signaling for neurite out-growth 6
- Netrin-1 signaling 8
- Signaling by ROBO receptors 9
- L1CAM interactions 11
- EPH-Ephrin signaling 13
- RET signaling 15
- Reelin signalling pathway 16

Table of Contents 17
