## Supplementary Data 1 for "Human SHED-derived extracellular cues activate a specialized neuroprotective and regenerative program in developing retinal ganglion cells": Extracellular Matrix Organization.pdf

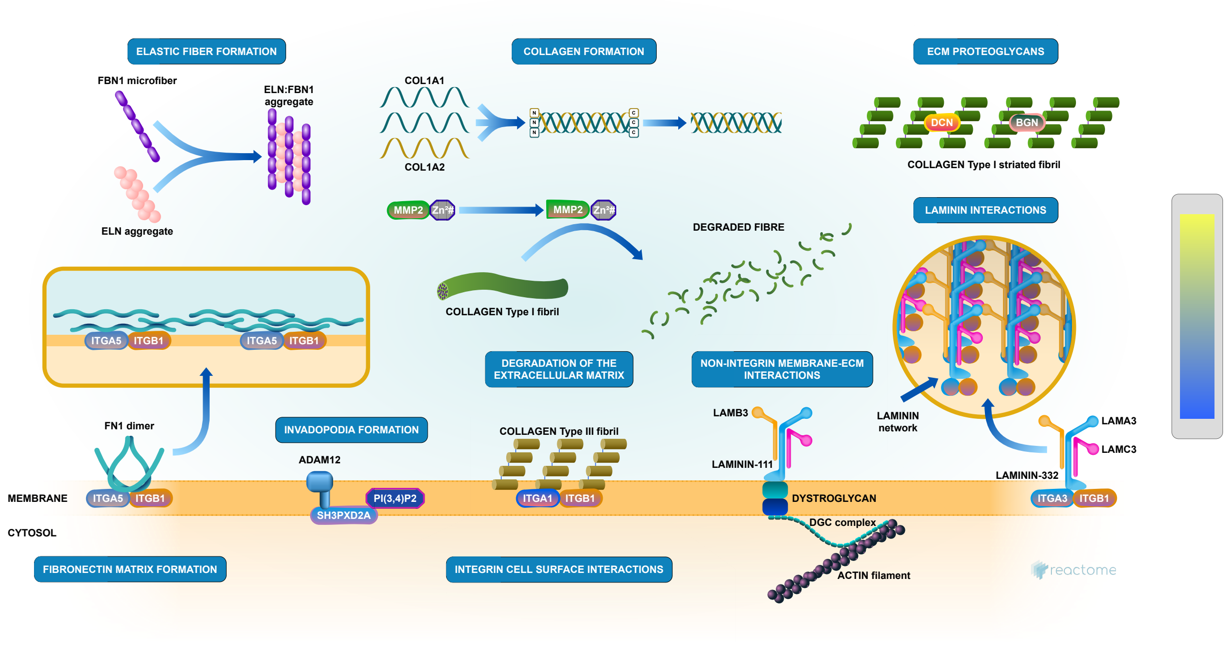


Canty-Laird, EG., D'Eustachio, P., Garapati, P V., Geiger, B., Horwitz, AR., Humphries, MJ., Hynes, R., Jupe, S., May, B., Moreau, V., Muiznieks, LD., Parkinson, J., Reinhardt, DP., Ricard-Blum, S., Venkatesan, N., Yamada, KM.

European Bioinformatics Institute, New York University Langone Medical Center, Ontario Institute for Cancer Research, Oregon Health and Science University.

The contents of this document may be freely copied and distributed in any media, provided the authors, plus the institutions, are credited, as stated under the terms of Creative Commons Attribution 4.0 International (CC BY 4.0) License. For more information see our license.

This is just an excerpt of a full-length report for this pathway. To access the complete report, please download it at the Reactome Textbook.

04/06/2026

### Introduction

Reactome is open-source, open access, manually curated and peer-reviewed pathway database. Pathway annotations are authored by expert biologists, in collaboration with Reactome editorial staff and cross-referenced to many bioinformatics databases. A system of evidence tracking ensures that all assertions are backed up by the primary literature. Reactome is used by clinicians, geneticists, genomics researchers, and molecular biologists to interpret the results of high-throughput experimental studies, by bioinformaticians seeking to develop novel algorithms for mining knowledge from genomic studies, and by systems biologists building predictive models of normal and disease variant pathways.

The development of Reactome is supported by grants from the US National Institutes of Health (P41 HG003751), University of Toronto (CFREF Medicine by Design), European Union (EU STRP, EMI-CD), and the European Molecular Biology Laboratory (EBI Industry program).

#### Literature references

Fabregat, A., Fabregat, A., Fabregat, A., Fabregat, A., Fabregat, A., Fabregat, A. et al. (2017). Reactome pathway analysis: a high-performance in-memory approach. *BMC bioinformatics, 18*, 142. ↗

Sidiropoulos, K., Sidiropoulos, K., Sidiropoulos, K., Sidiropoulos, K., Sidiropoulos, K., Sidiropoulos, K. et al. (2017). Reactome enhanced pathway visualization. *Bioinformatics, 33*, 3461-3467. ↗

Fabregat, A., Fabregat, A., Fabregat, A., Fabregat, A., Fabregat, A., Fabregat, A. et al. (2018). The Reactome Pathway Knowledgebase. *Nucleic Acids Res, 46*, D649-D655. ↗

Fabregat, A., Fabregat, A., Fabregat, A., Fabregat, A., Fabregat, A., Fabregat, A. et al. (2018). Reactome graph database: Efficient access to complex pathway data. *PLoS computational biology, 14*, e1005968. ↗

Reactome database release: 96

This document contains 10 pathways (see Table of Contents)

### Analysis properties

- This is an **expression** analysis: The numbers are used to produce a scaled coloured overlay over Reactome pathway diagrams, as a means to visualize relative expression levels. Note that the numeric values do not have to be expression data, for instance by using gene association scores the same analysis can be used to visualize genotyping results.
  See more
- 175 out of 192 identifiers in the sample were found in Reactome, where 1054 pathways were hit by at least one of them.
- All non-human identifiers have been converted to their human equivalent. ↗
- This report is filtered to show only results for species 'Homo sapiens' and resource 'all resources'.
- The unique ID for this analysis (token) is MjAyNjA2MDMxNDU1NDNfOTEzOA%3D%3D. This ID is valid for at least 7 days in Reactome's server. Use it to access Reactome services with your data.

#### Extracellular matrix organization ↗

**Stable identifier:** R-HSA-1474244

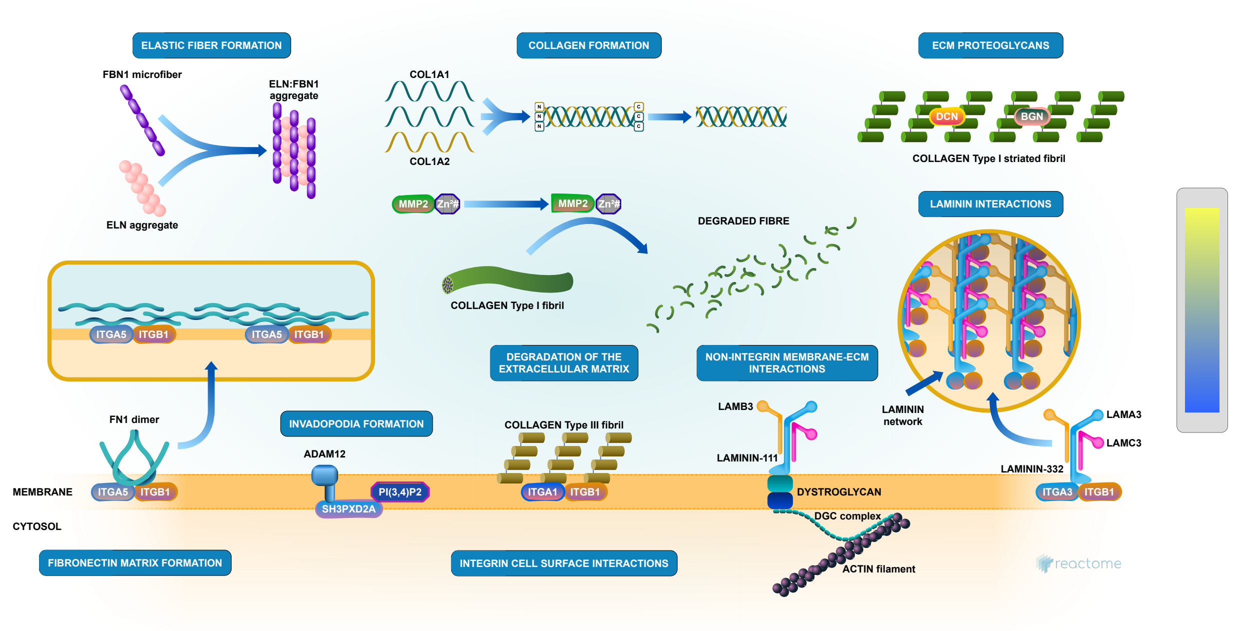


The extracellular matrix is a component of all mammalian tissues, a network consisting largely of the fibrous proteins collagen, elastin and associated-microfibrils, fibronectin and laminins embedded in a viscoelastic gel of anionic proteoglycan polymers. It performs many functions in addition to its structural role; as a major component of the cellular microenvironment it influences cell behaviours such as proliferation, adhesion and migration, and regulates cell differentiation and death (Hynes 2009).

ECM composition is highly heterogeneous and dynamic, being constantly remodeled (Frantz et al. 2010) and modulated, largely by matrix metalloproteinases (MMPs) and growth factors that bind to the ECM influencing the synthesis, crosslinking and degradation of ECM components (Hynes 2009). ECM remodeling is involved in the regulation of cell differentiation processes such as the establishment and maintenance of stem cell niches, branching morphogenesis, angiogenesis, bone remodeling, and wound repair. Redundant mechanisms modulate the expression and function of ECM modifying enzymes. Abnormal ECM dynamics can lead to deregulated cell proliferation and invasion, failure of cell death, and loss of cell differentiation, resulting in congenital defects and pathological processes including tissue fibrosis and cancer.

Collagen is the most abundant fibrous protein within the ECM constituting up to 30% of total protein in multicellular animals. Collagen provides tensile strength. It associates with elastic fibres, composed of elastin and fibrillin microfibrils, which give tissues the ability to recover after stretching. Other ECM proteins such as fibronectin, laminins, and matricellular proteins participate as connectors or linking proteins (Daley et al. 2008).

Chondroitin sulfate, dermatan sulfate and keratan sulfate proteoglycans are structural components associated with collagen fibrils (Scott & Haigh 1985; Scott & Orford 1981), serving to tether the fibril to the surrounding matrix. Decorin belongs to the small leucine-rich repeat proteoglycan family (SLRPs) which also includes biglycan, fibromodulin, lumican and asporin. All appear to be involved in collagen fibril formation and matrix assembly (Ameye & Young 2002).

ECM proteins such as osteonectin (SPARC), osteopontin and thrombospondins -1 and -2, collectively referred to as matricellular proteins (reviewed in Mosher & Adams 2012) appear to modulate cell-matrix interactions. In general they induce de-adhesion, characterized by disruption of focal adhesions and a reorganization of actin stress fibers (Bornstein 2009). Thrombospondin (TS)-1 and -2 bind MMP2. The resulting complex is endocytosed by the low-density lipoprotein receptor-related protein (LRP), clearing MMP2 from the ECM (Yang et al. 2001).

Osteopontin (SPP1, bone sialoprotein-1) interacts with collagen and fibronectin (Mukherjee et al. 1995). It also contains several cell adhesive domains that interact with integrins and CD44.

Aggrecan is the predominant ECM proteoglycan in cartilage (Hardingham & Fosang 1992). Its relatives include versican, neurocan and brevican (Iozzo 1998). In articular cartilage the major non-fibrous macromolecules are aggrecan, hyaluronan and hyaluronan and proteoglycan link protein 1 (HAPLN1). The high negative charge density of these molecules leads to the binding of large amounts of water (Bruckner 2006). Hyaluronan is bound by several large proteoglycans proteoglycans belonging to the hyalectan family that form high-molecular weight aggregates (Roughley 2006), accounting for the turgid nature of cartilage.

The most significant enzymes in ECM remodeling are the Matrix Metalloproteinase (MMP) and A disintegrin and metalloproteinase with thrombospondin motifs (ADAMTS) families (Cawston & Young 2010). Other notable ECM degrading enzymes include plasmin and cathepsin G. Many ECM proteinases are initially present as precursors, activated by proteolytic processing. MMP precursors include an amino prodomain which masks the catalytic Zn-binding motif (Page-McCawet al. 2007). This can be removed by other proteinases, often other MMPs. ECM proteinases can be inactivated by degradation, or blocked by inhibitors. Some of these inhibitors, including alpha2-macroglobulin, alpha1-proteinase inhibitor, and alpha1-chymotrypsin can inhibit a large variety of proteinases (Woessner & Nagase 2000). The tissue inhibitors of metalloproteinases (TIMPs) are potent MMP inhibitors (Brew & Nagase 2010).

#### Literature references


Bosman, FT., Stamenkovic, I. (2003). Functional structure and composition of the extracellular matrix. *J Pathol, 200,* 423-8. ↗

Lu, P., Takai, K., Weaver, VM., Werb, Z. (2011). Extracellular matrix degradation and remodeling in development and disease. *Cold Spring Harb Perspect Biol, 3.* ↗

Frantz, C., Stewart, KM., Weaver, VM. (2010). The extracellular matrix at a glance. *J Cell Sci, 123,* 4195-200. ↗


#### Editions

| | | |
|---|---|---|
| 2011-09-09 | Authored | Jupe, S. |
| 2012-02-21 | Edited | Jupe, S. |
| 2012-02-28 | Reviewed | D'Eustachio, P. |
| 2013-05-21 | Reviewed | Venkatesan, N. |
| 2013-05-22 | Reviewed | Ricard-Blum, S. |
| 2025-06-02 | Reviewed | May, B. |

#### 50 submitted entities found in this pathway, mapping to 55 Reactome entities

| Input | UniProt Id | col1 |
|---|---|---|
| CTSB | P07858 | 12 |
| TIMP2 | P16035 | 14 |
| MMP3 | P08254 | 9 |
| COL6A3 | A8TX70, P12111 | 11 |
| TIMP1 | P01033 | 25 |
| COL6A1 | P12109 | 17 |
| COL6A2 | P12110 | 2 |
| MMP2 | P08253 | 26 |
| LAMB1 | P07942 | 2 |
| MMP1 | P03956 | 15 |
| A2M | P01023 | 5 |
| ACTA2 | P62736, P63267 | 11 |
| PXDN | Q92626 | 2 |
| COL3A1 | P02461 | 12 |
| SERPINH1 | P50454 | 9 |
| FN1 | P02751 | 31 |
| COL4A2 | P08572 | 4 |
| COL4A1 | P02462, Q14055 | 3 |
| HTRA1 | Q92743 | 4 |
| DCN | P07585 | 4 |

| Input | UniProt Id | col1 |
|---|---|---|
| FBLN1 | P23142 | 12 |
| FBN1 | P35555 | 5 |
| FBN2 | P35556, Q75N90 | 2 |
| SPARC | P09486 | 42 |
| EFEMP1 | Q12805 | 4 |
| AGRN | O00468 | 3 |
| FMOD | Q06828 | 5 |
| HSPG2 | P98160 | 2 |
| NID1 | P14543 | 3 |
| COL5A2 | P05997 | 1 |
| ACTN1 | P12814 | 21 |
| COL5A1 | P20908 | 5 |
| LTBP2 | Q14767 | 4 |
| COL1A1 | P02452 | 39 |
| LOXL2 | Q9Y4K0 | 4 |
| COL1A2 | P08123 | 40 |
| ACTB | P60709, P63261 | 54 |
| LTBP1 | Q14766 | 1 |
| COMP | P49747 | 4 |
| CD44 | P16070 | 3 |
| SERPINE1 | P05121 | 24 |
| VCAN | P13611 | 1 |
| TNC | P24821 | 5 |
| PLOD1 | Q02809 | 4 |
| PCOLCE | Q15113 | 15 |
| PLG | P00747 | 2 |
| THBS1 | P07996 | 33 |
| LUM | P51884 | 22 |
| COL12A1 | Q99715 | 7 |
| TKT | Q16832 | 10 |

#### Collagen formation ↗

**Location:** Extracellular matrix organization

**Stable identifier:** R-HSA-1474290

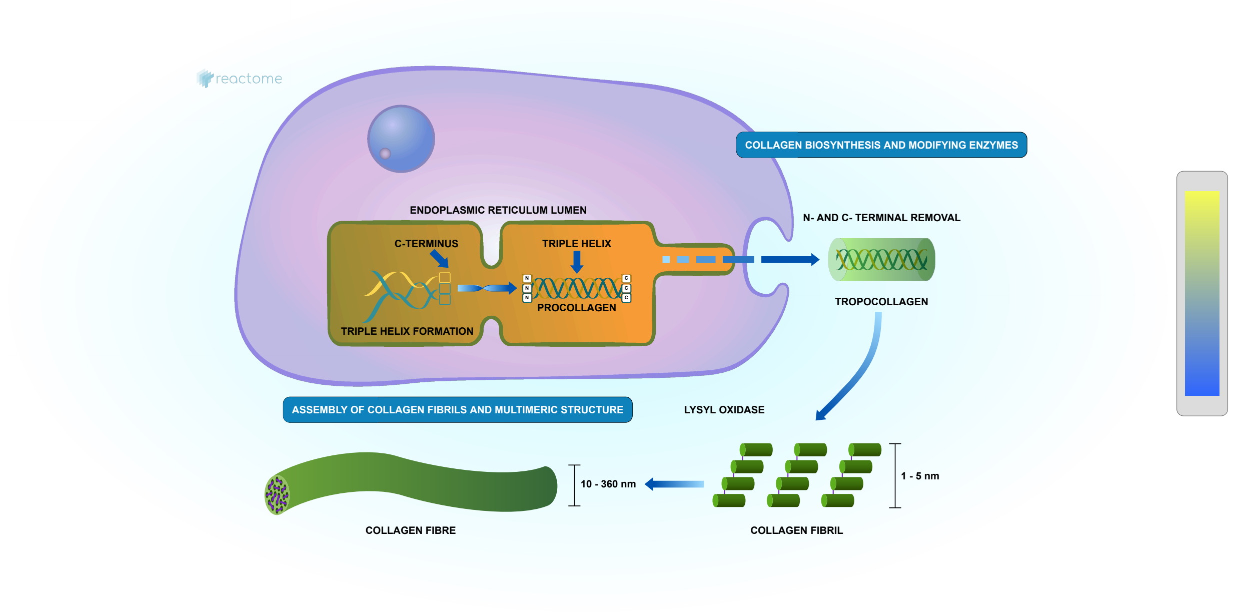


Collagen is a family of at least 29 structural proteins derived from over 40 human genes (Myllyharju & Kivirikko 2004). It is the main component of connective tissue, and the most abundant protein in mammals making up about 25% to 35% of whole-body protein content. A defining feature of collagens is the formation of trimeric left-handed polyproline II-type helical collagenous regions. The packing within these regions is made possible by the presence of the smallest amino acid, glycine, at every third residue, resulting in a repeating motif Gly-X-Y where X is often proline (Pro) and Y often 4-hydroxyproline (4Hyp). Gly-Pro-Hyp is the most common triplet in collagen (Ramshaw et al. 1998). Collagen peptide chains also have non-collagenous domains, with collagen subclasses having common chain structures. Collagen fibrils are mostly found in fibrous tissues such as tendon, ligament and skin. Other forms of collagen are abundant in cornea, cartilage, bone, blood vessels, the gut, and intervertebral disc. In muscle tissue, collagen is a major component of the endomysium, constituting up to 6% of muscle mass. Gelatin, used in food and industry, is collagen that has been irreversibly hydrolyzed. On the basis of their fibre architecture in tissues, the genetically distinct collagens have been divided into subgroups. Group 1 collagens have uninterrupted triple-helical domains of about 300 nm, forming large extracellular fibrils. They are referred to as the fibril-forming collagens, consisting of collagens types I, II, III, V, XI, XXIV and XXVII. Group 2 collagens are types IV and VII, which have extended triple helices (>350 nm) with imperfections in the Gly-X-Y repeat sequences. Group 3 are the short-chain collagens. These have two subgroups. Group 3A have continuous triple-helical domains (type VI, VIII and X). Group 3B have interrupted triple-helical domains, referred to as the fibril-associated collagens with interrupted triple helices (FACIT collagens, Shaw & Olsen 1991). FACITs include collagen IX, XII, XIV, XVI, XIX, XX, XXI, XXII and XXVI plus the transmembrane collagens (XIII, XVII, XXIII and XXV) and the multiple triple helix domains and interruptions (Multiplexin) collagens XV and XVIII (Myllyharju & Kivirikko 2004). The non-collagenous domains of collagens have regulatory functions; several are biologically active when cleaved from the main peptide chain. Fibrillar collagen peptides all have a large triple helical domain (COL1) bordered by N and C terminal extensions, called the N- and C-propeptides, which are cleaved prior to formation of the collagen fibril. The intact form is referred to as a collagen propeptide, not procollagen, which is used to refer to the trimeric triple-helical precursor of collagen before the propeptides are removed. The C-propeptide, also called the NC1 domain, directs chain association during assembly of the procollagen molecule from its three constituent alpha chains (Hulmes 2002).

Fibril forming collagens are the most familiar and best studied subgroup. Collagen fibres are aggregates or bundles of collagen fibrils, which are themselves polymers of tropocollagen complexes, each consisting of three polypeptide chains known as alpha chains. Tropocollagens are considered the subunit of larger collagen structures. They are approximately 300 nm long and 1.5 nm in diameter, with a left-handed triple-helical structure, which becomes twisted into a right-handed coiled-coil 'super helix' in the collagen fibril. Tropocollagens in the extracellular space polymerize spontaneously with regularly staggered ends (Hulmes 2002). In fibrillar collagens the molecules are staggered by about 67 nm, a unit known as D that changes depending upon the hydration state. Each D-period contains slightly more than four collagen molecules so that every D-period repeat of the microfibril has a region containing five molecules in cross-section, called the 'overlap', and a region containing only four molecules, called the 'gap'. The triple-helices are arranged in a hexagonal or quasi-hexagonal array in cross-section, in both the gap and overlap regions (Orgel et al. 2006). Collagen molecules cross-link covalently to each other via lysine and hydroxylysine side chains. These cross-links are unusual, occuring only in collagen and elastin, a related protein.

The macromolecular structures of collagen are diverse. Several group 3 collagens associate with larger collagen fibers, serving as molecular bridges which stabilize the organization of the extracellular matrix. Type IV collagen is arranged in an interlacing network within the dermal-epidermal junction and vascular basement membranes. Type VI collagen forms distinct microfibrils called beaded filaments. Type VII collagen forms anchoring fibrils. Type VIII and X collagens form hexagonal networks. Type XVII collagen is a component of hemidesmosomes where it is complexed wtih alpha6Beta4 integrin, plectin, and laminin-332 (de Pereda et al. 2009). Type XXIX collagen has been recently reported to be a putative epidermal collagen with highest expression in suprabasal layers (Soderhall et al. 2007). Collagen fibrils/aggregates arranged in varying combinations and concentrations in different tissues provide specific tissue properties. In bone, collagen triple helices lie in a parallel, staggered array with 40 nm gaps between the ends of the tropocollagen subunits, which probably serve as nucleation sites for the deposition of crystals of the mineral component, hydroxyapatite (Ca10(PO4)6(OH)2) with some phosphate. Collagen structure affects cell-cell and cell-matrix communication, tissue construction in growth and repair, and is changed in development and disease (Sweeney et al. 2006, Twardowski et al. 2007). A single collagen fibril can be heterogeneous along its axis, with significantly different mechanical properties in the gap and overlap regions, correlating with the different molecular organizations in these regions (Minary-Jolandan & Yu 2009).

#### Literature references


Prockop, DJ., Kivirikko, KI. (1995). Collagens: molecular biology, diseases, and potentials for therapy. *Annu Rev Biochem, 64*, 403-34. ↗

Gordon, MK., Hahn, RA. (2010). Collagens. *Cell Tissue Res, 339*, 247-57. ↗


#### Editions

| | | |
|---|---|---|
| 2011-08-05 | Authored | Jupe, S. |
| 2012-04-11 | Edited | Jupe, S. |
| 2012-05-24 | Reviewed | Canty-Laird, EG. |

#### 18 submitted entities found in this pathway, mapping to 20 Reactome entities

| Input | UniProt Id | col1 |
|---|---|---|
| CTSB | P07858 | 12 |
| MMP3 | P08254 | 9 |
| COL6A3 | A8TX70, P12111 | 11 |
| PCOLCE | Q15113 | 15 |
| COL6A1 | P12109 | 17 |
| COL6A2 | P12110 | 2 |
| COL4A1 | P02462, Q14055 | 3 |
| COL4A2 | P08572 | 4 |
| COL1A1 | P02452 | 39 |
| COL1A2 | P08123 | 40 |
| PXDN | Q92626 | 2 |
| COL5A1 | P20908 | 5 |
| COL5A2 | P05997 | 1 |
| COL3A1 | P02461 | 12 |
| SERPINH1 | P50454 | 9 |
| LOXL2 | Q9Y4K0 | 4 |
| PLOD1 | Q02809 | 4 |
| COL12A1 | Q99715 | 7 |

#### Fibronectin matrix formation ↗

**Location:** Extracellular matrix organization

**Stable identifier:** R-HSA-1566977

**Compartments:** extracellular region

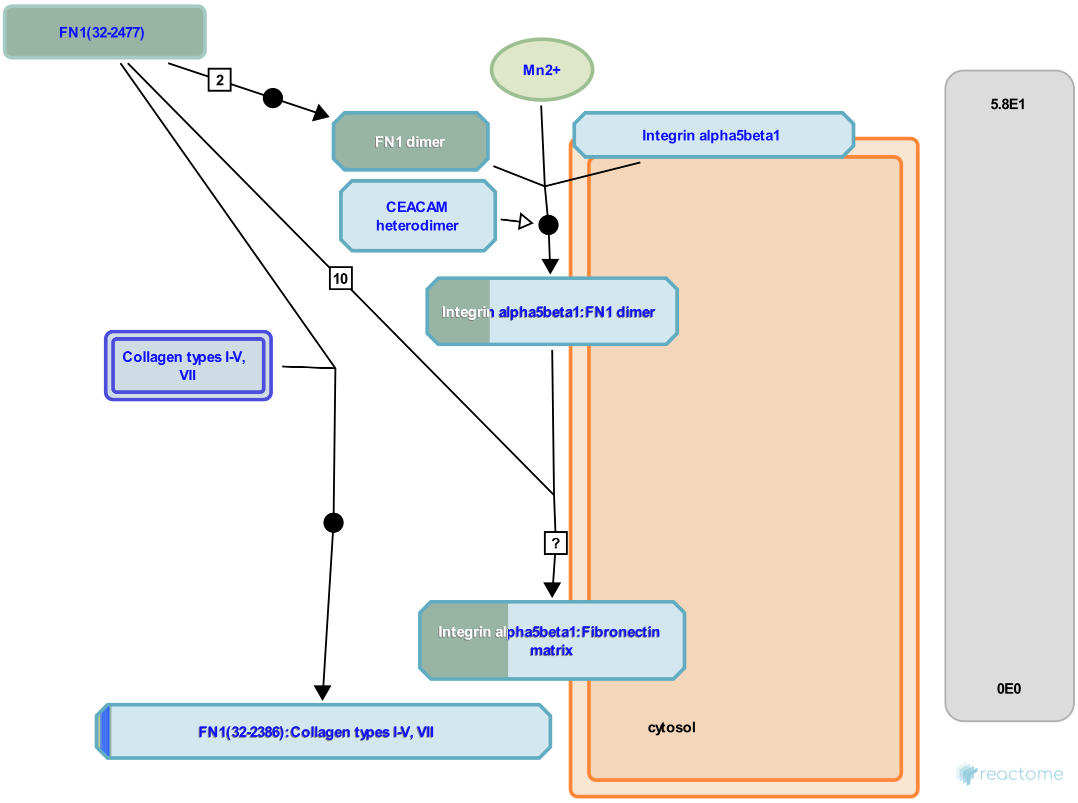


Fibronectin (FN1) is found in the extracellular matrix (ECM) of all cells as linear and branched networks that surround and connect neighbouring cells (Singh et al. 2010). Prior to matrix formation FN1 exists as a protein dimer. Often the two peptide chains represent differentially-spliced variants. The chains are linked by a pair of C-terminal disulfide bonds which are essential for subsequent multimerization (Schwarzbaur 1991). FN1 monomers have a molecular weight of 230-270 kDa depending on alternative splicing and contain three types of repeating unit, I, II, and III. I and II are stabilized by intra-chain disulfide bonds. The absence of disulfide bonds in type III modules allows them to partially unfold under applied force (Erickson 2002). Three regions of variable splicing occur along the length of the FN1 monomer (Mao & Schwarzbauer 2005). One or both of the 'extra' type III modules EIIIA and EIIIB may be present in cellular FN1, but never in plasma FN1. A variable (V) region exists between the 14th and 15th type III module. This contains the binding site for alpha4 beta1 and alpha4beta7 integrins. It is present in most cellular FN1, occasionally in plasma FN1. The modules are arranged into several functional and protein-binding domains. There are four FN1-binding domains (Mao & Schwarzbauer 2005). One of these domains (I1-5), referred to as the 'assembly domain', is required for the initiation of FN1 matrix assembly. Modules III9-10 correspond to the 'cell-binding domain' of FN1. The Arg-Gly-Asp (RGD) integrin binding sequence located in III10 is the primary site of FN1 to cell attachment, mediated predominantly by alpha5 beta1 and alphaV beta3 integrins. The 'synergy site' in III9 modulates FN1's association with alpha5 beta1 integrins. FN1 also contains interaction domains for fibrin (I1-5, I10-12), collagen (I6-9, II1-2), fibulin-1 (III13-14), heparin, syndecan (III12-14) and fibrillin-1 (I6-9) (Mao & Schwartzbauer 2005, Sabatier et al. 2009).

FN1 dimer binding to alpha5beta1 integrin stimulates self-association. Binding is thought to lead to a conformational change in FN1 that triggers the addition of further FN1 dimers (Singh et al. 2010). I1-5 functions as a unit that is the primary FN1 matrix assembly domain (Sottile et al. 1991) but other units are likely to be involved (Singh et al. 2010), the process is not fully understood.

Several ECM proteins appear to require the FN1 matrix for their own assembly. Fibrillin-1 containing microfibrils are formed when fibrillin-1 multimers bind to the FN1 matrix (Sabatier et al. 2009). FN1 polymerization promotes the deposition of type I and type III collagen (Sottile and Hocking 2002, Velling et al. 2002). Inhibition of FN1 polymerization increases its turnover and a concomitant loss of collagen types I and III from the ECM (Sottile and

Hocking 2002, Sottile et al. 2007). FN1 is regulated by matrix metalloproteinases, particularly MMP14 (Shi & Sottile 2011).

#### Literature references

Singh, P., Carraher, C., Schwarzbauer, JE. (2010). Assembly of fibronectin extracellular matrix. *Annu. Rev. Cell Dev. Biol., 26*, 397-419. ↗

#### Editions

| | | |
|---|---|---|
| 2011-07-12 | Authored, Edited | Jupe, S. |
| 2013-02-08 | Reviewed | Reinhardt, DP. |
| 2025-06-02 | Reviewed | May, B. |

#### 8 submitted entities found in this pathway, mapping to 8 Reactome entities

| Input | UniProt Id | col1 |
|---|---|---|
| COL5A2 | P05997 | 1 |
| COL4A1 | P02462 | 3 |
| COL4A2 | P08572 | 4 |
| COL5A1 | P20908 | 5 |
| FN1 | P02751 | 31 |
| COL1A1 | P02452 | 39 |
| COL1A2 | P08123 | 40 |
| COL3A1 | P02461 | 12 |

#### Elastic fibre formation ↗

**Location:** Extracellular matrix organization

**Stable identifier:** R-HSA-1566948

**Compartments:** extracellular region

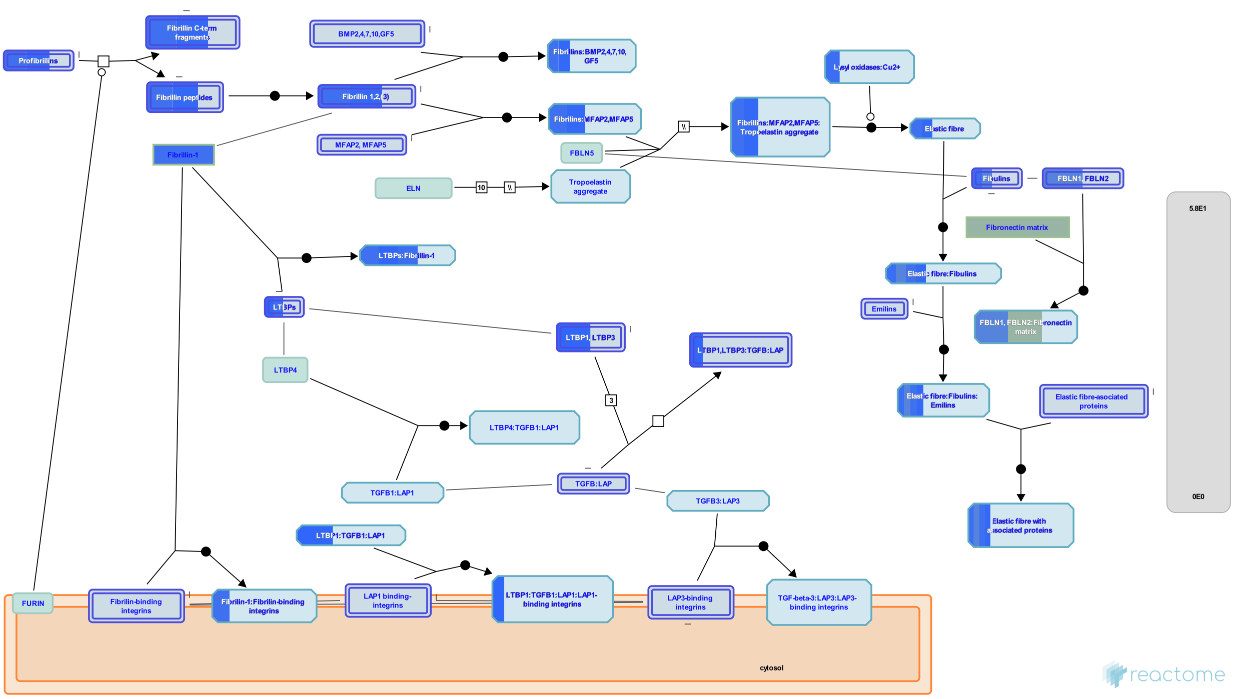


Elastic fibres (EF) are a major structural constituent of dynamic connective tissues such as large arteries and lung parenchyma, where they provide essential properties of elastic recoil and resilience. EF are composed of a central cross-linked core of elastin, surrounded by a mesh of microfibrils, which are composed largely of fibrillin. In addition to elastin and fibrillin-1, over 30 ancillary proteins are involved in mediating important roles in elastic fibre assembly as well as interactions with the surrounding environment. These include fibulins, elastin microfibril interface located proteins (EMILINs), microfibril-associated glycoproteins (MAGPs) and Latent TGF-beta binding proteins (LTBPs). Fibulin-5 for example, is expressed by vascular smooth muscle cells and plays an essential role in the formation of elastic fibres through mediating interactions between elastin and fibrillin (Yanigasawa et al. 2002, Freeman et al. 2005). In addition, it plays a role in cell adhesion through integrin receptors and has been shown to influence smooth muscle cell proliferation (Yanigasawa et al. 2002, Nakamura et al. 2002). EMILINs are a family of homologous glycoproteins originally identified in extracts of aortas. Found at the elastin-fibrillin interface, early studies showed that antibodies to EMILIN can affect the process of elastic fibre formation (Bressan et al. 1993). EMILIN1 has been shown to bind elastin and fibulin-5 and appears to coordinate their common interaction (Zanetti et al. 2004). MAGPs are found to co-localize with microfibrils. MAGP-1, for example, binds strongly to an N-terminal sequence of fibrillin-1. Other proteins found associated with microfibrils include vitronectin (Dahlback et al. 1990).

Fibrillin is most familiar as a component of elastic fibres but microfibrils with no elastin are found in the ciliary zonules of the eye and invertebrate circulatory systems. The addition of elastin to microfibrils is a vertebrate adaptation to high pulsatile pressures in their closed circulatory systems (Faury et al. 2003). Elastin appears to have emerged after the divergence of jawless vertebrates from other vertebrates (Sage 1982).

Fibrillin-1 is the major structural component of microfibrils. Fibrillin-2 is expressed earlier in development than fibrillin-1 and may be important for elastic fiber formation (Zhang et al. 1994). Fibrillin-3 arose as a duplication of fibrillin-2 that did not occur in the rodent lineage. It was first isolated from human brain (Corson et al. 2004).

Fibrillin assembly is not as well defined as elastin assembly. The primary structure of fibrillin is dominated by calcium binding epidermal growth factor like repeats (Kielty et al. 2002). Fibrillin may form dimers or trimers before secretion. However, multimerisation predominantly occurs outside the cell. Formation of fibrils appears to require cell surface structures suggesting an involvement of cell surface receptors. Fibrillin is assembled pericellularly (i.e. on or close to the cell surface) into microfibrillar arrays that undergo time dependent maturation into microfibrils with beaded-string appearance. Transglutaminase forms gamma glutamyl epsilon lysine isopeptide bonds within or between peptide chains. Additionally, intermolecular disulfide bond formation between fibrillins is an important contributor to fibril maturation (Reinhardt et al. 2000).

Models of fibrillin-1 microfibril structure suggest that the N-terminal half of fibrillin-1 is asymmetrically exposed in outer filaments, while the C-terminal half is buried in the interior (Kuo et al. 2007). Fibrillinopathies include Marfan syndrome, familial ectopia lentis, familial thoracic aneurysm, all due to mutations in the fibrillin-1 gene FBN1, and congenital contractural arachnodactyly which is caused by mutation of FBN2 (Maslen & Glanville 1993, Davis & Summers 2012).

In vivo assembly of fibrillin requires the presence of extracellular fibronectin fibres (Sabatier et al. 2009). Fibrillins have Arg-Gly-Asp (RGD) sequences that interact with integrins (Pfaff et al. 1996, Sakamoto et al. 1996, Bax et al., 2003, Jovanovic et al. 2008) and heparin-binding domains that interact with a cell-surface heparan sulfate proteoglycan (Tiedemann et al. 2001) possibly a syndecan (Ritty et al. 2003). Fibrillins also have a major role in binding and sequestering growth factors such as TGF beta into the ECM (Neptune et al. 2003). Proteoglycans such as versican (Isogai et al. 2002), biglycan, and decorin (Reinboth et al. 2002) can interact with the microfibrils. They confer specific properties including hydration, impact absorption, molecular sieving, regulation of cellular activities, mediation of growth factor association, and release and transport within the extracellular matrix (Buczek-Thomas et al. 2002). In addition, glycosaminoglycans have been shown to interact with tropoelastin through its lysine side chains (Wu et al. 1999), regulating tropoelastin assembly (Tu & Weiss 2008).

Elastin is synthesized as a 70kDa monomer called tropoelastin, a highly hydrophobic protein composed largely of two types of domains that alternate along the polypeptide chain. Hydrophobic domains are rich in glycine, proline, alanine, leucine and valine. These amino acids occur in characteristic short (3-9 amino acids) tandem repeats, with a flexible and highly dynamic structure (Floquet et al. 2004). Unlike collagen, glycine in elastin is not rigorously positioned every 3 residues. However, glycine is distributed frequently throughout all hydrophobic domains of elastin, and displays a strong preference for inter-glycine spacing of 0-3 residues (Rauscher et al. 2006).

Elastic fibre formation involves the deposition of tropoelastin onto a template of fibrillin rich microfibrils. Recent results suggest that the first step of elastic fiber formation is the organization of small globules of elastin on the cell surface followed by globule aggregation into microfibres (Kozel et al. 2006). An important contribution to the initial stages assembly is thought to be made by the intrinsic ability of the protein to direct its own polymeric organization in a process termed 'coacervation' (Bressan et al. 1986). This self-assembly process appears to be determined by interactions between hydrophobic domains (Bressan et al. 1986, Vrhovski et al. 1997, Bellingham et al. 2003, Cirulis & Keeley 2010) which result in alignment of the cross-linking domains, allowing the stabilization of elastin through the formation of cross-links generated through the oxidative deamination of lysine residues, catalyzed by members of the lysyl oxidase (LOX) family (Reiser et al. 1992, Mithieux & Weiss 2005). The first step in the cross-linking reaction is the oxidative formation of the delta aldehyde, known as alpha aminoadipic semialdehyde or allysine (Partridge 1963). Subsequent reactions that are probably spontaneous lead to the formation of cross-links through dehydrolysinonorleucine and allysine aldol, a trifunctional cross-link dehydromerodesmosine and two tetrafunctional cross-links desmosine and isodesmosine (Lucero & Kagan 2006), which are unique to elastin. These cross-links confer mechanical integrity and high durability. In addition to their role in self-assembly, hydrophobic domains provide elastin with its elastomeric properties, with initial studies suggesting that the elastomeric propereties of elastin are driven through changes in entropic interactions with surrounding water molecules (Hoeve & Flory 1974).

A very specific set of proteases, broadly grouped under the name elastases, is responsible for elastin remodelling (Antonicelli et al. 2007). The matrix metalloproteinases (MMPs) are particularly important in elastin breakdown, with MMP2, 3, 9 and 12 explicitly shown to degrade elastin (Ra & Parks 2007). Nonetheless, elastin typically displays a low turnover rate under normal conditions over a lifetime (Davis 1993).

#### Literature references

Kozel, BA., Rongish, BJ., Czirok, A., Zach, J., Little, CD., Davis, EC. et al. (2006). Elastic fiber formation: a dynamic view of extracellular matrix assembly using timer reporters. *J Cell Physiol, 207*, 87-96. ↗

#### Editions

| | | |
|---|---|---|
| 2012-04-30 | Authored | Jupe, S. |
| 2012-11-02 | Reviewed | Muiznieks, LD. |
| 2012-11-12 | Edited | Jupe, S. |
| 2013-02-27 | Reviewed | Parkinson, J. |

#### 8 submitted entities found in this pathway, mapping to 9 Reactome entities

| Input | UniProt Id | col1 |
| --- | --- | --- |
| FBLN1 | P23142 | 12 |
| FBN1 | P35555 | 5 |
| FBN2 | P35556, Q75N90 | 2 |
| EFEMP1 | Q12805 | 4 |
| FN1 | P02751 | 31 |
| LTBP2 | Q14767 | 4 |
| LOXL2 | Q9Y4K0 | 4 |
| LTBP1 | Q14766 | 1 |

#### Laminin interactions ↗

**Location:** Extracellular matrix organization

**Stable identifier:** R-HSA-3000157

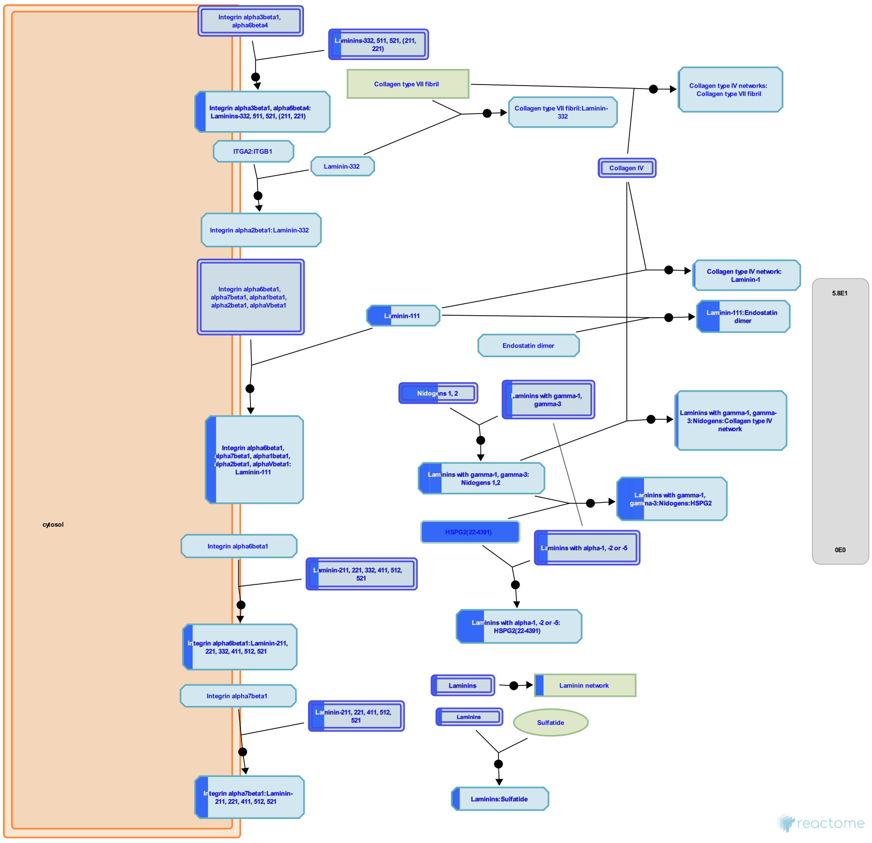

Laminins are a large family of conserved, multidomain trimeric basement membrane proteins. There are many theoretical trimer combinations but only 18 have been described (Domogatskaya et al. 2012, Miner 2008, Macdonald et al. 2010) and the existence of isoforms laminin-212 and/or laminin-222 (Durbeej et al. 2010) awaits further confirmation. The chains assemble through coiled-coil domains at their C-terminal end. Alpha chains additionally have a large C-terminal globular domain containing five LG subdomains (LG1-5). The N termini are often referred to as the short arms. These have varying numbers of laminin-type epidermal growth factor-like (LE) repeats. Trimer assembly is controlled by highly specific coiled-coil interactions (Domogatskaya et al. 2012). Some laminin isoforms are modified extracellularly by proteolytic processing at the N- or C-terminal ends prior to their binding to cellular receptors or other matrix molecules (Tzu & Marinkovitch 2008).

The cell adhesion properties of laminins are mediated primarily through the alpha chain G domain to integrins, dystroglycan, Lutheran glycoprotein, or sulfated glycolipids. The N-terminal globular domains of the alpha-1 (Colognato-Pyke et al. 1995) and alpha-2 chains (Colognato et al. 1997) and globular domains VI (Nielsen & Yamada 2001) and IVa (Sasaki & Timpl 2001) of the alpha-5 chain can bind to several integrin isoforms (alpha1beta1, alpha2beta1, alpha3beta1, and alphaVbeta3), which enables cell binding at both ends of laminins with these alpha chains.

#### Literature references


Domogatskaya, A., Rodin, S., Tryggvason, K. (2012). Functional diversity of laminins. *Annu. Rev. Cell Dev. Biol., 28*, 523-53. ↗


#### Editions

| | | |
|---|---|---|
| 2008-05-07 | Reviewed | Humphries, MJ., Yamada, KM., Hynes, R. |
| 2012-08-08 | Authored | Jupe, S. |
| 2013-08-13 | Edited | Jupe, S. |
| 2013-08-13 | Reviewed | Ricard-Blum, S. |

##### 5 submitted entities found in this pathway, mapping to 5 Reactome entities

| Input | UniProt Id | col1 |
|---|---|---|
| LAMB1 | P07942 | 2 |
| COL4A1 | P02462 | 3 |
| COL4A2 | P08572 | 4 |
| HSPG2 | P98160 | 2 |
| NID1 | P14543 | 3 |

#### Non-integrin membrane-ECM interactions ↗

**Location:** Extracellular matrix organization

**Stable identifier:** R-HSA-3000171

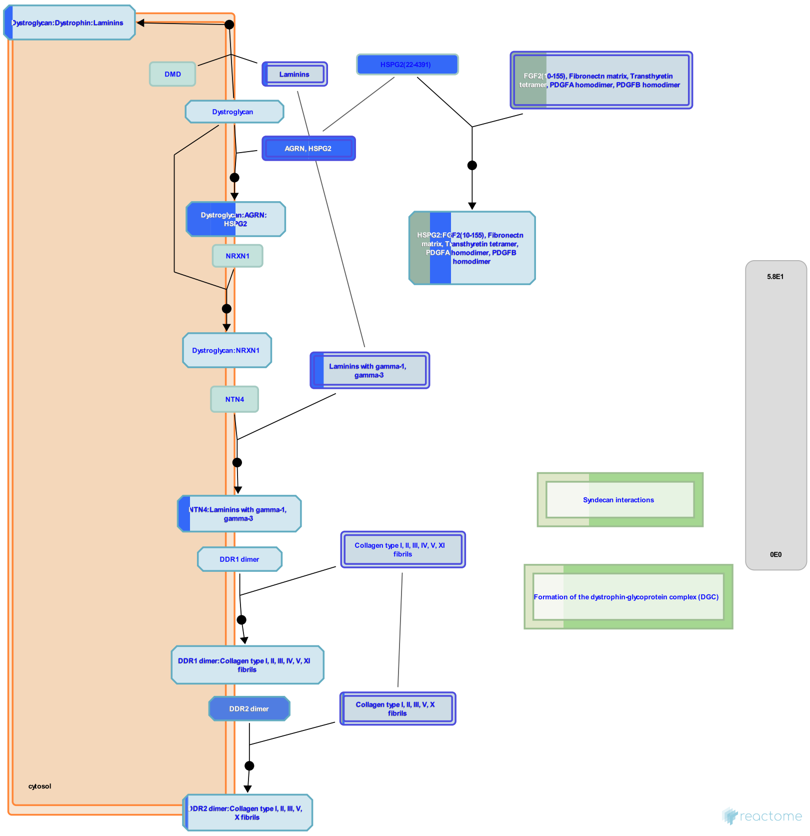


Several non-integrin membrane proteins interact with extracellular matrix proteins. Transmembrane proteoglycans may associate with integrins and growth factor receptors to influence their function, or they can signal independently, often influencing the actin cytoskeleton.

#### Literature references


Rosso, F., Giordano, A., Barbarisi, M., Barbarisi, A. (2004). From cell-ECM interactions to tissue engineering. *J. Cell. Physiol., 199*, 174-80. ↗

Couchman, JR. (2010). Transmembrane signaling proteoglycans. *Annu. Rev. Cell Dev. Biol., 26*, 89-114. ↗


#### Editions

| | | |
|---|---|---|
| 2012-07-31 | Authored | Jupe, S. |
| 2013-04-26 | Edited | Jupe, S. |
| 2013-05-22 | Reviewed | Ricard-Blum, S. |

#### 17 submitted entities found in this pathway, mapping to 19 Reactome entities

| Input | UniProt Id | col1 |
|---|---|---|
| ACTN1 | P12814 | 21 |
| COL4A1 | P02462 | 3 |
| LAMB1 | P07942 | 2 |
| COL4A2 | P08572 | 4 |
| COL1A1 | P02452 | 39 |
| ACTA2 | P62736, P63267 | 11 |
| COL1A2 | P08123 | 40 |

| Input | UniProt Id | col1 |
|---|---|---|
| ACTB | P60709, P63261 | 54 |
| COL5A1 | P20908 | 5 |
| COL5A2 | P05997 | 1 |
| COL3A1 | P02461 | 12 |
| FN1 | P02751 | 31 |
| AGRN | O00468 | 3 |
| TNC | P24821 | 5 |
| HSPG2 | P98160 | 2 |
| TKT | Q16832 | 10 |
| THBS1 | P07996 | 33 |

#### ECM proteoglycans ↗

**Location:** Extracellular matrix organization

**Stable identifier:** R-HSA-3000178

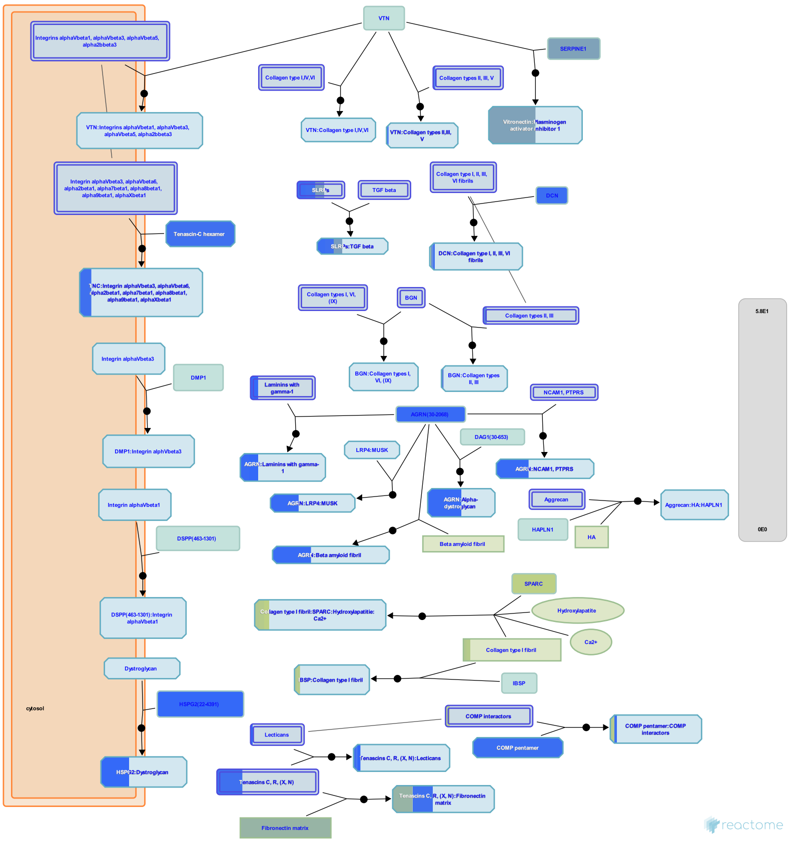

Proteoglycans are major components of the extracellular matrix. In cartilage the matrix constitutes more than 90% of tissue dry weight. Proteoglycans are proteins substituted with glycosaminoglycans (GAGs), linear polysaccharides consisting of a repeating disaccharide, generally of an acetylated amino sugar alternating with a uronic acid. Most proteoglycans are located in the extracellular space. Proteoglycans are highly diverse, both in terms of the core proteins and the subtypes of GAG chains, namely chondroitin sulfate (CS), keratan sulfate (KS), dermatan sulfate (DS) and heparan sulfate (HS). Hyaluronan is a non-sulfated GAG whose molecular weight runs into millions of Dalton; in articular cartilage, a single hyaluronan molecule can hold upto 100 aggrecan molecules and these aggregates are stabilized by a link protein.

#### Literature references


Esko, JD., Esko, JD., Kimata, K., Lindahl, U., Varki, A., Cummings, RD. et al. (2009). Proteoglycans and Sulfated Glycosaminoglycans.

Kim, SH., Turnbull, J., Guimond, S. (2011). Extracellular matrix and cell signalling: the dynamic cooperation of integrin, proteoglycan and growth factor receptor. *J. Endocrinol.*, *209*, 139-51. ↗

Hay, E. (1991). Cell Biology of Extracellular Matrix. *Springer-Verlag*.

Hay, E. (1991). Proteoglycans: structure and function, Cell Biology of Extracellular Matrix.


#### Editions

| | | |
|---|---|---|
| 2013-01-10 | Authored | Jupe, S. |
| 2013-04-26 | Edited | Jupe, S. |
| 2013-05-21 | Reviewed | Venkatesan, N. |
| 2013-05-22 | Reviewed | Ricard-Blum, S. |

#### 22 submitted entities found in this pathway, mapping to 24 Reactome entities

| Input | UniProt Id | col1 |
|---|---|---|
| COL6A3 | A8TX70, P12111 | 11 |
| COL6A1 | P12109 | 17 |
| COL6A2 | P12110 | 2 |
| COL4A1 | P02462, Q14055 | 3 |
| LAMB1 | P07942 | 2 |
| COL4A2 | P08572 | 4 |
| LUM | P51884 | 22 |
| COL1A1 | P02452 | 39 |
| COMP | P49747 | 4 |
| COL1A2 | P08123 | 40 |
| COL5A1 | P20908 | 5 |
| COL5A2 | P05997 | 1 |
| COL3A1 | P02461 | 12 |
| FN1 | P02751 | 31 |
| SPARC | P09486 | 42 |
| DCN | P07585 | 4 |
| AGRN | O00468 | 3 |
| SERPINE1 | P05121 | 24 |
| VCAN | P13611 | 1 |
| FMOD | Q06828 | 5 |
| TNC | P24821 | 5 |
| HSPG2 | P98160 | 2 |

#### Degradation of the extracellular matrix ↗

**Location:** Extracellular matrix organization

**Stable identifier:** R-HSA-1474228

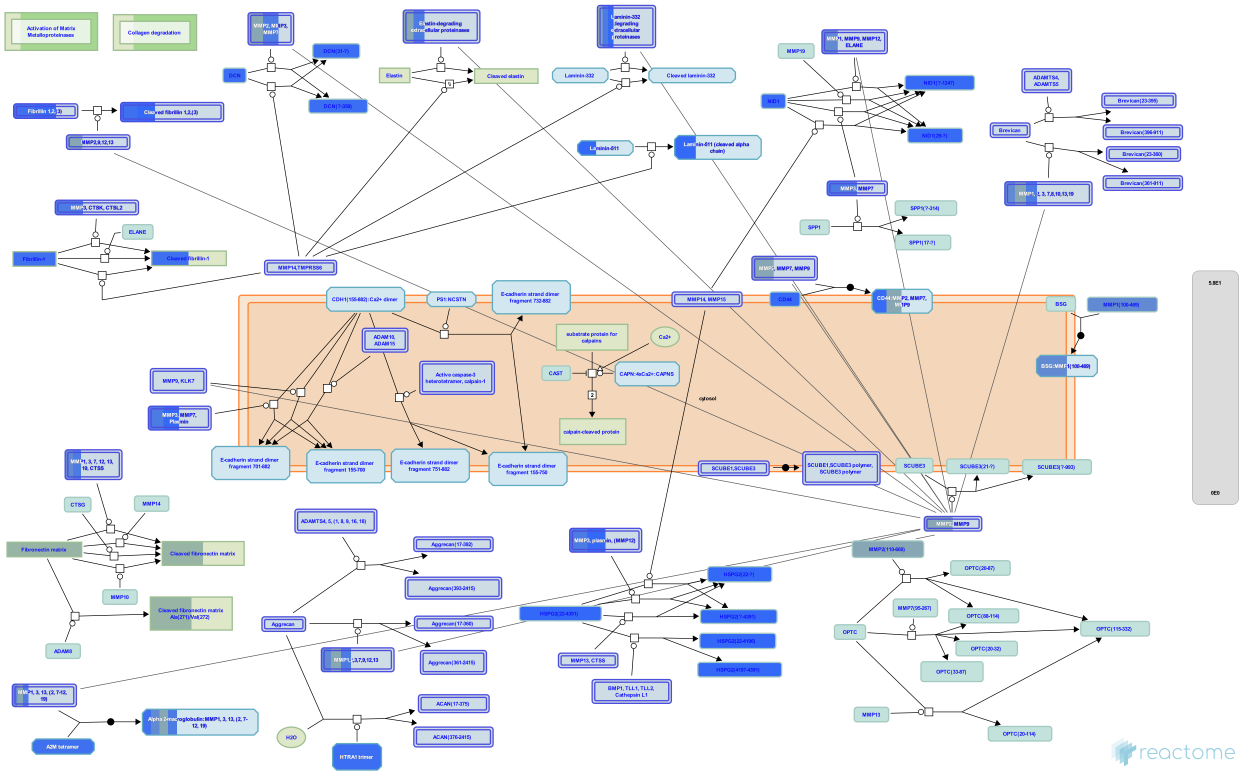

Matrix metalloproteinases (MMPs), previously referred to as matrixins because of their role in degradation of the extracellular matrix (ECM), are zinc and calcium dependent proteases belonging to the metzincin family. They contain a characteristic zinc-binding motif HEXXHXXGXXH (Stocker & Bode 1995) and a conserved Methionine which forms a Met-turn. Humans have 24 MMP genes giving rise to 23 MMP proteins, as MMP23 is encoded by two identical genes. All MMPs contain an N-terminal secretory signal peptide and a prodomain with a conserved PRCGXPD motif that in the inactive enzyme is localized with the catalytic site, the cysteine acting as a fourth unpaired ligand for the catalytic zinc atom. Activation involves delocalization of the domain containing this cysteine by a conformational change or proteolytic cleavage, a mechanism referred to as the cysteine-switch (Van Wart & Birkedal-Hansen 1990). Most MMPs are secreted but the membrane type MT-MMPs are membrane anchored and some MMPs may act on intracellular proteins. Various domains determine substrate specificity, cell localization and activation (Hadler-Olsen et al. 2011). MMPs are regulated by transcription, cellular location (most are not activated until secreted), activating proteinases that can be other MMPs, and by metalloproteinase inhibitors such as the tissue inhibitors of metalloproteinases (TIMPs). MMPs are best known for their role in the degradation and removal of ECM molecules. In addition, cleavage of the ECM and other cell surface molecules can release ECM-bound growth factors, and a number of non-ECM proteins are substrates of MMPs (Nagase et al. 2006). MMPs can be divided into subgroups based on domain structure and substrate specificity but it is clear that these are somewhat artificial, many MMPs belong to more than one functional group (Vise & Nagase 2003, Somerville et al. 2003).

##### Literature references


Lu, P., Takai, K., Weaver, VM., Werb, Z. (2011). Extracellular matrix degradation and remodeling in development and disease. *Cold Spring Harb Perspect Biol, 3.* ↗


##### Editions

| | | |
|---|---|---|
| 2011-09-09 | Authored | Jupe, S. |
| 2012-02-21 | Edited | Jupe, S. |
| 2012-02-28 | Reviewed | D'Eustachio, P. |

#### 28 submitted entities found in this pathway, mapping to 31 Reactome entities

| Input | UniProt Id | col1 |
|---|---|---|
| CTSB | P07858 | 12 |
| TIMP2 | P16035 | 14 |
| MMP3 | P08254 | 9 |
| COL6A3 | A8TX70, P12111 | 11 |
| TIMP1 | P01033 | 25 |
| COL6A1 | P12109 | 17 |
| COL6A2 | P12110 | 2 |
| COL4A1 | P02462, Q14055 | 3 |
| MMP2 | P08253 | 26 |
| LAMB1 | P07942 | 2 |
| COL4A2 | P08572 | 4 |
| MMP1 | P03956 | 15 |
| A2M | P01023 | 5 |
| COL1A1 | P02452 | 39 |
| COL1A2 | P08123 | 40 |
| COL5A1 | P20908 | 5 |
| COL5A2 | P05997 | 1 |
| COL3A1 | P02461 | 12 |
| FN1 | P02751 | 31 |
| HTRA1 | Q92743 | 4 |
| DCN | P07585 | 4 |
| FBN1 | P35555 | 5 |
| FBN2 | P35556, Q75N90 | 2 |
| COL12A1 | Q99715 | 7 |
| PLG | P00747 | 2 |
| HSPG2 | P98160 | 2 |
| NID1 | P14543 | 3 |
| CD44 | P16070 | 3 |

#### Integrin cell surface interactions ↗

**Location:** Extracellular matrix organization

**Stable identifier:** R-HSA-216083

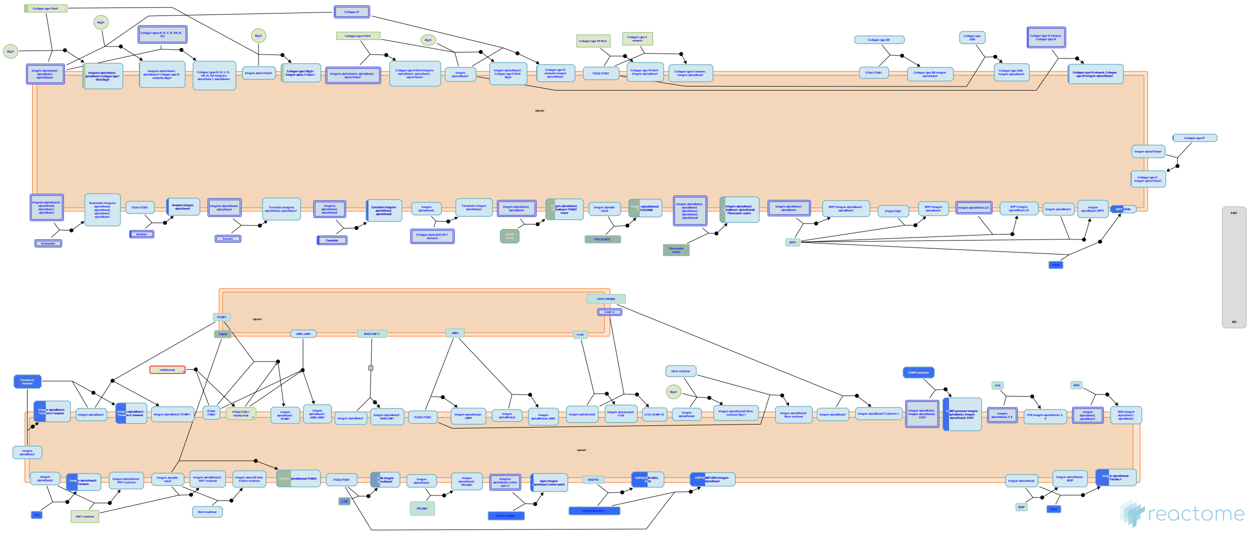

The extracellular matrix (ECM) is a network of macro-molecules that underlies all epithelia and endothelia and that surrounds all connective tissue cells. This matrix provides the mechanical strength and also influences the behavior and differentiation state of cells in contact with it. The ECM are diverse in composition, but they generally comprise a mixture of fibrillar proteins, polysaccharides synthesized, secreted and organized by neighboring cells. Collagens, fibronectin, and laminins are the principal components involved in cell matrix interactions; other components, such as vitronectin, thrombospondin, and osteopontin, although less abundant, are also important adhesive molecules.
Integrins are the receptors that mediate cell adhesion to ECM. Integrins consists of one alpha and one beta subunit forming a noncovalently bound heterodimer. 18 alpha and 8 beta subunits have been identified in humans that combine to form 24 different receptors.
The integrin dimers can be broadly divided into three families consisting of the beta1, beta2/beta7, and beta3/alphaV integrins. beta1 associates with 12 alpha-subunits and can be further divided into RGD-, collagen-, or laminin binding and the related alpha4/alpha9 integrins that recognise both matrix and vascular ligands. beta2/beta7 integrins are restricted to leukocytes and mediate cell-cell rather than cell-matrix interactions, although some recognize fibrinogen. The beta3/alphaV family members are all RGD receptors and comprise aIIbb3, an important receptor on platelets, and the remaining b-subunits, which all associate with alphaV. It is the collagen receptors and leukocyte-specific integrins that contain alpha A-domains.

#### Literature references


Faull, RJ., Ginsberg, MH. (1996). Inside-out signaling through integrins. *J Am Soc Nephrol, 7*, 1091-7. ↗

Arnaout, MA., Goodman, SL., Xiong, JP. (2002). Coming to grips with integrin binding to ligands. *Curr Opin Cell Biol, 14*, 641-51. ↗

White, DJ., Puranen, S., Johnson, MS., Heino, J. (2004). The collagen receptor subfamily of the integrins. *Int J Biochem Cell Biol, 36*, 1405-10. ↗

Boudreau, NJ., Jones, PL. (1999). Extracellular matrix and integrin signalling: the shape of things to come. *Biochem. J., 339*, 481-8. ↗

Lal, H., Verma, SK., Foster, DM., Golden, HB., Reneau, JC., Watson, LE. et al. (2009). Integrins and proximal signaling mechanisms in cardiovascular disease. *Front. Biosci., 14*, 2307-34. ↗

#### Editions

| | | |
|---|---|---|
| 2008-03-11 | Edited | Garapati, P V. |
| 2008-05-07 | Authored | Geiger, B., Horwitz, AR. |
| 2008-05-07 | Reviewed | Humphries, MJ., Yamada, KM., Hynes, R. |
| 2012-08-08 | Authored | Jupe, S. |
| 2013-08-13 | Edited | Jupe, S. |
| 2013-08-13 | Reviewed | Ricard-Blum, S. |

#### 19 submitted entities found in this pathway, mapping to 21 Reactome entities

| Input | UniProt Id | col1 |
|---|---|---|
| COL6A3 | A8TX70, P12111 | 11 |
| COL6A1 | P12109 | 17 |
| COL6A2 | P12110 | 2 |
| COL4A1 | P02462, Q14055 | 3 |
| COL4A2 | P08572 | 4 |
| LUM | P51884 | 22 |
| COL1A1 | P02452 | 39 |
| COMP | P49747 | 4 |
| COL1A2 | P08123 | 40 |
| COL5A1 | P20908 | 5 |
| COL5A2 | P05997 | 1 |
| COL3A1 | P02461 | 12 |
| FN1 | P02751 | 31 |
| FBN1 | P35555 | 5 |
| AGRN | O00468 | 3 |
| TNC | P24821 | 5 |
| HSPG2 | P98160 | 2 |
| CD44 | P16070 | 3 |
| THBS1 | P07996 | 33 |

#### Invadopodia formation ↗

**Location:** Extracellular matrix organization

**Stable identifier:** R-HSA-8941237

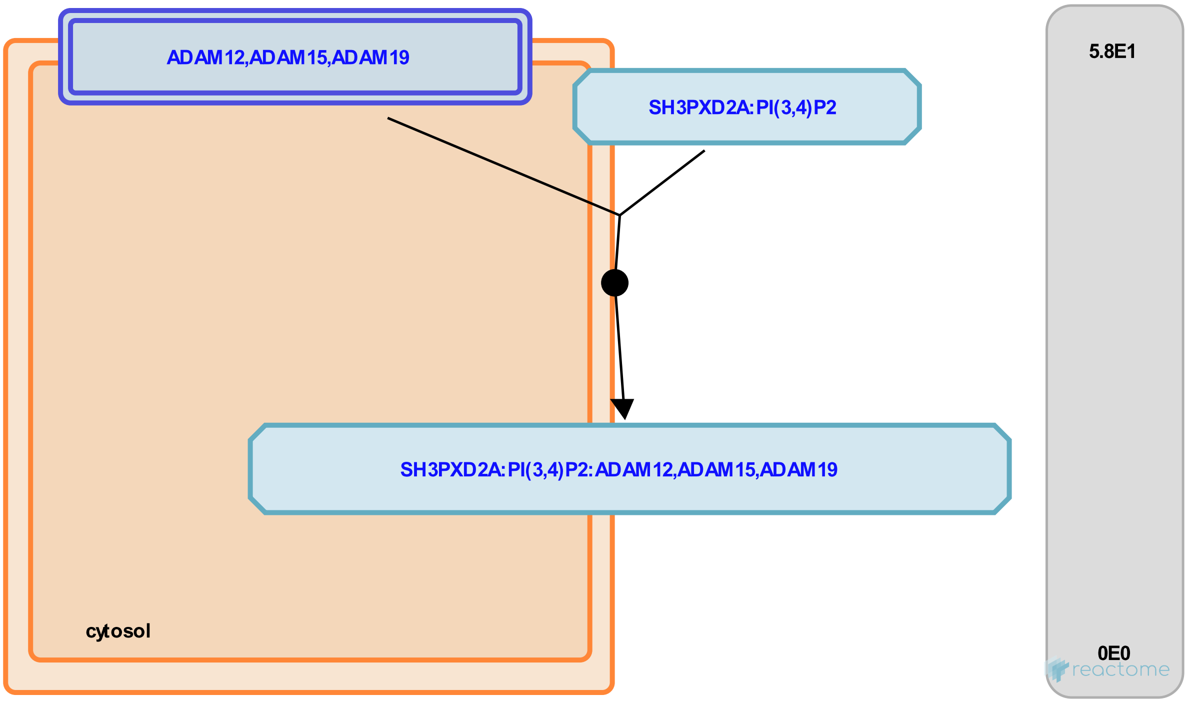


Podosomes and invadopodia are actin-based dynamic protrusions of the plasma membrane of metazoan cells that represent sites of attachment to and degradation of the extracellular matrix (Linder & Kopp 2005, Murphy & Courtneidge 2011). They are characteristically composed of an actin-rich core surrounded by adhesion and scaffolding proteins. Current convention is to use the term podosome for the structures found in normal cells (such as monocytic cells, endothelial cells and smooth muscle cells) and in Src-transformed fibroblasts, and invadopodium for the structures found in cancer cells. The maturation process for podosomes and invadopodia involves the recruitment and activation of multiple pericellular proteases, which facilitates ECM degradation (Artym et al. 2006).

#### Literature references

Gimona, M., Buccione, R., Courtneidge, SA., Linder, S. (2008). Assembly and biological role of podosomes and invadopodia. *Curr. Opin. Cell Biol.*, *20*, 235-41. ↗

#### Editions

| | | |
|---|---|---|
| 2016-09-29 | Authored | Jupe, S. |
| 2017-01-23 | Reviewed | Moreau, V. |
| 2017-02-28 | Edited | Jupe, S. |

### Table of Contents


Introduction 1

Analysis properties 2

Extracellular matrix organization 3

- Collagen formation 6
- Fibronectin matrix formation 8
- Elastic fibre formation 10
- Laminin interactions 13
- Non-integrin membrane-ECM interactions 15
- ECM proteoglycans 17
- Degradation of the extracellular matrix 19
- Integrin cell surface interactions 21
- Invadopodia formation 23

Table of Contents 24
