## Supplementary Data 1 for "Human SHED-derived extracellular cues activate a specialized neuroprotective and regenerative program in developing retinal ganglion cells": Nervous system development.pdf

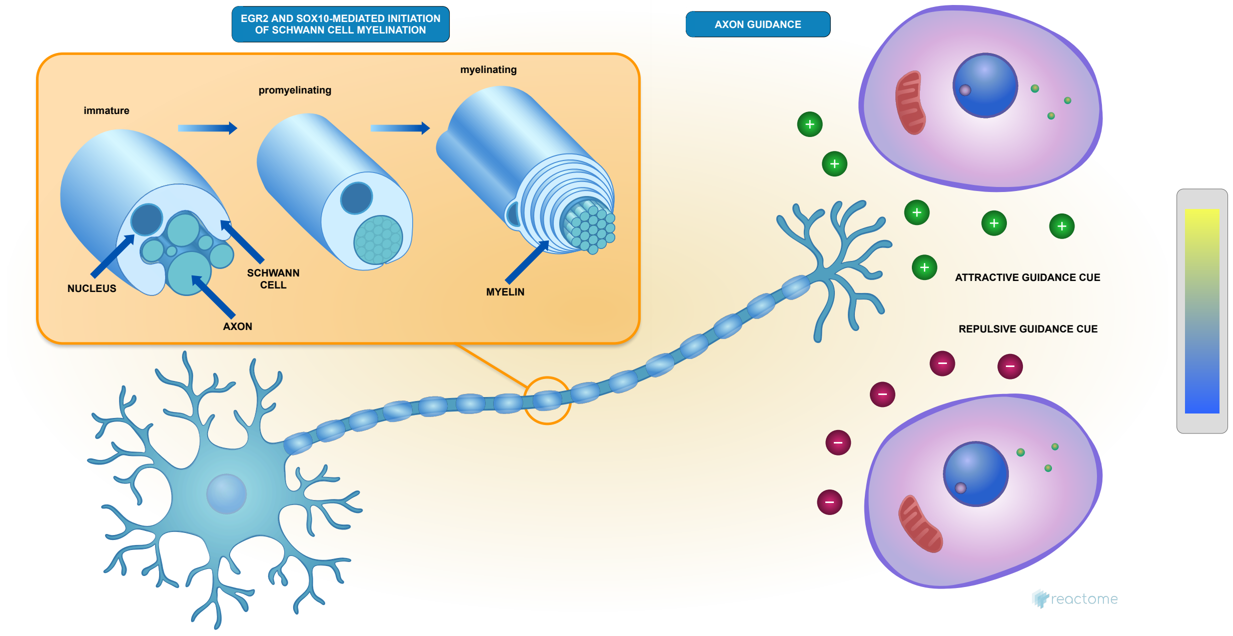


Aletta, J M., Garapati, P V., Maness, PF., Orlic-Milacic, M., Rothfels, K., Walmod, PS.

European Bioinformatics Institute, New York University Langone Medical Center, Ontario Institute for Cancer Research, Oregon Health and Science University.

#### Nervous system development ↗

**Stable identifier:** R-HSA-9675108

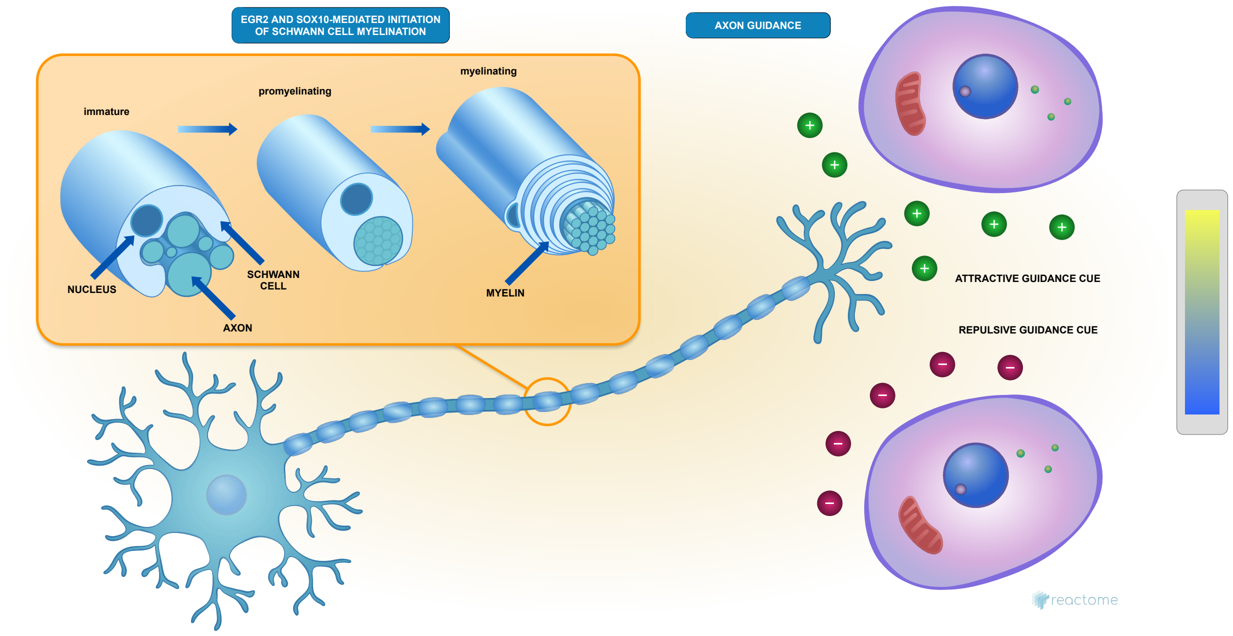


Neurogenesis is the process by which neural stem cells give rise to neurons, and occurs both during embryonic and perinatal development as well as in specific brain lineages during adult life (reviewed in Gotz and Huttner, 2005; Yao et al, 2016; Kriegstein and Alvarez-Buylla, 2009).

#### Literature references

Götz, M., Huttner, WB. (2005). The cell biology of neurogenesis. *Nat. Rev. Mol. Cell Biol., 6*, 777-88. ↗

Kriegstein, A., Alvarez-Buylla, A. (2009). The glial nature of embryonic and adult neural stem cells. *Annu. Rev. Neurosci., 32*, 149-84. ↗

Yao, B., Christian, KM., He, C., Jin, P., Ming, GL., Song, H. (2016). Epigenetic mechanisms in neurogenesis. *Nat. Rev. Neurosci., 17*, 537-49. ↗

#### Editions

| | | |
|---|---|---|
| 2020-01-23 | Reviewed | Orlic-Milacic, M. |
| 2020-01-31 | Authored, Edited | Rothfels, K. |

#### 27 submitted entities found in this pathway, mapping to 33 Reactome entities

| Input | UniProt Id | col1 |
|---|---|---|
| COL6A3 | A8TX70, P12111 | 11 |
| HSPA8 | P11142 | 12 |
| COL6A1 | P12109 | 17 |
| COL6A2 | P12110 | 2 |
| COL4A1 | P02462 | 3 |
| MMP2 | P08253 | 26 |
| LAMB1 | P07942 | 2 |
| COL4A2 | P08572 | 4 |
| GPC1 | P35052 | 9 |
| ACTB | P60709, P63261 | 54 |
| TUBB4B | P04350, P68371 | 3 |
| COL5A1 | P20908 | 5 |

| Input | UniProt Id | col1 |
|---|---|---|
| COL5A2 | P05997 | 1 |
| COL3A1 | P02461 | 12 |
| CFL1 | P23528 | 20 |
| MSN | P26038 | 4 |
| HSP90AB1 | P08238 | 10 |
| AGRN | O00468 | 3 |
| PFN1 | P07737 | 17 |
| MYH9 | P35579 | 4 |
| RPS23 | P62266 | 8 |
| TUBA1A | Q71U36 | 14 |
| RPL35 | P42766 | 10 |
| UBB | P0CG47, P62979, P62987 | 33 |
| RPL31 | P62899 | 21 |
| NRP2 | O60462, Q99435 | 2 |
| TLN1 | Q9Y490 | 8 |

### Axon guidance ↗

**Location:** Nervous system development

**Stable identifier:** R-HSA-422475

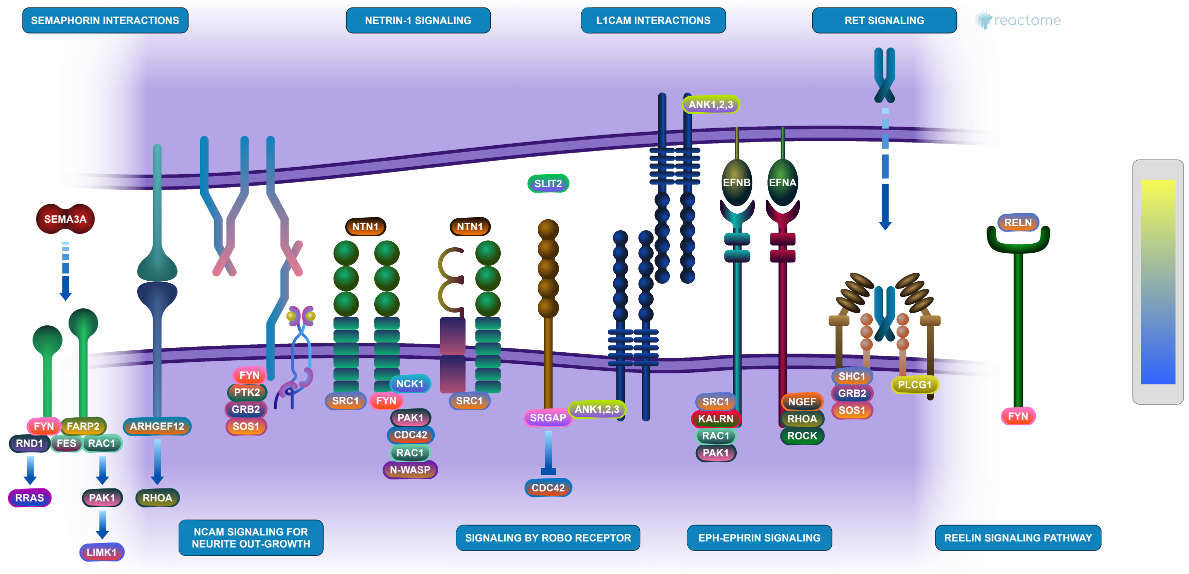


Axon guidance / axon pathfinding is the process by which neurons send out axons to reach the correct targets. Growing axons have a highly motile structure at the growing tip called the growth cone, which senses the guidance cues in the environment through guidance cue receptors and responds by undergoing cytoskeletal changes that determine the direction of axon growth.

#### EGR2 and SOX10-mediated initiation of Schwann cell myelination ↗

**Location:** Nervous system development

**Stable identifier:** R-HSA-9619665

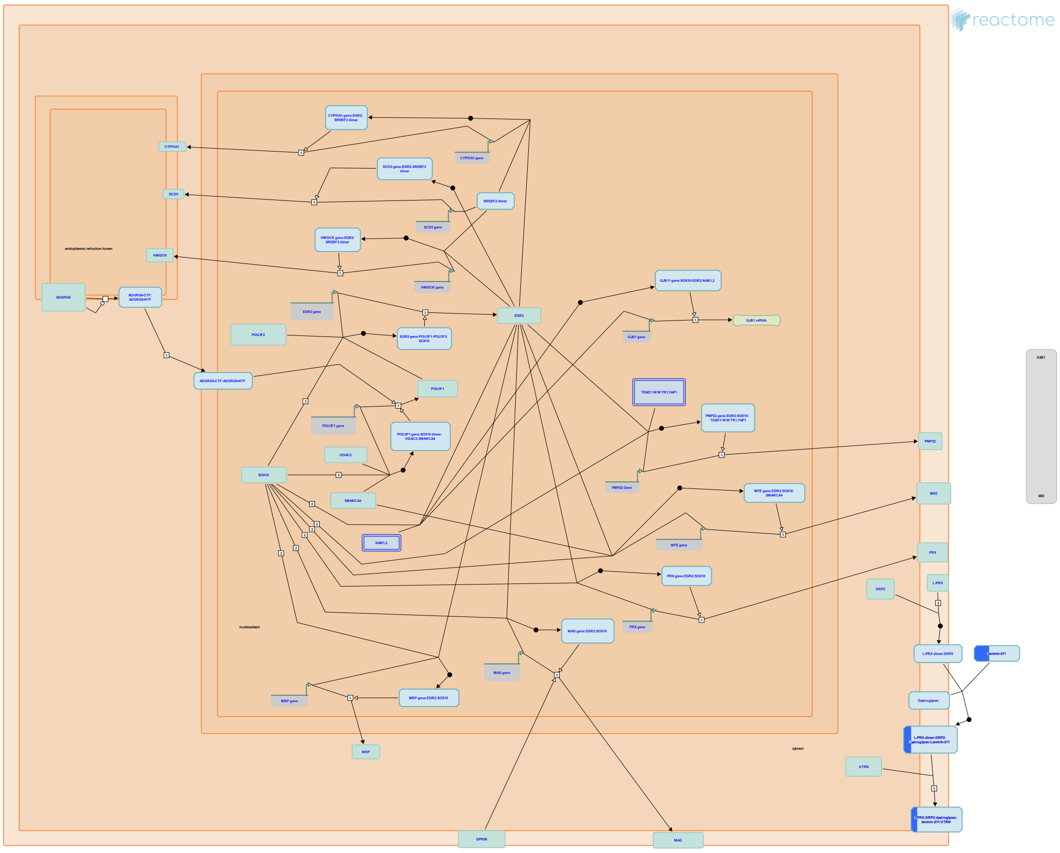

Schwann cells are glial cells of the peripheral nervous system that ensheath the peripheral nerves within a compacted lipid-rich myelin structure that is required for optimal transduction of nerve signals in motor and sensory nerves. Schwann cells develop from the neural crest in a differentiation process driven by factors derived from the Schwann cell itself, from the adjacent neuron or from the extracellular matrix (reviewed in Jessen and Mirsky, 2005). Upon peripheral nerve injury, mature Schwann cells can form repair cells that allow peripheral nerve regeneration through myelin phagocytosis and remyelination of the peripheral nerve. This process in some ways recapitulates the maturation of immature Schwann cells during development (reviewed in Jessen and Mirsky, 2016). Mature, fully myelinated Schwann cells exhibit longitudinal and radial polarization. The axon-distal abaxonal membrane interacts with elements of the basal lamina through integrins and lamins and in this way resembles the basolateral domain of polarized epithelial cells. In contrast, the axon-proximal adaxonal membrane resembles the apical domain of an epithelial cell, and is enriched with adhesion molecules and receptors that mediate interaction with ligands from the axon (reviewed in Salzer, 2015).
Schwann cells express a number of Schwann-cell specific proteins, including components of the myelin sheath such as myelin basic protein (MBP) and myelin protein zero (MPZ). In addition, Schwann cells have high lipid content relative to other membranes, and are enriched in galactosphingolipids, cholesterol and saturated long chain fatty acids (reviewed in Garbay et al, 2000). This protein and lipid profile is driven by a Schwann cell myelination transcriptional program controlled by master regulators SOX10, POU3F1 and EGR2, among others (reviewed in Svaren and Meijer, 2008; Stolt and Wegner, 2016).

#### Literature references


Jessen, KR., Mirsky, R. (2005). The origin and development of glial cells in peripheral nerves. *Nat. Rev. Neurosci., 6*, 671-82. ↗

Jessen, KR., Mirsky, R. (2016). The repair Schwann cell and its function in regenerating nerves. *J. Physiol. (Lond.), 594*, 3521-31. ↗

Salzer, JL. (2015). Schwann cell myelination. *Cold Spring Harb Perspect Biol, 7*, a020529. ↗

Garbay, B., Heape, AM., Sargueil, F., Cassagne, C. (2000). Myelin synthesis in the peripheral nervous system. *Prog. Neurobiol., 61*, 267-304. ↗

Svaren, J., Meijer, D. (2008). The molecular machinery of myelin gene transcription in Schwann cells. *Glia, 56*, 1541-51. ↗

#### Editions

| | | |
|---|---|---|
| 2019-08-16 | Authored | Rothfels, K. |
| 2020-01-17 | Reviewed | Aletta, J M. |
| 2020-02-24 | Edited | Rothfels, K. |

#### 1 submitted entities found in this pathway, mapping to 1 Reactome entities

| Input | UniProt Id | col1 |
|---|---|---|
| LAMB1 | P07942 | 2 |

### Table of Contents


Introduction 1

Analysis properties 2

Nervous system development 3

Axon guidance 5

EGR2 and SOX10-mediated initiation of Schwann cell myelination 7

Table of Contents 9
