## Supplementary Data 1 for "Human SHED-derived extracellular cues activate a specialized neuroprotective and regenerative program in developing retinal ganglion cells": Reactome full report.pdf

### Pathway Analysis Report

This report contains the pathway analysis results for the submitted sample ''. Analysis was performed against Reactome version 96 on 03/06/2026. The web link to these results is:

https://reactome.org/PathwayBrowser/#/ANALYSIS=MjAyNjA2MDMxNDU1NDNfOTEzOA%3D%3D

Please keep in mind that analysis results are temporarily stored on our server. The storage period depends on usage of the service but is at least 7 days. As a result, please note that this URL is only valid for a limited time period and it might have expired.

### Table of Contents

1. Introduction
2. Properties
3. Genome-wide overview
4. Most significant pathways
5. Pathways details
6. Identifiers found
7. Identifiers not found

### 1. Introduction

Reactome is a curated database of pathways and reactions in human biology. Reactions can be considered as pathway 'steps'. Reactome defines a 'reaction' as any event in biology that changes the state of a biological molecule. Binding, activation, translocation, degradation and classical biochemical events involving a catalyst are all reactions. Information in the database is authored by expert biologists, entered and maintained by Reactome's team of curators and editorial staff. Reactome content frequently cross-references other resources e.g. NCBI, Ensembl, UniProt, KEGG (Gene and Compound), ChEBI, PubMed and GO. Orthologous reactions inferred from annotation for Homo sapiens are available for 14 non-human species including mouse, rat, chicken, puffer fish, worm, fly and yeast. Pathways are represented by simple diagrams following an SBGN-like format.

Reactome's annotated data describe reactions possible if all annotated proteins and small molecules were present and active simultaneously in a cell. By overlaying an experimental dataset on these annotations, a user can perform a pathway over-representation analysis. By overlaying quantitative expression data or time series, a user can visualize the extent of change in affected pathways and its progression. A binomial test is used to calculate the probability shown for each result, and the p-values are corrected for the multiple testing (Benjamini–Hochberg procedure) that arises from evaluating the submitted list of identifiers against every pathway.

To learn more about our Pathway Analysis, please have a look at our relevant publications:


Fabregat A, Sidiropoulos K, Garapati P, Gillespie M, Hausmann K, Haw R, … D'Eustachio P (2016). The reactome pathway knowledgebase. Nucleic Acids Research, 44(D1), D481–D487. https://doi.org/10.1093/nar/gkv1351.

Fabregat A, Sidiropoulos K, Viteri G, Forner O, Marin-Garcia P, Arnau V, … Hermjakob H (2017). Reactome pathway analysis: a high-performance in-memory approach. BMC Bioinformatics, 18.

### 2. Properties

- This is an **expression** analysis: The numbers are used to produce a scaled coloured overlay over Reactome pathway diagrams, as a means to visualize relative expression levels. Note that the numeric values do not have to be expression data, for instance by using gene association scores the same analysis can be used to visualize genotyping results.
- 175 out of 192 identifiers in the sample were found in Reactome, where 1054 pathways were hit by at least one of them.
- All non-human identifiers have been converted to their human equivalent.
- This report is filtered to show only results for species 'Homo sapiens' and resource 'all resources'.
- The unique ID for this analysis (token) is MjAyNjA2MDMxNDU1NDNfOTEzOA%3D%3D. This ID is valid for at least 7 days in Reactome's server. Use it to access Reactome services with your data.

### 3. Genome-wide overview

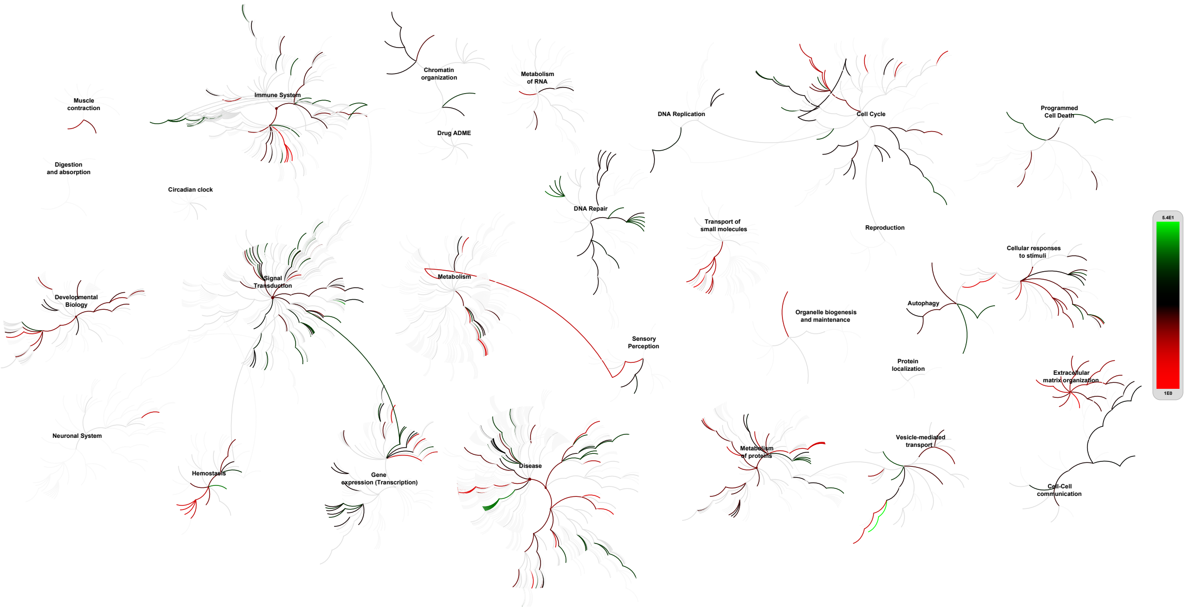


reactome

This figure shows a genome-wide overview of the results of your pathway analysis. Reactome pathways are arranged in a hierarchy. The center of each of the circular "bursts" is the root of one top-level pathway, for example "DNA Repair". Each step away from the center represents the next level lower in the pathway hierarchy. The color code denotes over-representation of that pathway in your input dataset. Light grey signifies pathways which are not significantly over-represented.

### 4. Most significant pathways

The following table shows the 25 most relevant pathways sorted by p-value.

| Pathway name | Entities | | | | Reactions | |
|---|---|---|---|---|---|---|
| | found | ratio | p-value | FDR* | found | ratio |
| Regulation of Insulin-like Growth Factor (IGF) transport and uptake by Insulin-like Growth Factor Binding Proteins (IGFBPs) | 37 / 127 | 0.008 | 1.11e-16 | 1.41e-14 | 14 / 14 | 8.75e-04 |
| Post-translational protein phosphorylation | 30 / 109 | 0.007 | 1.11e-16 | 1.41e-14 | 1 / 1 | 6.25e-05 |
| ECM proteoglycans | 24 / 79 | 0.005 | 1.11e-16 | 1.41e-14 | 19 / 23 | 0.001 |
| Degradation of the extracellular matrix | 31 / 148 | 0.009 | 1.11e-16 | 1.41e-14 | 74 / 105 | 0.007 |
| Extracellular matrix organization | 55 / 351 | 0.022 | 1.11e-16 | 1.41e-14 | 211 / 330 | 0.021 |
| Platelet degranulation | 32 / 142 | 0.009 | 1.11e-16 | 1.41e-14 | 7 / 11 | 6.87e-04 |
| Response to elevated platelet cytosolic Ca2+ | 32 / 149 | 0.009 | 1.11e-16 | 1.41e-14 | 7 / 14 | 8.75e-04 |
| Integrin cell surface interactions | 21 / 86 | 0.005 | 1.11e-16 | 1.41e-14 | 23 / 55 | 0.003 |
| Platelet activation, signaling and aggregation | 36 / 294 | 0.018 | 1.11e-16 | 1.41e-14 | 43 / 119 | 0.007 |
| Assembly of collagen fibrils and other multimeric structures | 18 / 67 | 0.004 | 5.55e-16 | 6.33e-14 | 20 / 26 | 0.002 |
| Non-integrin membrane-ECM interactions | 19 / 83 | 0.005 | 1.33e-15 | 1.39e-13 | 14 / 33 | 0.002 |
| Hemostasis | 51 / 821 | 0.051 | 3.44e-15 | 3.27e-13 | 106 / 366 | 0.023 |
| Collagen formation | 20 / 104 | 0.006 | 6.00e-15 | 5.28e-13 | 50 / 77 | 0.005 |
| Collagen degradation | 17 / 69 | 0.004 | 1.34e-14 | 1.09e-12 | 22 / 34 | 0.002 |
| Collagen biosynthesis and modifying enzymes | 16 / 76 | 0.005 | 8.51e-13 | 6.47e-11 | 30 / 51 | 0.003 |
| Collagen chain trimerization | 13 / 44 | 0.003 | 1.94e-12 | 1.38e-10 | 8 / 28 | 0.002 |
| Amyloid fiber formation | 15 / 89 | 0.005 | 9.50e-11 | 6.37e-09 | 8 / 33 | 0.002 |
| Insulin-like Growth Factor-2 mRNA Binding Proteins (IGF2BPs/IMPs/VICKZs) bind RNA | 8 / 13 | 8.03e-04 | 1.28e-10 | 8.04e-09 | 3 / 3 | 1.87e-04 |
| Immune System | 93 / 2,810 | 0.174 | 1.63e-10 | 9.77e-09 | 291 / 1,842 | 0.115 |
| Signaling by PDGF | 13 / 70 | 0.004 | 5.43e-10 | 3.10e-08 | 2 / 31 | 0.002 |
| Cytokine Signaling in Immune system | 50 / 1,106 | 0.068 | 6.20e-10 | 3.35e-08 | 51 / 806 | 0.05 |
| Developmental Cell Lineages | 23 / 283 | 0.017 | 1.70e-09 | 8.82e-08 | 14 / 22 | 0.001 |
| Axon guidance | 33 / 566 | 0.035 | 1.98e-09 | 9.68e-08 | 32 / 295 | 0.018 |
| Syndecan interactions | 9 / 29 | 0.002 | 3.34e-09 | 1.57e-07 | 5 / 15 | 9.37e-04 |

| Pathway name | Entities | | | | Reactions | |
|---|---|---|---|---|---|---|
| | found | ratio | p-value | FDR* | found | ratio |
| Fibronectin matrix formation | 8 / 21 | 0.001 | 5.25e-09 | 2.36e-07 | 4 / 4 | 2.50e-04 |

* False Discovery Rate

### 5. Pathways details

For every pathway of the most significant pathways, we present its diagram, as well as a short summary, its bibliography and the list of inputs found in it.

#### 1. Regulation of Insulin-like Growth Factor (IGF) transport and uptake by Insulin-like Growth Factor Binding Proteins (IGFBPs) (R-HSA-381426)

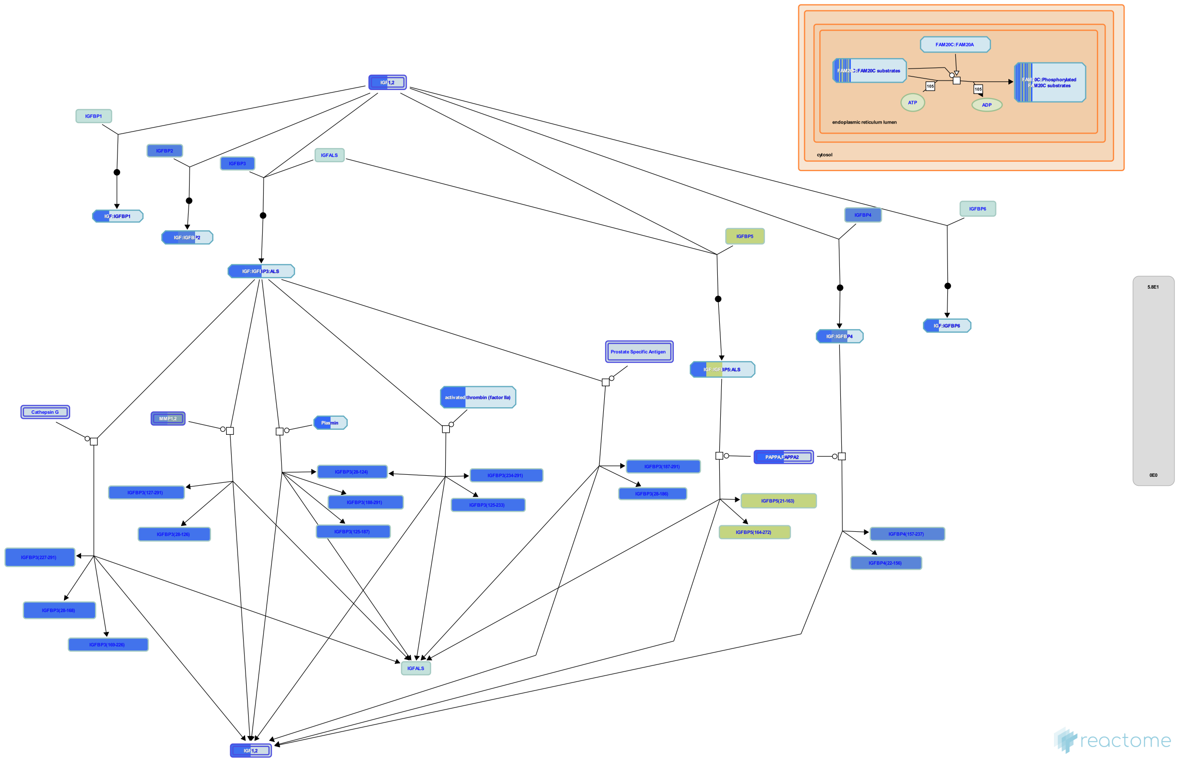

**Cellular compartments:** extracellular region.

The family of Insulin like Growth Factor Binding Proteins (IGFBPs) share 50% amino acid identity with conserved N terminal and C terminal regions responsible for binding Insulin like Growth Factors I and II (IGF I and IGF II). Most circulating IGFs are in complexes with IGFBPs, which are believed to increase the residence of IGFs in the body, modulate availability of IGFs to target receptors for IGFs, reduce insulin like effects of IGFs, and act as signaling molecules independently of IGFs.
About 75% of circulating IGFs are in 1500 220 KDa complexes with IGFBP3 and ALS. Such complexes are too large to pass the endothelial barrier. The remaining 20 25% of IGFs are bound to other IGFBPs in 40 50 KDa complexes. IGFs are released from IGF:IGFBP complexes by proteolysis of the IGFBP. IGFs become active after release, however IGFs may also have activity when still bound to some IGFBPs.
IGFBP1 is enriched in amniotic fluid and is produced in the liver under control of insulin (insulin suppresses production). IGFBP1 binding stimulates IGF function. It is unknown which if any protease degrades IGFBP1.
IGFBP2 is enriched in cerebrospinal fluid; its binding inhibits IGF function. IGFBP2 is not significantly degraded in circulation.
IGFB3, which binds most IGF in the body is enriched in follicular fluid and found in many other tissues. IGFBP 3 may be cleaved by plasmin, thrombin, Prostate specific Antigen (PSA, KLK3), Matrix Metalloprotease-1 (MMP1), and Matrix Metalloprotease-2 (MMP2). IGFBP3 also binds extracellular matrix and binding lowers its affinity for IGFs. IGFBP3 binding stimulates the effects of IGFs.
IGFBP4 acts to inhibit IGF function and is cleaved by Pregnancy associated Plasma Protein A (PAPPA) to release IGF.
IGFBP5 is enriched in bone matrix; its binding stimulates IGF function. IGFBP5 is cleaved by Pregnancy Associated Plasma Protein A2 (PAPPA2), ADAM9, complement C1s from smooth muscle, and thrombin. Only the cleavage site for PAPPA2 is known.
IGFBP6 is enriched in cerebrospinal fluid. It is unknown which if any protease degrades IGFBP6.


##### Edit history

| Date | Action | Author |
|---|---|---|
| 2008-11-20 | Edited | May B, Gopinathrao G |
| 2008-11-20 | Created | May B |
| 2008-12-02 | Reviewed | Matthews L, D'Eustachio P, Gillespie ME |

| Date | Action | Author |
|---|---|---|
| 2026-03-16 | Modified | Weiser JD |

#### 37 submitted entities found in this pathway, mapping to 37 Reactome entities

| Input | UniProt Id | col1 |
|---|---|---|
| NUCB1 | Q02818 | 10 |
| TIMP1 | P01033 | 25 |
| AHSG | P02765 | 8 |
| CCN1 | O00622 | 7 |
| MMP2 | P08253 | 26 |
| LAMB1 | P07942 | 2 |
| MMP1 | P03956 | 15 |
| QSOX1 | O00391 | 18 |
| APOE | P02649 | 5 |
| IGFBP7 | Q16270 | 41 |
| LGALS1 | P09382 | 38 |
| APOB | P04114 | 1 |
| IGF2 | P01344 | 5 |
| TF | P02787 | 54 |
| F2 | P00734 | 4 |
| PAPPA | Q13219 | 1 |
| FN1 | P02751 | 31 |
| SPARC | Q14515 | 42 |
| LTBP1 | Q14766 | 1 |
| SERPINC1 | P01008 | 7 |
| FBN1 | P35555 | 5 |
| ITIH2 | P19823 | 9 |
| ALB | P02768 | 12 |
| STC2 | O76061 | 5 |
| VCAN | P13611 | 1 |
| TNC | P24821 | 5 |
| PLG | P00747 | 2 |
| C4A | P0C0L4 | 2 |
| AFP | P02771 | 8 |
| CALU | O43852 | 11 |
| C3 | P01024 | 4 |
| IGFBP2 | P18065 | 11 |
| IGFBP3 | P17936 | 6 |
| IGFBP4 | P22692 | 13 |
| IGFBP5 | P24593 | 44 |
| GAS6 | Q14393 | 6 |
| FSTL1 | Q12841 | 51 |

#### 2. Post-translational protein phosphorylation (R-HSA-8957275)

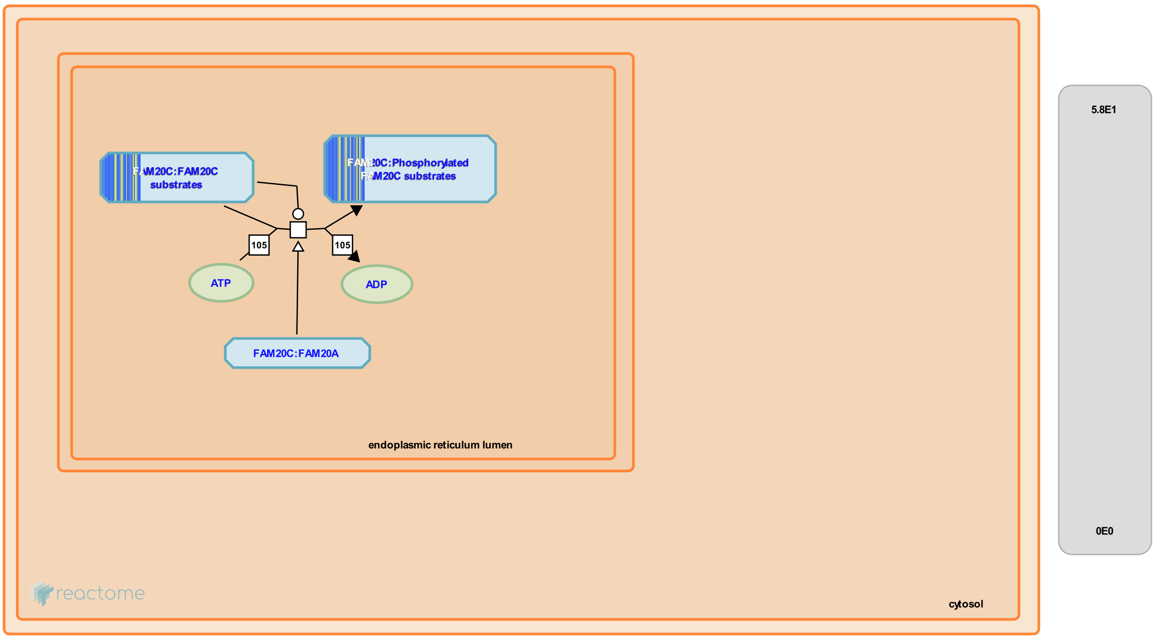


Secretory pathway kinases phosphorylate a diverse array of substrates involved in many physiological processes.


##### Edit history

| Date | Action | Author |
|---|---|---|
| 2016-12-08 | Authored | Jupe S |
| 2017-01-23 | Reviewed | Wiley SE |
| 2017-01-24 | Edited | Jupe S |
| 2017-01-24 | Created | Jupe S |

##### 30 submitted entities found in this pathway, mapping to 30 Reactome entities

| Input | UniProt Id | col1 |
|---|---|---|
| NUCB1 | Q02818 | 10 |
| TIMP1 | P01033 | 25 |
| AHSG | P02765 | 8 |
| CCN1 | O00622 | 7 |
| LAMB1 | P07942 | 2 |
| QSOX1 | O00391 | 18 |
| APOE | P02649 | 5 |
| IGFBP7 | Q16270 | 41 |
| LGALS1 | P09382 | 38 |
| APOB | P04114 | 1 |
| TF | P02787 | 54 |

| Input | UniProt Id | col1 |
|---|---|---|
| FN1 | P02751 | 31 |
| SPARC | Q14515 | 42 |
| LTBP1 | Q14766 | 1 |
| SERPINC1 | P01008 | 7 |
| FBN1 | P35555 | 5 |
| ITIH2 | P19823 | 9 |
| ALB | P02768 | 12 |
| STC2 | O76061 | 5 |
| VCAN | P13611 | 1 |
| TNC | P24821 | 5 |
| C4A | P0C0L4 | 2 |
| AFP | P02771 | 8 |
| CALU | O43852 | 11 |
| C3 | P01024 | 4 |
| IGFBP3 | P17936 | 6 |
| IGFBP4 | P22692 | 13 |
| IGFBP5 | P24593 | 44 |
| GAS6 | Q14393 | 6 |
| FSTL1 | Q12841 | 51 |

#### 3. ECM proteoglycans (R-HSA-3000178)

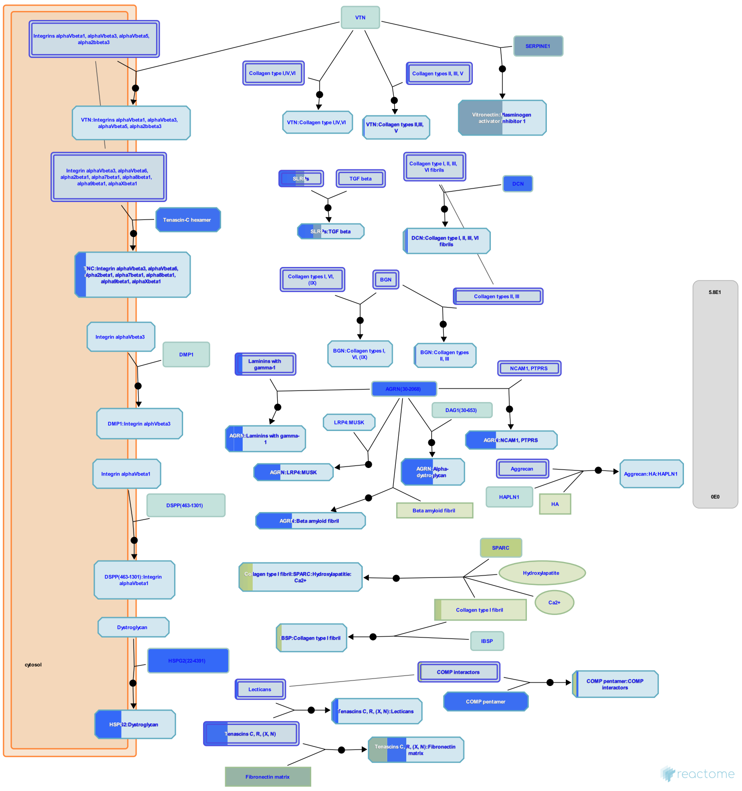

Proteoglycans are major components of the extracellular matrix. In cartilage the matrix constitutes more than 90% of tissue dry weight. Proteoglycans are proteins substituted with glycosaminoglycans (GAGs), linear polysaccharides consisting of a repeating disaccharide, generally of an acetylated amino sugar alternating

with a uronic acid. Most proteoglycans are located in the extracellular

space. Proteoglycans are highly diverse, both in terms of the core proteins and the subtypes of GAG chains, namely chondroitin sulfate (CS), keratan sulfate (KS), dermatan sulfate (DS) and heparan sulfate (HS). Hyaluronan is a non-sulfated GAG whose molecular weight runs into millions of Dalton; in articular cartilage, a single hyaluronan molecule can hold upto 100 aggrecan molecules and these aggregates are stabilized by a link protein.


##### Edit history

| Date | Action | Author |
|---|---|---|
| 2013-01-10 | Authored | Jupe S |

| Date | Action | Author |
|---|---|---|
| 2013-01-24 | Created | Jupe S |
| 2013-04-26 | Edited | Jupe S |
| 2013-05-21 | Reviewed | Venkatesan N |
| 2013-05-22 | Reviewed | Ricard-Blum S |
| 2026-03-16 | Modified | Weiser JD |

#### 22 submitted entities found in this pathway, mapping to 24 Reactome entities

| Input | UniProt Id | col1 |
|---|---|---|
| COL6A3 | A8TX70, P12111 | 11 |
| COL6A1 | P12109 | 17 |
| COL6A2 | P12110 | 2 |
| COL4A1 | P02462, Q14055 | 3 |
| LAMB1 | P07942 | 2 |
| COL4A2 | P08572 | 4 |
| LUM | P51884 | 22 |
| COL1A1 | P02452 | 39 |
| COMP | P49747 | 4 |
| COL1A2 | P08123 | 40 |
| COL5A1 | P20908 | 5 |
| COL5A2 | P05997 | 1 |
| COL3A1 | P02461 | 12 |
| FN1 | P02751 | 31 |
| SPARC | P09486 | 42 |
| DCN | P07585 | 4 |
| AGRN | O00468 | 3 |
| SERPINE1 | P05121 | 24 |
| VCAN | P13611 | 1 |
| FMOD | Q06828 | 5 |
| TNC | P24821 | 5 |
| HSPG2 | P98160 | 2 |

#### 4. Degradation of the extracellular matrix (R-HSA-1474228)

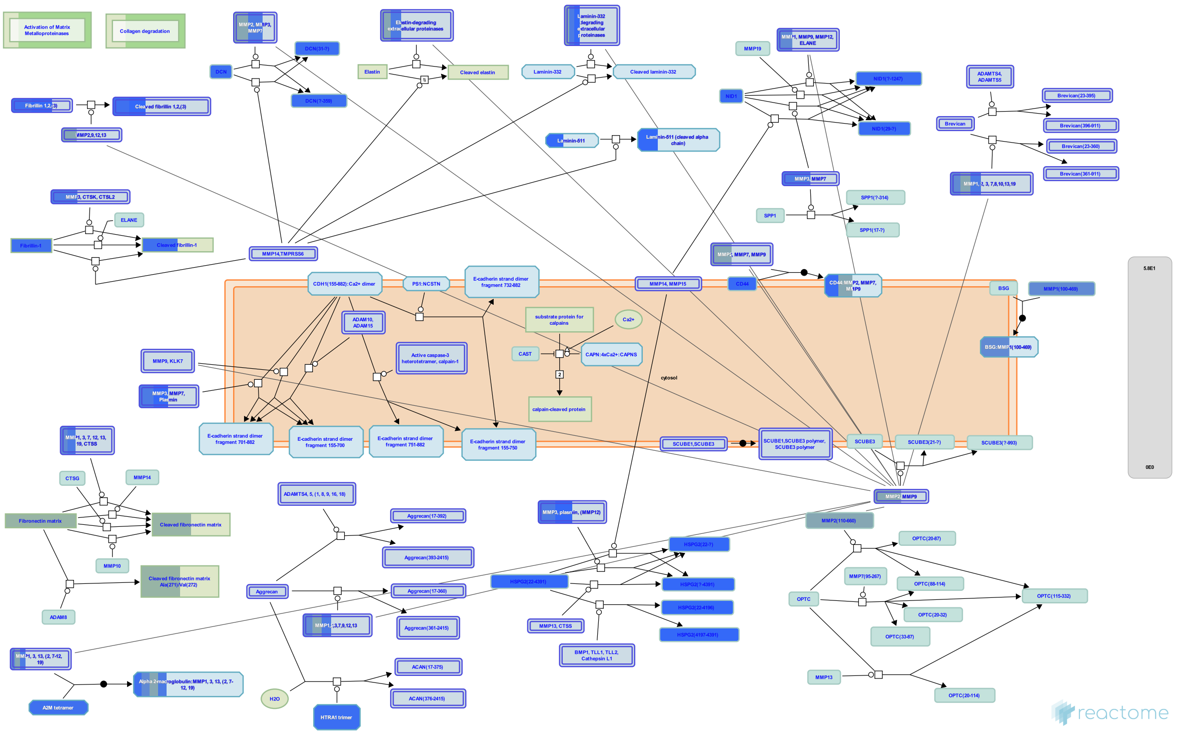


Matrix metalloproteinases (MMPs), previously referred to as matrixins because of their role in degradation of the extracellular matrix (ECM), are zinc and calcium dependent proteases belonging to the metzincin family. They contain a characteristic zinc-binding motif HEXXHXXGXXH (Stocker & Bode 1995) and a conserved Methionine which forms a Met-turn. Humans have 24 MMP genes giving rise to 23 MMP proteins, as MMP23 is encoded by two identical genes. All MMPs contain an N-terminal secretory signal peptide and a prodomain with a conserved PRCGXPD motif that in the inactive enzyme is localized with the catalytic site, the cysteine acting as a fourth unpaired ligand for the catalytic zinc atom. Activation involves delocalization of the domain containing this cysteine by a conformational change or proteolytic cleavage, a mechanism referred to as the cysteine-switch (Van Wart & Birkedal-Hansen 1990). Most MMPs are secreted but the membrane type MT-MMPs are membrane anchored and some MMPs may act on intracellular proteins. Various domains determine substrate specificity, cell localization and activation (Hadler-Olsen et al. 2011). MMPs are regulated by transcription, cellular location (most are not activated until secreted), activating proteinases that can be other MMPs, and by metalloproteinase inhibitors such as the tissue inhibitors of metalloproteinases (TIMPs). MMPs are best known for their role in the degradation and removal of ECM molecules. In addition, cleavage of the ECM and other cell surface molecules can release ECM-bound growth factors, and a number of non-ECM proteins are substrates of MMPs (Nagase et al. 2006). MMPs can be divided into subgroups based on domain structure and substrate specificity but it is clear that these are somewhat artificial, many MMPs belong to more than one functional group (Vise & Nagase 2003, Somerville et al. 2003).


##### Edit history

| Date | Action | Author |
|---|---|---|
| 2011-08-05 | Created | Jupe S |
| 2011-09-09 | Authored | Jupe S |
| 2012-02-21 | Edited | Jupe S |
| 2012-02-28 | Reviewed | D'Eustachio P |
| 2026-03-16 | Modified | Weiser JD |

#### 28 submitted entities found in this pathway, mapping to 31 Reactome entities

| Input | UniProt Id | col1 |
|---|---|---|
| CTSB | P07858 | 12 |
| TIMP2 | P16035 | 14 |
| MMP3 | P08254 | 9 |
| COL6A3 | A8TX70, P12111 | 11 |
| TIMP1 | P01033 | 25 |
| COL6A1 | P12109 | 17 |
| COL6A2 | P12110 | 2 |
| COL4A1 | P02462, Q14055 | 3 |
| MMP2 | P08253 | 26 |
| LAMB1 | P07942 | 2 |
| COL4A2 | P08572 | 4 |
| MMP1 | P03956 | 15 |
| A2M | P01023 | 5 |
| COL1A1 | P02452 | 39 |
| COL1A2 | P08123 | 40 |
| COL5A1 | P20908 | 5 |
| COL5A2 | P05997 | 1 |
| COL3A1 | P02461 | 12 |
| FN1 | P02751 | 31 |
| HTRA1 | Q92743 | 4 |
| DCN | P07585 | 4 |
| FBN1 | P35555 | 5 |
| FBN2 | P35556, Q75N90 | 2 |
| COL12A1 | Q99715 | 7 |
| PLG | P00747 | 2 |
| HSPG2 | P98160 | 2 |
| NID1 | P14543 | 3 |
| CD44 | P16070 | 3 |

5. **Extracellular matrix organization** (R-HSA-1474244)
