## Supplementary Figure Legends for "Human SHED-derived extracellular cues activate a specialized neuroprotective and regenerative program in developing retinal ganglion cells"

**Supplementary Figure 1**. **SHED and DPSC exhibit preserved mesenchymal stem cell properties.**  Representative phase-contrast micrographs of DPSC and SHED at passage 2 (a) reveal the characteristic spindle-shaped fibroblast-like morphology associated with mesenchymal stromal cells, whereas DPSC at passage 5 (b) display increased cellular heterogeneity, enlarged cell bodies and flattened morphology consistent with progressive replicative aging. In contrast, SHED maintain a highly homogeneous fibroblastoid morphology at both passage 2 (c) and passage 5 (d), exhibiting limited evidence of senescence-associated changes. Following lineage-specific induction, osteogenic differentiation was confirmed by Alizarin Red staining (e), adipogenic differentiation by Oil Red O staining (f), and chondrogenic differentiation by Alcian Blue staining (g). Together, these findings confirm the preservation of mesenchymal stem cell identity and multilineage differentiation potential in both populations while highlighting the superior proliferative stability and morphological consistency of SHED cultures.

**Supplementary Figure 2. Schematic representation of the experimental paradigms used to distinguish developmental axonogenesis from regenerative axon growth in embryonic chick retinal explants.** At embryonic day 5 (E5) (a), retinal progenitor cells actively generate retinal ganglion cells (RGC) that simultaneously undergo neuronal differentiation, axonogenesis and physiological developmental cell death. EdU incorporation *in vitro* enables identification of newly generated neurons. At embryonic day 13 (E13) (b), retinal neurogenesis is largely complete and retinal neurons are postmitotic. Explant preparation results in axotomy of mature retinal ganglion cells, creating a model of regenerative neurite growth. This experimental design enables independent assessment of SHED-mediated effects on developmental neuronal growth and regenerative responses within the same biological system.

**Supplementary Figure 3. Additional representative examples of SHED-induced neurite outgrowth in embryonic retinal explants.** High-magnification images of E5 retinal explants (a) demonstrate increased neurite density, enhanced axonal elongation and more extensive neuronal networks following SHED treatment compared with control and HaCaT conditions. Similarly, representative E13 retinal explants (b) illustrate robust regenerative axonal growth from axotomized retinal ganglion cells exposed to SHED-derived signals. Scale bar = 50 μm.

**Supplementary Figure 4. Representative images stained for TUNEL (green), and DAPI (blue) provide additional evidence supporting the neuroprotective effects of SHED treatment in E13.** Across all experimental conditions SHED-treated retinal explants consistently exhibit reduced numbers of apoptotic nuclei together with compared with control and HaCaT-treated cultures. These findings confirm the reproducibility of the neuroprotective phenotype observed in both developmental and injury-associated paradigms.

**Supplementary Figure 5. Reactome pathway enrichment analysis of proteins identified within the SHED secretome.** The global Reactome network (a) reveals extensive enrichment of pathways associated with developmental biology, nervous system development, axon guidance, neuronal differentiation and regenerative responses. Functional clustering highlights coordinated biological programs associated with (b) developmental biology/nervous system development specifically axonogenesis and neuronal growth, whereas additional clusters identify pathways involved in insulin-like growth factor (IGF) signaling modules, are also significantly enriched (c). Growth factor-associated pathways, including extracellular matrix organization, cell adhesion and integrin-mediated signaling (c). (d)Node size reflects pathway significance and color intensity corresponds to enrichment magnitude (protein sequence coverage).

**Supplementary Figure 6. Gene Ontology analyses further validate the regenerative profile of the SHED secretome. g:Profiler e**nriched biological processes (a) include nervous system development, neurite morphogenesis, extracellular matrix organization and tissue remodeling. All significantly enriched molecular functions are associated with extracellular matrix interactions, receptor binding, growth factor modulation and cell adhesion, whereas enriched cellular componentshighlight extracellular vesicles, extracellular matrix structures and secreted protein complexes as listed in (b). Collectively, these analyses support the existence of a coordinated extracellular signaling network capable of promoting neuronal growth, survival and regenerative remodeling.

**Supplementary Figure 7. Inhibition of THBS1 signaling alters neurite architecture and axonal integrity.** Representative TUJ1-positive neurons cultured alone or in the presence of SHED together with control IgG, anti-THBS1 antibody,(**a**) demonstrate marked alterations in neuronal morphology following disruption of THBS1–α2δ1 signaling. Quantitative analyses reveal **partial** reductions in the proportion of neurons bearing clearly identifiable axons after THBS1–α2δ1 inhibition conditions (b) In parallel, inhibition of THBS1 signaling increases the occurrence of neuritic swellings and dystrophic neurites (c), consistent with impaired axonal maintenance and regenerative failure. These findings support a central role for THBS1-dependent signaling in mediating SHED-induced axon growth.

**Supplementary Figure 8. Functional characterization of the damage-responsive SHED secretome.** Gene Ontology enrichment analysis of the 24 proteins significantly upregulated following retinal co-culture (a) identifies biological processes associated with regulation of programmed cell death, neuronal survival and cellular adaptation to injury (b). Gene Ontology enrichment analysis of the 30 proteins listed here significantly downregulated following retinal co-culture (b).

**Supplementary Table 1.** **Comprehensive proteomic analysis of the SHED secretome.** The table summarizes the proteins identified in the SHED secretome together with their corresponding statistical parameters. Results are shown after protein annotation and identification against (a) the *UniProtKB Homo sapiens* database and (b) the *UniProtKB Gallus gallus* database. Statistical values and protein identification metrics obtained from the mass spectrometry analysis are provided for each detected protein.

**Supplementary Table 2.** **Reactome pathway enrichment analysis of the SHED secretome.** The table includes all significantly enriched Reactome pathways identified from the proteins detected in the SHED secretome, together with the corresponding enrichment metrics (including statistical significance and pathway enrichment parameters). For each pathway, the associated proteins contributing to the enrichment are listed.

**Supplementary Data 1. Complete Reactome pathway enrichment report generated from the SHED secretome dataset.** The file contains the full Reactome analysis output, including all enriched pathways and associated enrichment statistics. Detailed results are provided for pathways of particular relevance to this study, including Axon Guidance, Nervous System Development, and Extracellular Matrix Organization, together with the proteins contributing to each enriched pathway.
